# Interpretable genotype-to-phenotype classifiers with performance guarantees

**DOI:** 10.1101/388348

**Authors:** Alexandre Drouin, Gaël Letarte, Frédéric Raymond, Mario Marchand, Jacques Corbeil, François Laviolette

## Abstract

Understanding the relationship between the genome of a cell and its phenotype is a central problem in precision medicine. Nonetheless, genotype-to-phenotype prediction comes with great challenges for machine learning algorithms that limit their use in this setting. The high dimensionality of the data tends to hinder generalization and challenges the scalability of most learning algorithms. Additionally, most algorithms produce models that are complex and difficult to interpret. We alleviate these limitations by proposing strong performance guarantees, based on sample compression theory, for rule-based learning algorithms that produce highly interpretable models. We show that these guarantees can be leveraged to accelerate learning and improve model interpretability. Our approach is validated through an application to the genomic prediction of antimicrobial resistance, an important public health concern. Highly accurate models were obtained for 12 species and 56 antibiotics, and their interpretation revealed known resistance mechanisms, as well as some potentially new ones. An open-source disk-based implementation that is both memory and computationally efficient is provided with this work. The implementation is turnkey, requires no prior knowledge of machine learning, and is complemented by comprehensive tutorials.

## Introduction

The relationship between the genome of a cell and its phenotype is central to precision medicine. Specific mutations in the human genome are known to affect the metabolism of drugs and thus influence the response to treatments and the toxicity of common drugs like warfarin or azathioprine^1^. Similarly, mutated genes in bacteria lead to increased virulence or resistance to antimicrobial agents, which leads to an increased risk of morbidity^2^. Large-scale studies that aim to link genomic features to clinical outcomes are now common in both eukaryotes and prokaryotes.

The most common type of studies are genome-wide association studies (GWAS), which aim to identify all statistically significant genotype-to-phenotype associations^3, 4^. An alternative approach consists of using machine learning algorithms to build models that correlate genomic variations with phenotypes^5, 6^. This approach contrasts with GWAS in that the objective shifts from thoroughly understanding the phenotype, to accurately predicting it based on the occurrence of genomic variations.

Nevertheless, the accurate prediction of phenotypes is insufficient for many applications in biology, such as clinical diagnostics. Models must rely on a decision process that can be validated by domain experts and thus, algorithms that produce interpretable models are preferred^6, 7^. Rule-based classifiers are models that make predictions by answering a series of questions, such as “Is there a mutation at base pair 42 in this gene?” Such models are highly interpretable and the decision logic can be validated experimentally to confirm accuracy and potentially lead to the extraction of new biological knowledge.

In this study, two algorithms that learn rule-based models are explored: i) Classification and Regression Trees^8^(CART) and ii) Set Covering Machines^9^ (SCM). The former learns decision trees, which are hierarchical arrangements of rules and the latter learns conjunctions (logical-AND) and disjunctions (logical-OR), which are simple logical combinations of rules. Their accuracy and interpretability are demonstrated with an application to the prediction of antimicrobial resistance (AMR) in bacteria, a global public health concern of high significance. Several thousand bacterial genomes and their susceptibility to antimicrobial agents are publicly available^10^ and make for an ideal study set. The use of machine learning to predict AMR phenotypes has previously been investigated using two approaches: 1) considering only known resistance genes and mutations^11–14^, 2) considering whole genomes with no prior knowledge of resistance mechanisms^15–20^. The work described hereafter relies on the latter approach.

The contributions of this study are multidisciplinary. From the biological perspective, 107 highly accurate models of antimicrobial resistance, covering 12 human pathogens and 56 antibiotics, are obtained using each of the previously described algorithms. These models identify known resistance mechanisms, as well as some potentially new ones. From the machine learning perspective, the study establishes that rule-based models are well suited for genotype-to-phenotype prediction and demonstrates their accuracy in comparison to state-of-the-art models. A mathematical analysis based on sample compression theory^21, 22^ provides strong statistical guarantees on the accuracy of the obtained models, which are essential if the models are to be applied in diagnosis and prognosis^23^. These guarantees are used to prune the models, increasing their interpretability, while dramatically accelerating computing times. Finally, an efficient, disk-based implementation of both algorithms, which efficiently scales to increasingly large genomic datasets, is proposed and made publicly available. Importantly, the proposed approach is not limited to AMR prediction and could be applied to a plethora of phenotypes.

## Results

### Overview of the data

The data used in this study were extracted from the Pathosystems Resource Integration Center (PATRIC) database, one of the most comprehensive public databases of bacterial genomes and antimicrobial resistance metadata^10, 24^. The protocol used to acquire the data is detailed in *Methods* and the amount of data extracted is shown, per species, in Fig. 1a. In total, 107 binary classification datasets were extracted, each consisting of discriminating isolates that are resistant or susceptible to an antimicrobial agent, based on their genome, in a given species (e.g., kanamycin resistance in *M. tuberculosis*). The genomes in each dataset were represented by the presence and absence of every *k*-mer (i.e., sequence of *k* nucleotides) of length 31 that occurred in the data^17^ (see *Methods*). As illustrated in Fig. 1b, species with high genomic plasticity, such as *Klebsiella pneumoniae*, are associated with greater *k*-mer counts, whereas species with low diversity, such as *Mycobacterium tuberculosis*, are associated with lower *k*-mer counts.

**Figure 1.**
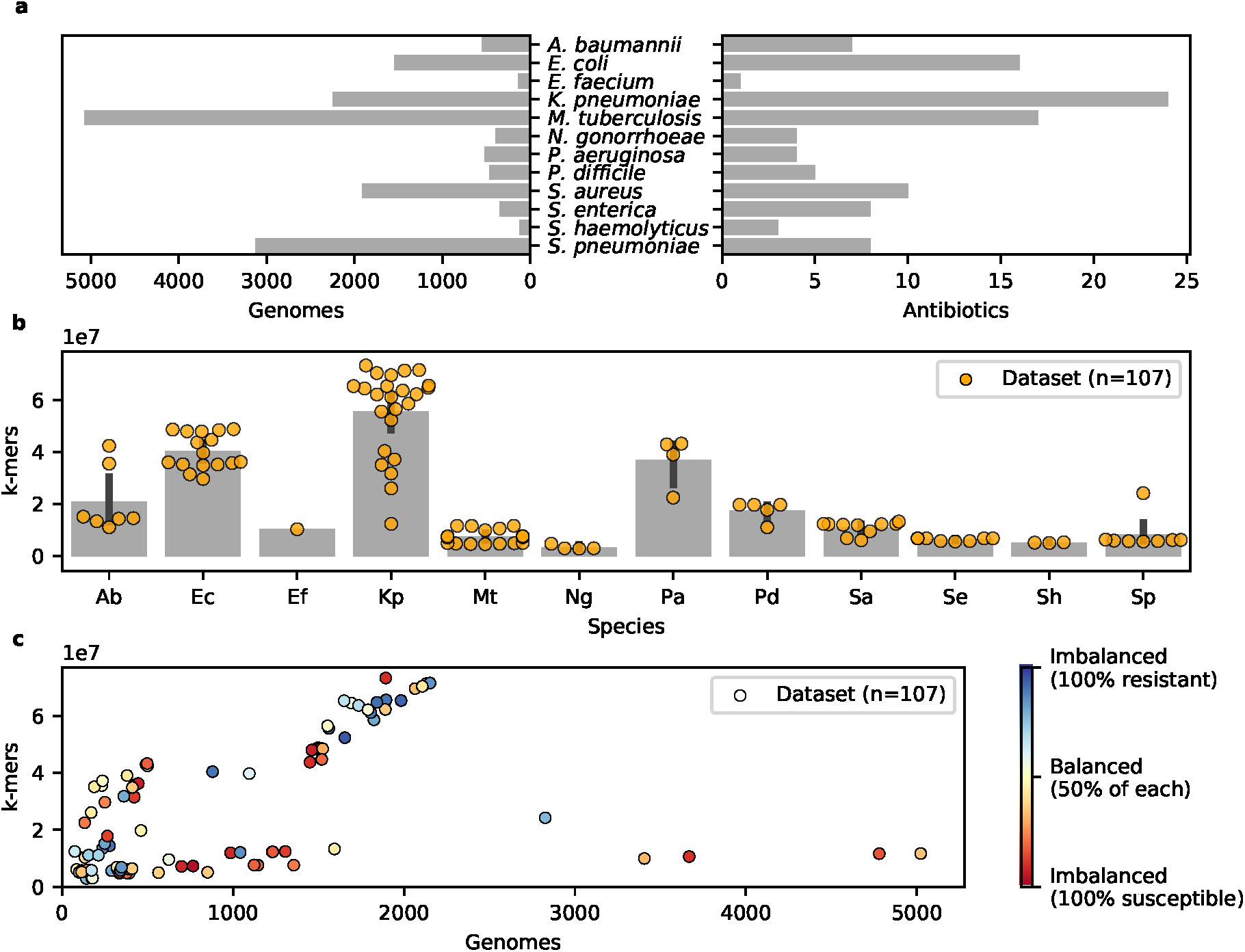
Summary of the PATRIC data. a) Number of genomes and antibiotics for which data was extracted, shown by species. b) Number of *k*-mers in each dataset (dots), shown by species. Low *k*-mer counts reflect populations with homogeneous genomes, whereas the converse indicates high genomic diversity. c) For each dataset (dots) the number of examples (genomes) and features (*k*-mers) is shown, along with a measure of class imbalance. Clearly, some datasets contain more examples of one of the classes (resistant or susceptible) and each dataset shows a strong discrepancy between the number of examples and features. Together, these conditions make for challenging learning tasks.

The resulting datasets pose significant challenges for machine learning algorithms, which are reflected in Fig. 1c. First, the *sample size* (number of genomes) is extremely small compared to the size of the *feature space* (number of *k*-mers). This setting, known as *fat data*^25, 26^, is very common in genotype-to-phenotype studies and generally leads to models that overfit the data^23^, i.e., a situation in which the model fits the training data perfectly, but performs poorly on unseen data. Second, many datasets show strong class imbalance (i.e., one class is more abundant than the other), due to the fact that more resistant or susceptible genomes were available. This can lead to models that are accurate for the most abundant class and perform poorly on the least abundant one. Hence, this study will show that accurate models, that are also interpretable, can be learned despite these challenges and provide a theoretical justification of these results.

### Rule-based models based on performance guarantees

This study proposes performance guarantees for the CART and SCM algorithms under the form of sample compression bounds (see *Methods*). Such bounds are of theoretical interest, as they explain how accurate models can be learned despite the challenging setting. Nonetheless, the relevance of these bounds goes beyond simple theoretical justification, since they can be used to improve *model selection*. This essential phase of the learning process consists of setting the hyperparameters of the algorithms (e.g., the maximum depth of a decision tree) and pruning the resulting models to reduce their complexity. The typical approach to model selection relies on cross-validation, a computationally intensive procedure that involves training the algorithms several times on subsets of the data. In this study, the proposed sample compression bounds are used to dramatically accelerate model selection, while allowing all the data to be used for training (see *Methods*). In the following experiments, the proposed *bound-based* algorithms are referred to as CART_*b*_ and SCM_*b*_, whereas the typical *cross-validation-based* algorithms are referred to as CART_*cv*_ and SCM_*cv*_.

### Genotype-to-phenotype prediction with rule-based models

The CART_*b*_ and SCM_*b*_ algorithms were trained on the aforementioned datasets to obtain rule-based predictors of antimicrobial resistance. Each dataset was randomly partitioned into disjoint training and validation sets, using 80% and 20% of the data respectively. The training sets were used to construct predictive models and the validation sets were used to assess their ability to generalize to unseen genomes. This procedure was repeated ten times, using different random partitions, in order to obtain accurate estimates of generalization performance despite the small number of examples in some datasets.

#### The models are highly accurate

Fig. 2 illustrates the accuracies of the models, which correspond to the proportion of correct AMR phenotype assignments in the validation data, for CART_*b*_ and SCM_*b*_ across all datasets. Both algorithms perform comparably (*p* = 0.603 – according to a Wilcoxon signed-rank test), which is reflected in the highly similar distribution of model accuracies over the datasets. Moreover, both rule-based algorithms learn highly accurate models, despite the challenging nature of the datasets. In fact, 95% of the models have accuracies greater than 80%, 75% greater than 90%, and 45% (almost half) greater than 95%. This suggests that the rule-based models produced by CART_*b*_ and SCM_*b*_ are well-suited for genotype-to-phenotype prediction. The ability of CART_*b*_ and SCM_*b*_ to learn accurate models in this setting is characteristic of their strong resistance to overfitting. This counterintuitive result is further supported by Supplementary Fig. S1, which shows that the accuracy of the models does not depend on the number of *k*-mers in the data and that accurate models can be learned regardless of the sample size. The theoretical performance guarantees, presented in *Methods*, provide a mathematical justification of these empirical results. Detailed results for each dataset are available in Supplementary Table S1, where several metrics are reported. A detailed comparison of the accuracy of these algorithms is kept for the next section, where they are compared to other state-of-the-art algorithms.

**Figure 2.**
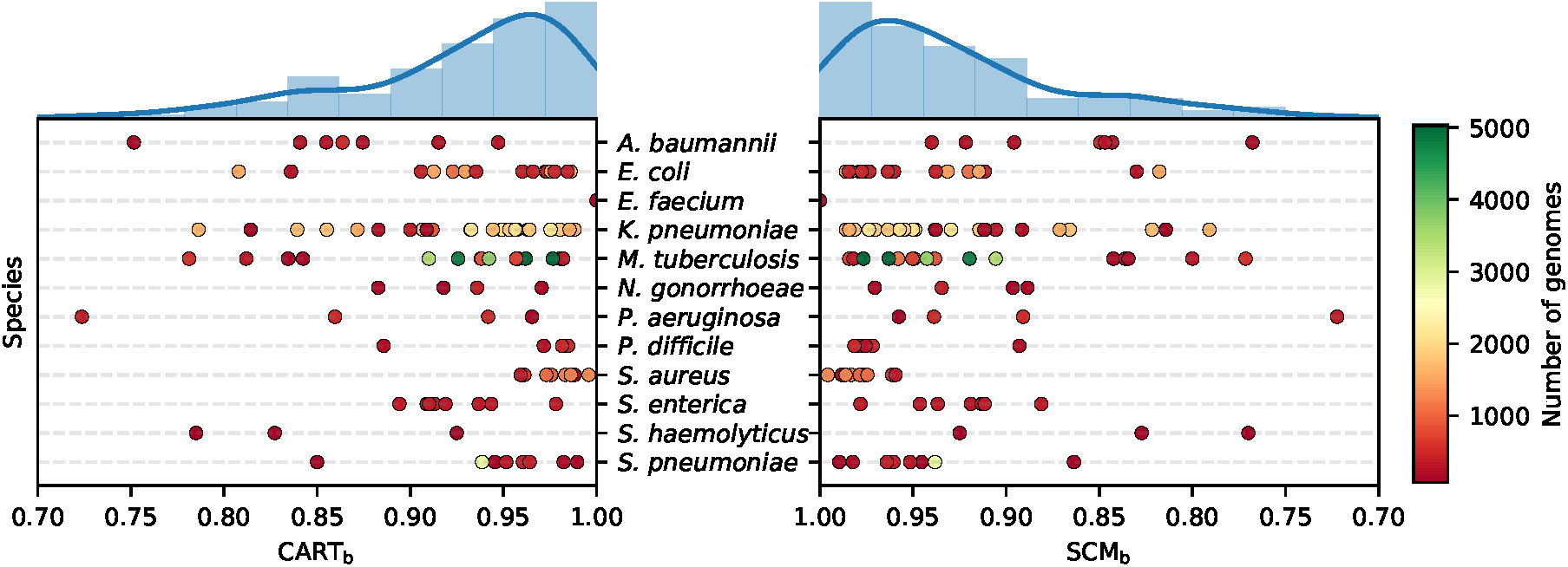
Accuracies of CART_*b*_ and SCM_*b*_ on the validation data of each dataset, grouped by species. The datasets are shown as dots and colored according to the number of genomes that they contain. The distribution of accuracies for each method is also shown (top). All models have an accuracy greater than 70%.

#### The models are highly interpretable

Fig. 3 illustrates rule-based models learned for two datasets: kanamycin resistance in *M. tuberculosis* and meropenem resistance in *K. pneumoniae*. Three properties are illustrated for each rule in the models: 1) the locus at which the corresponding *k*-mer can be found, 2) a measure of rule importance, and 3) the number of equivalent rules. The first is the region of the genome in which the *k*-mer is located and was determined using the Basic Local Alignment Search Tool^27^ (BLAST). The second quantifies the contribution of a rule to the model’s predictions. The greater a rule’s importance, the more examples of the training data it serves to discriminate. The rule importances are measured according to Breiman et al. (1984)^8^ for CART_*b*_ and Drouin et al. (2016)^17^ for SCM_*b*_, and were normalized to sum to one. The third results from *k*-mers that are equally predictive of the phenotype. For instance, *k*-mers located on the same gene may always be present or absent simultaneously, resulting in several rules that are equally predictive for the model. Equivalent rules were shown to be indicative of the nature of genomic variations^17^. A small number of equivalent rules, with *k*-mers overlapping a certain position of the genome, suggests a point mutation, whereas a large number, targeting multiple contiguous *k*-mers, indicates large scale genomic rearrangements, such as gene insertions and deletions. Finally, for visualization purposes, the SCM_*b*_ and CART_*b*_ models were selected so that the SCM_*b*_ model was a subset of the CART_*b*_ one. This illustrates the ability of CART_*b*_ to learn models that are slightly more complex than those of SCM_*b*_, extending them beyond simple conjunctions and disjunctions. While the SCM_*b*_ models are not always subsets of the CART_*b*_ models, it was observed that 81% of the models have at least their most important rule in common with their counterpart.

**Figure 3.**
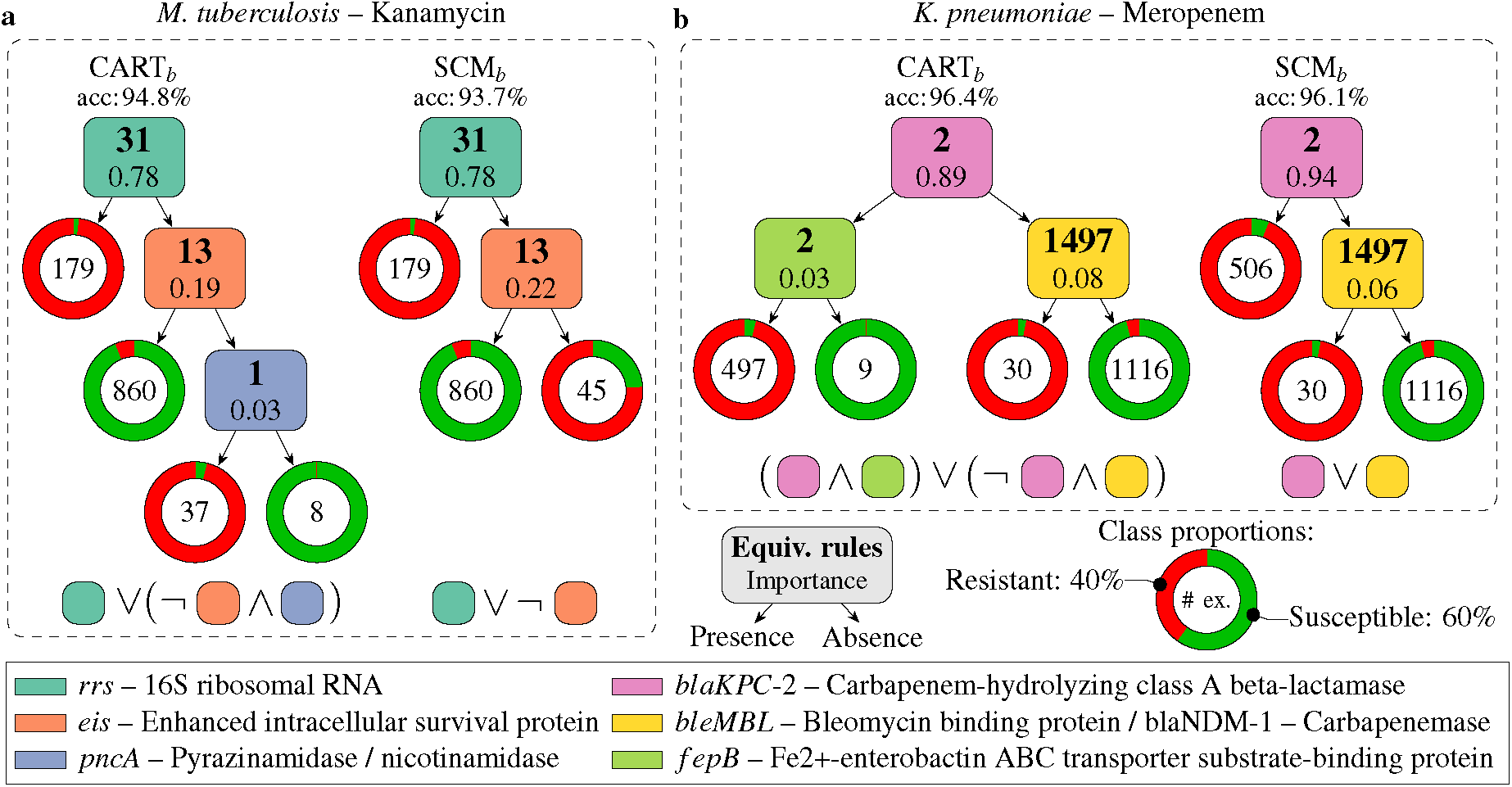
Rule-based genotype-to-phenotype classifiers. Each classifier is shown as a hierarchical arrangement of rules (boxes) and leaves (rings). Predictions are made by placing a genome at the root and branching left or right based on the outcome of the rules until a leaf is reached. A leaf predicts the most abundant class among the training examples that were guided into it. The number of such examples is shown in its center and the ring is colored according to the distribution of their phenotypes (classes). Each rule detects the presence/absence of a *k*-mer and was colored according to the genomic locus at which it was found. Additionally, each box contains a measure of rule importance and the number of rules that were found to be equivalent during training. Finally, the logical expression leading to the prediction of the *resistant* phenotype is shown, under each model, to contrast the structure of models learned by each algorithm. In these expressions, the rules are shown as squares with colors that match those of the models and the negation of a rule is used to indicate the “absence” of the *k*-mer. In both cases, the CART models are disjunctions of conjunctions (ORs of ANDs) and the SCM models are disjunctions (ORs).

The first set of models, shown in Fig. 3a, are tasked with predicting kanamycin resistance in *Mycobacterium tuberculosis*. Kanamycin is an aminoglycoside antibiotic and a key second-line drug in the fight against multi-drug resistant infections^28^. This drug acts by binding to the 16S rRNA (*rrs* gene) in the 30S ribosomal subunit to inhibit protein synthesis^29^. Mutations in *rrs* are known to confer kanamycin resistance^29, 30^. Consistently, both models predict resistance in the presence of the A1401G mutation in *rrs*, a known resistance determinant^29, 31, 32^. The nature of this mutation was determined by observing that the 31 equivalent rules target *k*-mers that overlap at a single base-pair location on *rrs* (i.e., 1401) and detect the presence of a guanine at this locus. The second most important rule in both models, and its equivalent rules, target *k*-mers in the promoter region of the *eis* gene, which harbors several resistance-conferring mutations^33^. This rule predicts the susceptible phenotype in the presence of the wild-type sequence. In its absence, the SCM_*b*_ model predicts resistance, which efficiently captures the occurrence of several known resistance-conferring mutations using a single rule. The CART_*b*_ model also uses this rule, but adds an additional requirement for resistance: the presence of a *k*-mer in the *pncA* gene, which is associated with resistance to pyrazinamide, a first-line anti-tuberculosis drug^28^. This is consistent with the fact that kanamycin is a second-line drug, used in the presence of resistance to first-line treatments. Based on the leaves of the models, it can be observed that this additional requirement allows a better separation of resistant and susceptible isolates, resulting in a more accurate model. Of note, SCM_*b*_ could not have added such a rule to its model, since the resulting model is more complex than a single conjunction or disjunction.

The second set of models, shown in Fig. 3b, are tasked with predicting resistance to meropenem, a broad spectrum carbapenem antibiotic, in *Klebsiella pneumoniae*. The first rule in both models targets the *blaKPC-2* gene, a carbapenem hydrolyzing beta-lactamase that is known to confer resistance to carbapenem antibiotics^34^. The small number of equivalent rules likely results from the models targeting specificities of *blaKPC-2* to discriminate it from its variants, such as *blaKPC-1, blaKPC-3*. The SCM_*b*_ model predicts resistance in the presence of a *k*-mer in this gene. However, the CART_*b*_ model adds another requirement: the presence of a *k*-mer in the *fepB* gene, which encodes a periplasmic protein that is essential for virulence^35^. Based on the leaves of the models, it can be observed that this additional requirement allows to correctly classify nine susceptible isolates that were misclassified by SCM_*b*_. Furthermore, both models capture another resistance mechanism, represented by 1497 equivalent rules. Interestingly, the *k*-mers targeted by these rules completely cover the *bleMBL* and *blaNDM-1* genes, which are generally present simultaneously and part of the same operon^36, 37^. *bleMBL* encodes a protein responsible for resistance to bleomycin, an anti-cancer drug, and is not causal of meropenem resistance^36, 37^. *blaNDM-1* encodes a carbapenemase and is a known resistance determinant for meropenem^38^. Nevertheless, since both genes generally occur simultaneously, they were found to be equally good predictors of meropenem resistance. Finally, notice that, once again, the CART_*b*_ model is slightly more accurate than that of SCM_*b*_ and that its structure goes beyond a simple conjunction or disjunction.

In summary, the CART_*b*_ and SCM_*b*_ can learn highly accurate genotype-to-phenotype models from which relevant biological knowledge can be extracted. Several confirmed antibiotic resistance mechanisms were identified using only genome sequences categorized according to their phenotypes. This demonstrates the great potential of rule-based classifiers in predicting and understanding the genomic foundations of phenotypes that are currently misunderstood.

### Comparison to state-of-the-art classifiers

The CART_*b*_ and SCM_*b*_ algorithms were compared to state-of-the-art classifiers on a benchmark of several datasets, described in Table 1. The benchmark includes one dataset per species selected in order to have the largest possible sample size with minimal class imbalance (see *Methods*). While we only report results for the benchmark datasets, results for the remaining datasets are available in Supplementary Table S1. For each dataset, the protocol described in the previous section (ten repetitions) was used. The CART_*b*_ and SCM_*b*_ algorithms were compared to five other learning algorithms, including *L*1-regularized logistic regression^39^(*L*1-Logistic), *L*2-regularized logistic regression^40^(*L*2-Logistic), Polynomial Kernel Support Vector Machines^41^ (Poly-SVM), and two baseline methods: Naive Bayes^42^, and the simple predictor that returns the most abundant class in the training data (Majority). These choices are motivated in *Methods*. An extended benchmark with a comparison to additional methods, which were not included for conciseness, is available in Supplementary Table S2.

**Table 1.** Overview of the benchmark datasets.

| Species | Antibiotic | Examples | Resistant | Susceptible | *k*-mers |
| --- | --- | --- | --- | --- | --- |
| *A. baumannii* | Imipenem | 499 | 325 | 174 | 42 406 238 |
| *E. faecium* | Vancomycin | 134 | 51 | 83 | 10 341 091 |
| *E. coli* | Amoxicillin/Clavulanic acid | 1 524 | 464 | 1 060 | 48 456 086 |
| *K. pneumoniae* | Gentamicin | 2 107 | 906 | 1 201 | 70 347 931 |
| *M. tuberculosis* | Isoniazid | 5 022 | 1 719 | 3 303 | 11 688 883 |
| *N. gonorrhoeae* | Azithromycin | 392 | 214 | 178 | 4 766 702 |
| *P. difficile* | Moxifloxacin | 462 | 188 | 274 | 19 753 432 |
| *P. aeruginosa* | Levofloxacin | 491 | 201 | 290 | 42 961 897 |
| *S. enterica* | Chloramphenicol | 347 | 251 | 96 | 6 864 155 |
| *S. aureus* | Methicillin | 1 593 | 707 | 886 | 13 289 281 |
| *S. haemolyticus* | Ciprofloxacin | 120 | 74 | 46 | 5 341 646 |
| *S. pneumoniae* | Chloramphenicol | 409 | 149 | 260 | 6 380 123 |

The results of the benchmark are shown in Table 2, where the accuracy and complexity of the models learned by each algorithm are compared. Once again, the results indicate that the CART_*b*_ and SCM_*b*_ algorithms perform comparably in terms of accuracy (*p* = 0.600). However, CART_*b*_ learns models that rely on slightly more *k*-mers than those of SCM_*b*_ for 5 out of 12 (5/12) datasets (*p* = 0.046). In addition, it can be observed that both algorithms compare favorably to the other algorithms in the benchmark in terms of accuracy and model complexity. The accuracy of CART_*b*_ is better or equal to that of *L*1-logistic on 6/12 datasets (*p* = 0.879), *L*2-logistic on 10/12 datasets (*p* = 0.075), Poly-SVM on 9/12 datasets (*p* = 0.028), and Naive Bayes on all datasets (*p* = 0.002). Similarly, the accuracy of SCM_*b*_ is better or equal to that of *L*1-logistic on 7/12 datasets (*p* = 0.879), *L*2-logistic on 10/12 datasets (*p* = 0.075), Poly-SVM on 9/12 datasets (*p* = 0.050), and Naive Bayes on all datasets (*p* = 0.002). Regarding model complexity, both rule-based algorithms learn models that rely on strictly less *k*-mers than the other algorithms for all datasets, including *L*1-logistic, which is well-known to yield sparse models^39^. Hence, the rule-based models show state-of-the-art accuracy, while relying on significantly less genomic variants, making them easier to interpret, validate, and translate to clinical settings.

**Table 2.**
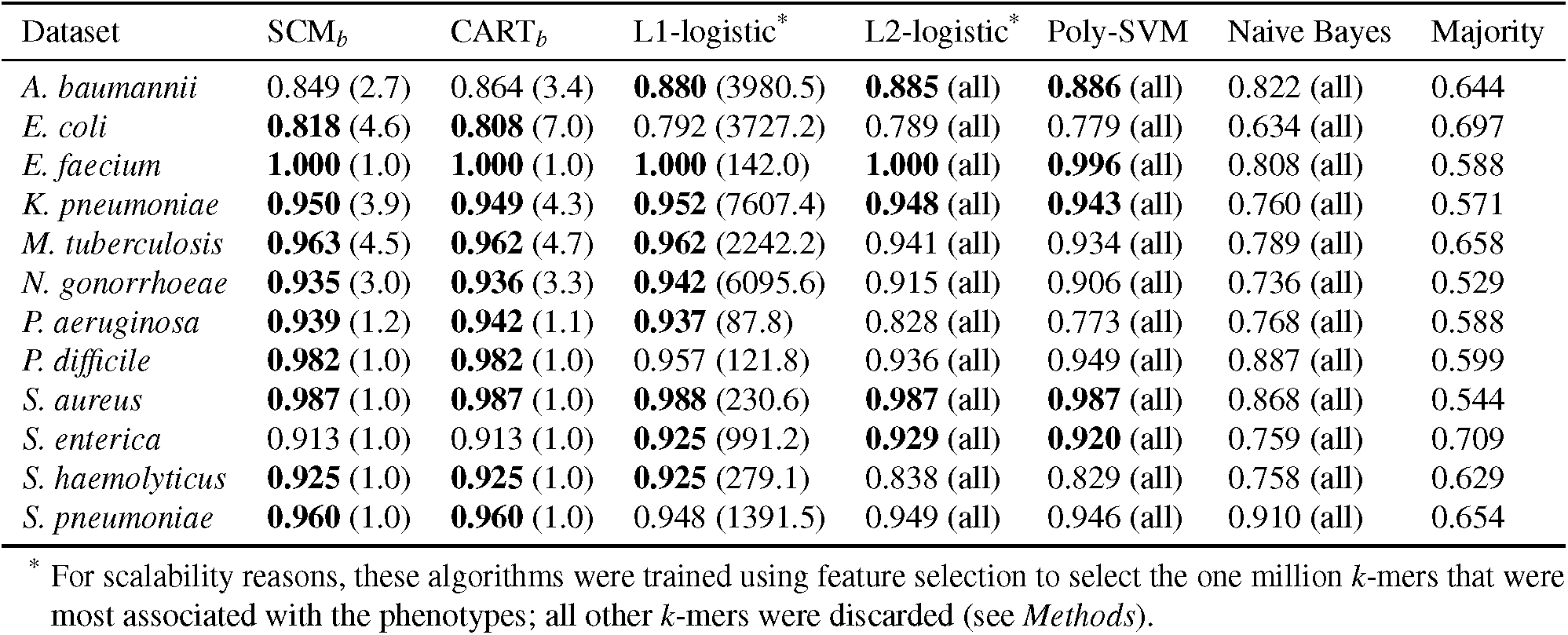
Comparison to state-of-the-art classifiers in terms of accuracy and model complexity. For each dataset the accuracy is shown, along with the number of *k*-mers used by the model (in parentheses). Results are shown for Set Covering Machines (SCM), Classification trees (CART), Logistic regression with L1 and L2 regularization and *χ*^2^ feature selection (L1-logistic, L2-logistic), Polynomial kernel Support Vector Machines (Poly-SVM), Naive Bayes, and a baseline predictor that predicts the most abundant class in the data (Majority). Accuracies within 1% of the maximum value are shown in bold. Results are averaged over ten repetitions of the experiment.

| Dataset | SCM<sub>b</sub> | CART<sub>b</sub> | L1-logistic* | L2-logistic* | Poly-SVM | Naive Bayes | Majority |
| --- | --- | --- | --- | --- | --- | --- | --- |
| *A. baumannii* | 0.849 (2.7) | 0.864 (3.4) | **0.880** (3980.5) | **0.885** (all) | **0.886** (all) | 0.822 (all) | 0.644 |
| *E. coli* | **0.818** (4.6) | **0.808** (7.0) | 0.792 (3727.2) | 0.789 (all) | 0.779 (all) | 0.634 (all) | 0.697 |
| *E. faecium* | **1.000** (1.0) | **1.000** (1.0) | **1.000** (142.0) | **1.000** (all) | **0.996** (all) | 0.808 (all) | 0.588 |
| *K. pneumoniae* | **0.950** (3.9) | **0.949** (4.3) | **0.952** (7607.4) | **0.948** (all) | **0.943** (all) | 0.760 (all) | 0.571 |
| *M. tuberculosis* | **0.963** (4.5) | **0.962** (4.7) | **0.962** (2242.2) | 0.941 (all) | 0.934 (all) | 0.789 (all) | 0.658 |
| *N. gonorrhoeae* | **0.935** (3.0) | **0.936** (3.3) | **0.942** (6095.6) | 0.915 (all) | 0.906 (all) | 0.736 (all) | 0.529 |
| *P. aeruginosa* | **0.939** (1.2) | **0.942** (1.1) | **0.937** (87.8) | 0.828 (all) | 0.773 (all) | 0.768 (all) | 0.588 |
| *P. difficile* | **0.982** (1.0) | **0.982** (1.0) | 0.957 (121.8) | 0.936 (all) | 0.949 (all) | 0.887 (all) | 0.599 |
| *S. aureus* | **0.987** (1.0) | **0.987** (1.0) | **0.988** (230.6) | **0.987** (all) | **0.987** (all) | 0.868 (all) | 0.544 |
| *S. enterica* | 0.913 (1.0) | 0.913 (1.0) | **0.925** (991.2) | **0.929** (all) | **0.920** (all) | 0.759 (all) | 0.709 |
| *S. haemolyticus* | **0.925** (1.0) | **0.925** (1.0) | **0.925** (279.1) | 0.838 (all) | 0.829 (all) | 0.758 (all) | 0.629 |
| *S. pneumoniae* | **0.960** (1.0) | **0.960** (1.0) | 0.948 (1391.5) | 0.949 (all) | 0.946 (all) | 0.910 (all) | 0.654 |

\* For scalability reasons, these algorithms were trained using feature selection to select the one million $k$ -mers that were most associated with the phenotypes; all other $k$ -mers were discarded (see *Methods*).

### Sample compression bounds for model selection

In the previous experiments, model selection was performed using the proposed sample compression bounds in lieu of cross-validation (see *Methods*). As illustrated in Fig. 4, this approach, referred to as *bound selection*, is much faster than ten-fold cross-validation. In fact, for every set of hyperparameter values, ten-fold cross-validation requires ten trainings of the learning algorithm, while bound selection only requires one. However, the success of bound selection is highly dependent on the quality (tightness) of the generalization bounds used. A bound that is insufficiently tight results in inaccurate estimates of the generalization performance of models and, consequently, inaccurate model selection. To assess the accuracy of the proposed bounds in the context of model selection, models trained using cross-validation (CART_*cv*_, SCM_*cv*_) and bound selection (CART_*b*_, SCM_*b*_) were compared in terms of accuracy and model complexity. The same protocol as in the previous sections was used. The results, shown in Table 3, indicate that bound selection leads to models that are comparably accurate, but considerably more concise. This reflects a fundamental principle that is embedded in the mathematical expression of the bounds: models that are both simple and accurate are less subject to overfitting than their more complex counterparts (see *Methods*). In complement, Supplementary Fig. S2 and S3 show the value of the bounds for various combinations of hyperparameter values. Clearly, some combinations lead to smaller bound values and thus bound-based model selection is possible. Additionally, Supplementary Tables S3 and S4 show the best bound values achieved for each benchmark dataset. In summary, these results indicate that the proposed sample compression bounds are sufficiently tight to be used in model selection and can be used to drastically reduce the training time of the CART and SCM algorithms, facilitating their scaling to large *genotype-to-phenotype* problems.

**Table 3.**
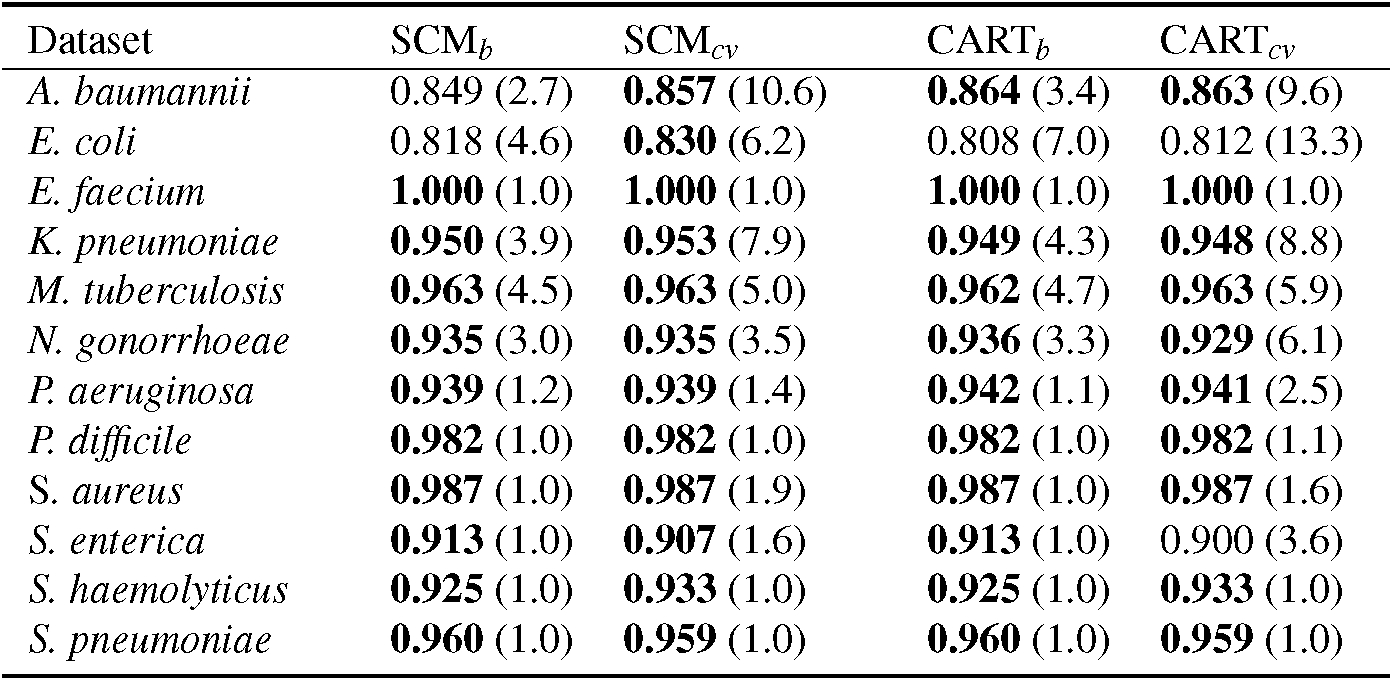
Comparison of models learned using bound selection (CART_*b*_, SCM_*b*_) and cross-validation (CART_*cv*_, SCM_*cv*_) as model selection strategies. For each dataset the accuracy is shown, along with the number of *k*-mers used by the model (in parentheses). Accuracies within 1% of the maximum value are shown in bold.

| Dataset | SCM<sub>b</sub> | SCM<sub>cv</sub> | CART<sub>b</sub> | CART<sub>cv</sub> |
| --- | --- | --- | --- | --- |
| *A. baumannii* | 0.849 (2.7) | **0.857** (10.6) | **0.864** (3.4) | **0.863** (9.6) |
| *E. coli* | 0.818 (4.6) | **0.830** (6.2) | 0.808 (7.0) | 0.812 (13.3) |
| *E. faecium* | **1.000** (1.0) | **1.000** (1.0) | **1.000** (1.0) | **1.000** (1.0) |
| *K. pneumoniae* | **0.950** (3.9) | **0.953** (7.9) | **0.949** (4.3) | **0.948** (8.8) |
| *M. tuberculosis* | **0.963** (4.5) | **0.963** (5.0) | **0.962** (4.7) | **0.963** (5.9) |
| *N. gonorrhoeae* | **0.935** (3.0) | **0.935** (3.5) | **0.936** (3.3) | **0.929** (6.1) |
| *P. aeruginosa* | **0.939** (1.2) | **0.939** (1.4) | **0.942** (1.1) | **0.941** (2.5) |
| *P. difficile* | **0.982** (1.0) | **0.982** (1.0) | **0.982** (1.0) | **0.982** (1.1) |
| *S. aureus* | **0.987** (1.0) | **0.987** (1.9) | **0.987** (1.0) | **0.987** (1.6) |
| *S. enterica* | **0.913** (1.0) | **0.907** (1.6) | **0.913** (1.0) | 0.900 (3.6) |
| *S. haemolyticus* | **0.925** (1.0) | **0.933** (1.0) | **0.925** (1.0) | **0.933** (1.0) |
| *S. pneumoniae* | **0.960** (1.0) | **0.959** (1.0) | **0.960** (1.0) | **0.959** (1.0) |

**Figure 4.**
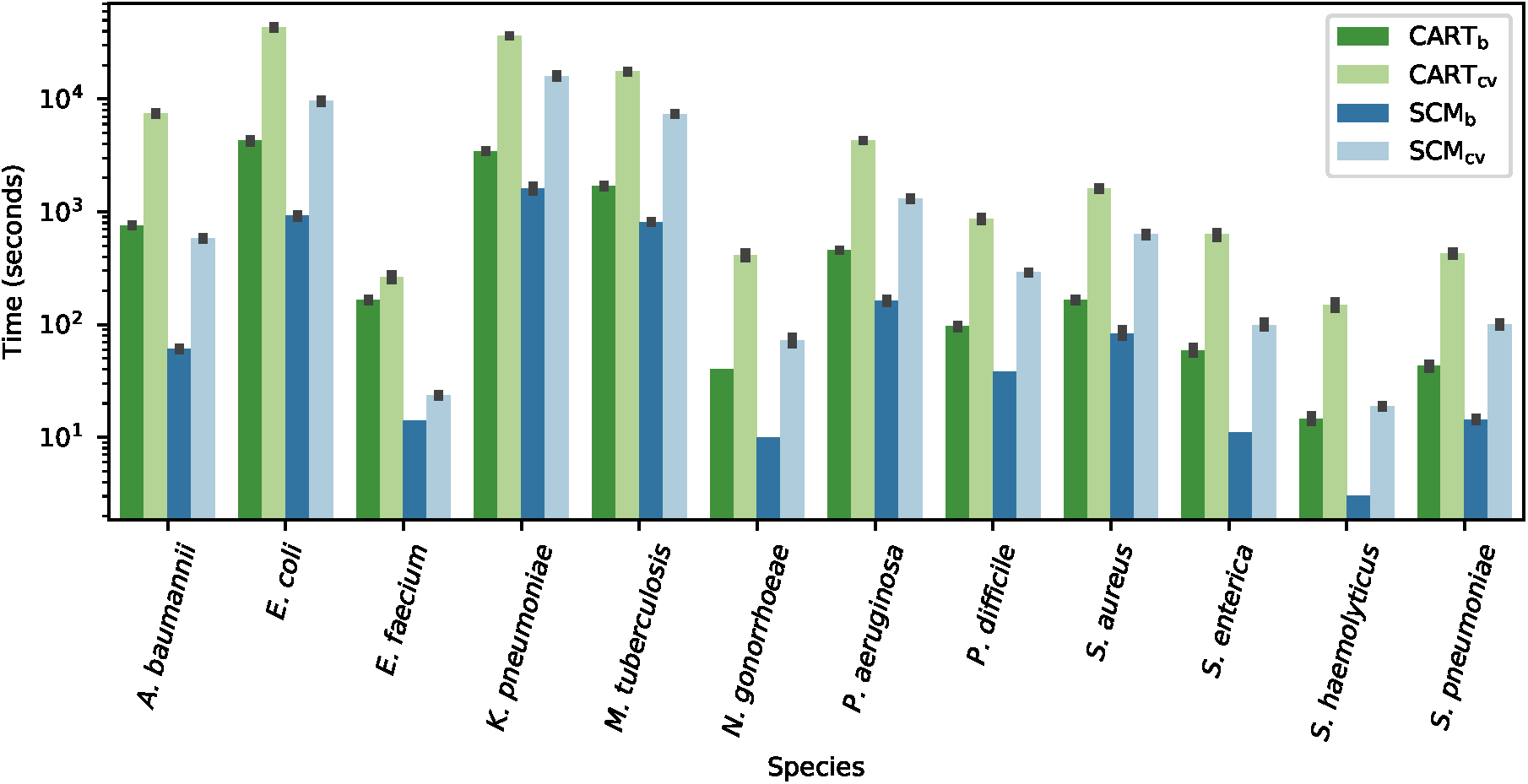
Running time (in seconds) of each algorithm and each model selection strategy on the benchmark datasets. For most datasets, using the sample compression bounds as a model selection strategy considerably reduces the running time compared to ten-fold cross-validation. SCM tends to run faster than CART, which was expected given that for models of equal depth, CART must evaluate many more rules than SCM. Of note, the time is shown on a logarithmic scale and thus, a difference in one unit corresponds to a running time that is ten times smaller.

### Multi-class classification with decision trees

This work has been thus far concerned with the prediction of binary phenotypes (e.g., case vs. control, resistant vs. susceptible). Yet, many phenotypes of practical importance are composed of more than two states. While Set Covering Machines do not directly support more than two classes, CART can work with an arbitrary number of classes. To demonstrate this property of CART, two multi-class antibiotic resistance prediction tasks were created. Specifically, the genomes of *Klebsiella pneumoniae* isolates with susceptible (S), intermediate (I), or resistant (R) phenotypes were collected to create three-class classification datasets, where the task consisted of discriminating between each level of resistance. This resulted in two datasets: gentamycin (2222 genomes, 74 million *k*-mers) and tobramycin (2068 genomes, 71 million *k*-mers). The accuracy of the resulting models is shown in Fig. 5a-b. Highly accurate predictions were obtained for the resistant and susceptible phenotypes, but not for the intermediate one. Interestingly, many intermediate isolates were predicted as resistant or susceptible, but the converse rarely occurred. We hypothesize that this is due to the presence of resistant and susceptible isolates that are mislabeled as intermediate in the data. In fact, according to CLSI testing standards^43^, the minimum inhibitory concentration (MIC) breakpoint for the intermediate class for both antibiotics is flanked by the resistant and susceptible class breakpoints within one two-fold dilution on each side. Given that this corresponds to the typical accuracy of MIC measurements^44^, it is likely that the intermediate class contains a fair amount of noise. Therefore, to strengthen our claim that CART_*b*_ can learn accurate multi-class models, we performed another experiment in which 100 genomes of each species were used to create a dataset where the task was to classify each genome into its correct species. This resulted in a dataset with 1200 genomes, 136 million *k*-mers, and 12 classes. The results, shown in Fig. 5c, indicate that a near perfect model was learned. Although solving this task based on the presence/absence of *k*-mers is not particularly challenging, the results clearly demonstrate that CART_*b*_ can learn highly accurate multi-class models.

**Figure 5.**
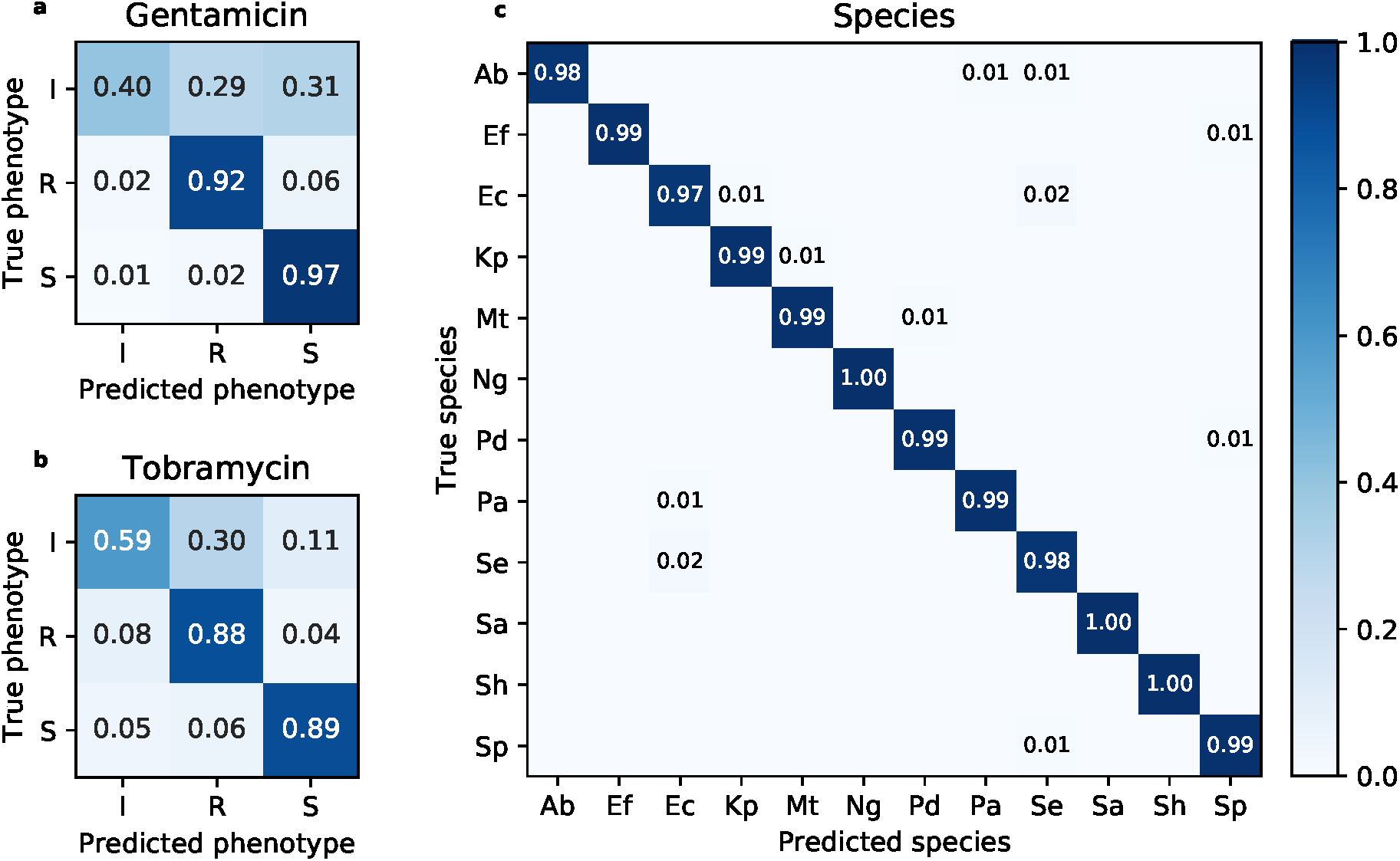
Confusion matrices for the multi-class classification tasks. a-b) Prediction of susceptible (S), intermediate (I), and resistant (R) AMR phenotypes for gentamicin and tobramycin (*K. pneumoniae*), respectively. c) Multi-species classification of the twelve species present in our dataset. Results are averaged over ten repetitions of the experiments.

## Discussion

Accurately predicting phenotypes from genotypes is a problem of high significance for biology that comes with great challenges for learning algorithms. Difficulties arise when learning from high dimensional genomic data with sample sizes that are minute in comparison^23^. Furthermore, the ability of experts to understand the resulting models is paramount and is not possible with most state-of-the-art algorithms. This study has shown the CART and SCM rule-based learning algorithms can meet these challenges and successfully learn highly accurate and interpretable genotype-to-phenotype models.

Notably, accurate genotype-to-phenotype models were obtained for 107 antimicrobial resistance phenotypes, spanning 12 eukaryotic species and 56 antimicrobial agents, which is an unprecedented scale for a machine learning analysis of this problem^19^. The obtained models were shown to be highly interpretable and to rely on confirmed drug resistance mechanisms, which were recovered by the algorithms without any prior knowledge of the genome. In addition, the models highlight previously unreported mechanisms, which remain to be investigated. Hence, the learned models are provided as *Additional data* with the hope that they will seed new research in understanding and diagnosing AMR phenotypes. A tutorial explaining how to visualize and annotate the models is also included.

Furthermore, a theoretical analysis of the CART and SCM algorithms, based on sample compression theory, revealed strong guarantees on the accuracy of the obtained models. Such guarantees are essential if models are to be applied in diagnosis or prognosis^23^. To date, these algorithms are among those that perform the highest degree of sample compression and thus, they currently provide the strongest guarantees (in terms of a sample compression risk bounds) for applications to high dimensional genomic data. Moreover, it was shown that these guarantees can be used for model selection, leading to significantly reduced learning times and models with increased interpretability. This serves as a good example of how theoretical machine learning research can be transferred to practical applications of high significance.

Finally, it is important to mention the generality of the proposed method, which makes no assumption on the species and phenotypes under study, except that the phenotypes must be categorical. The same algorithms could be used to predict phenotypes of tumor cells based on their genotype (e.g., malignant vs. benign, drug resistance), or to make predictions based on metagenomic data. To facilitate further biological applications of this work, an open-source implementation of the method, that does not require prior knowledge of machine learning, is provided with this work, along with comprehensive tutorials (see *Additional data*). The implementation is highly optimized and the algorithms are trained without loading all the genomic data into the computer’s memory.

Several extensions to this work are envisaged. The algorithms and their performance guarantees could be adapted to other types of representations for genomic variants, such as single nucleotide polymorphisms (SNP) and unitigs^45^. The techniques proposed by Hardt et al. (2016)^46^ could be used to ensure that the models are not biased towards undesirable covariates, such as population structure^47, 48^. This could potentially increase the interpretability of models, by avoiding the inclusion of rules that are associated with biases in the data. In addition, it would be interesting to generalize this work to continuous phenotypes, such as the prediction of minimum inhibitory concentrations in AMR^20^. Furthermore, another extension would be the integration of multiple omic data types to model phenotypes that result from variations at multiple molecular levels^49^. Additionally, this work could serve as a basis for efficient ensemble methods for genotype-to-phenotype prediction, such as random forest classifiers^50^, which could improve the accuracy of the resulting models, but would complexify the interpretation. Last but not least, the rule-based methods presented here assure good generalization if sparse sample-compressed classifiers with small empirical errors can be found. Nevertheless, it is known that good generalization can also be achieved in very high dimensional spaces with other learning strategies, such as achieving a large separating margin^51, 52^ on a large subset of examples or by using learning algorithms that are algorithmically stable^53^. Although it remains a challenge to obtain interpretable models with these learning approaches, they could eventually be useful to measure the extent to which the rule-based methods are losing predictive power at the expense of interpretability.

## Methods

### Data acquisition

The data were extracted from the Pathosystems Resource Integration Center (PATRIC) database^10, 24^ FTP backend using the *PATRIC tools* Python package^54^ (February 4, 2018). First, AMR phenotypes taking the form of SIR (susceptible, intermediate, resistant) labels were extracted for several bacterial isolates and antibiotics. Isolates associated with the intermediate phenotype were not considered, except in the multi-class experiments, to form two groups of phenotypically distinct isolates. Second, the metadata were segmented by species and antibiotic to form datasets, each corresponding to a single antibiotic/species pair. Datasets containing at least 25 isolates of each phenotype (107 in total) were retained and the rest were discarded. Third, the genomes in each dataset were downloaded, using the preassembled versions provided by PATRIC.

### A contextual introduction to supervised machine learning

Machine learning is a subfield of computer science that aims to create algorithms that learn from experience. Such algorithms learn how to perform tasks by analyzing a set of examples. In supervised learning, each example consists of an input and an expected outcome. The goal of the algorithm is to learn a model that accurately maps any input to the correct outcome. In the present study, it is assumed that the inputs are genomes and the expected outcomes are discrete phenotypes (e.g., resistance vs. susceptibility to an antimicrobial agent). Formally, let **x** *∈* {*A,C,G,T*}^*\**^ be a genome, represented by any string of DNA nucleotides, and *y ∈* {*p*_1_, *…, p*_*c*_} be its corresponding phenotype, where {*p*_1_, *…, p*_*c*_} is any set of *c* arbitrary phenotypes. The learning algorithm is given a set of examples 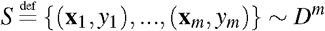, where each example (**x**_*i*_, *y*_*i*_) is generated independently according to the same distribution *D*. This distribution is unobserved and represents the unknown factors that generate the data (e.g., the biological mechanisms that underlie a phenotype). Learning algorithms often work with a vector representation of data (our case) and thus, it is necessary to transform the genome sequences into vectors. Let ***φ*** : {*A,C, G, T*} ^*\**^ → ℝ ^*d*^ be an arbitrary function that maps a genome to a vector of *d* dimensions. In this study, ***φ*** (**x**) is a *k*-mer profile (described below) that characterizes the presence and absence of every *k*-mer in the genome. The objective of the algorithm is to learn a model *h* : ℝ ^*d*^ → {*p*_1_, *…, p*_*c*_} that accurately maps the representation of a genome to its phenotype, i.e., *h*(***φ*** (**x**)) ≈ *y*. This corresponds to minimizing the expected error for any example drawn according to the data-generating distribution, defined as:

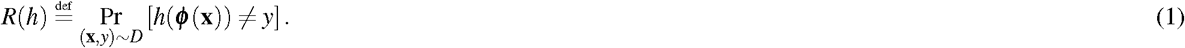

### The *k*-mer profile

Genome sequences are often represented as sets of *single nucleotide polymorphisms* (SNP), which are variations that occur at a single base pair location within a population^55–57^. This approach relies on multiple sequence alignment, which is computationally expensive and can fail in the presence of large-scale genomic rearrangements, such as horizontal gene transfer, that are common in bacterial populations^58–62^. In contrast, *reference-free* methods that represent each genome by a set of words, alleviate the need for multiple sequence alignment^58–62^. For instance, in *k*-mer-based representations, each genome is characterized by the set of *k*-mers (i.e., short words of *k* nucleotides) that it contains. Genomes can then be compared based on the presence and absence of such words. This approach is computationally efficient, since the representation can be computed independently, in parallel, for each genome. However, its main downside is that the representation contains a lot of redundancy, due to the fact the many k-mers are always present or absent simultaneously (e.g., gene deletion/insertion). In this sense, Jaillard et al. (2017)^45^ and Jaillard et al. (2018)^63^ proposed to replace *k*-mers by *unitigs*, i.e., words of variable length with unique presence/absence patterns that are generated using compacted De Bruijn graphs. In this study, we adopt a classical *k*-mer-based representation (referred to as *k*-mer profile) due to its simplicity and effectiveness. Nonetheless, it is important to note that the proposed algorithms could be adapted to work with other representations, such as SNPs and unitigs.

A *k*-mer profile is a vector of binary values that characterizes the presence or absence of every possible sequence of *k* DNA nucleotides in a genome. In theory, the dimension of *k*-mer profiles is 4^*k*^, which is approximately 4.6 × 10^18^ for *k* = 31. However, in practice, *k*-mers that do not occur in the set of genomes to be compared can be omitted since they cannot be used in the model^17^. This dramatically reduces the number of possible *k*-mers and thus, the size of the representation. Formally, let *K* be the set of all (possibly overlapping) *k*-mers that occur more than once in the genomes of a dataset *S*. For a genome **x** *∈*{*A,C,G,T*} ^*\**^, the corresponding *k*-mer profile is given by the |*K*|-dimensional boolean vector ***φ*** (**x**) *∈* 0, 1 ^|*K*|^, such that *φ* (**x**)_*i*_ = 1 if *k*-mer *k*_*i*_ ∈ *K* appears in **x** and *φ* (**x**)_*i*_ = 0 otherwise. In this work, the *k*-mers in each genome were determined using the DSK *k*-mer counter^64^ and the length *k* was set to 31, since extensive experiments showed that this length was appropriate for bacterial genome comparison^17, 65^. The reader is referred to Drouin et al. (2016)^17^ for an illustration of the *k*-mer profile and a discussion on choosing an appropriate *k*-mer length. While the practical size of *k*-mer profiles is much smaller than their theoretical limit, they remain extremely high dimensional data representations that push the limits of current learning algorithms in terms of scalability and generalization.

### Boolean-valued rules based on *k***-mers**

The rule-based algorithms used in this work learn models that are arrangements of *boolean-valued rules*. Such rules take a *k*-mer profile as input and output *true* or *false*. We consider one *presence rule* and one *absence rule* for each *k*-mer *k*_*i*_ ∈ *K*, which are defined as 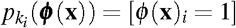 and 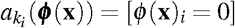, respectively. The goal of the learning algorithms is to find the arrangement of such rules that gives the most accurate predictions of the phenotype. The resulting models are interpretable and directly highlight the importance of using a small set of *k*-mers.

### Performance guarantees based on sample compression theory

A generalization bound (or risk bound) is a function *ε*(*h, S, δ*) that depends on what the model *h* achieves on the training set *S*. Such a function upper bounds, with probability at least 1 *-δ* (over the random draws of *S* according to *D*^*m*^), the generalization error of *h* as defined by Equation (1). Among other things, *ε*(*h, S, δ*) depends on the training error of *h* on *S* and on its complexity (as measured here by the number of rules used by *h* and its *sample-compression size*^*66*^). Furthermore, *ε*(*h, S, δ*) is valid simultaneously for all possible predictors *h* that can be constructed by the learning algorithm, but *ε*(*h, S, δ*) increases with the complexity of *h* and its training error. The guarantee on the generalization error of *h* thus deteriorates as the model *h* becomes complex and/or inaccurate on the training data. Consequently, such a bound *ε*(*h, S, δ*) can be analyzed to understand what should be optimized in order to learn models that achieve good generalization.

Drouin et al. (2016) proposed a generalization bound for the SCM algorithm, which shed light on this algorithm’s strong resistance to overfitting in the challenging genotype-to-phenotype setting^17^. Based on their work, we propose a new bound for the CART algorithm and demonstrate that its models can also achieve good generalization in this setting. Together, these theoretical results support the empirical results reported in this study and strengthen our claim that rule-based classifiers are well-suited for genotype-to-phenotype studies.

The generalization bound *ε*_*SCM*_(*h, S, δ*) proposed by Drouin et al. (2016)^17^ for the SCM is given by

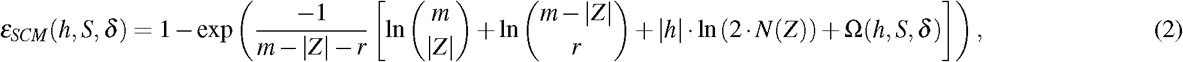

where *m* is the number of training examples in *S*, and | *h* | is the number of rules in the conjunction/disjunction model. Furthermore, *Z*, called the *compression set*^21, 22, 66^, is a small subset of *S* in which all *k*-mers used by the model occur. We denote by |*Z*|, the number of genomes in *Z*, and by *N*(*Z*) the total number of nucleotides in *Z*. Moreover, *r* is the number of prediction errors made by *h* on *S\ Z*, i.e., the examples of the training set that are not in *Z*. Finally,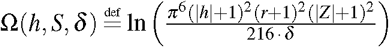.

Using the same definitions and notation, the generalization bound *ε*_*CART*_ (*h, S, δ*) that we propose for CART is given by

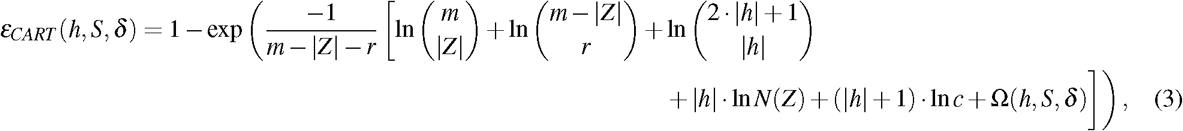

where *c* is the number of classes in the data. Interestingly, this bound shares many terms in common with the bound for the SCM, but also supports the multi-class setting. A detailed derivation of this bound is given in the Supplementary Methods, along with a discussion of related work on generalization bounds for decision tree models. Of note, we compare our bound to a related sample compression bound for decision tree models^67^ and show that it is better-suited for applications to genomic data.

These bounds indicate that any model that makes few errors on the training data, while using a small number of rules should achieve good generalization. This is precisely the type of models that were obtained in the experiments of this work. Surprisingly, the length *k* of the *k*-mers, which increases the size of the feature space exponentially, does not appear in these equations. This indicates that SCM and CART can achieve good generalization despite the immense dimensionality (4^*k*^) of the feature space that results from using large values of *k*. This property makes them ideal for genotype-to-phenotype prediction.

### Fast model selection with bounds

Model selection consists of choosing the configuration of the learning algorithm that yields the model with the smallest generalization error, as defined by Equation (1). Such a configuration is an arrangement of user-defined parameters that control the behavior of the algorithm, which are referred to as *hyperparameters* (HPs). For instance, in the SCM algorithm, the maximum number of rules in a conjunction/disjunction model is a HP^9^. Similarly, in the CART algorithm, the minimum cost-complexity pruning algorithm is used to reduce the complexity of the resulting models and the level of pruning is controlled by a HP^8^.

For settings where the data is scarce, such as genotype-to-phenotype studies, model selection is typically performed using *k*-fold cross-validation (see Hastie et al. (2015) for an introduction^68^). This method consists of partitioning the training data into *k* (typically 5 or 10) disjoint subsets of equal size (referred to as folds) and training the algorithm *k* times, each time using *k*-1 folds for training and the remaining one for validation. The score attributed to each configuration of the algorithm is the proportion of prediction errors in validation over all folds, which is an empirical estimation of the generalization error given by Equation (1), and the one with the smallest score is selected. This procedure has two limitations: it is computationally intensive, since the algorithm is trained *k* times for each configuration, and it requires that some data be left out for validation.

An alternative method, referred to as *bound selection*, consists of using a generalization bound, such as those described at Equations (2) and (3), in replacement for cross-validation^9, 17^. The idea is to train the algorithm with several configurations and score them using the bound value of the resulting model. The configuration that leads to the smallest bound value is retained. This approach is computationally and data efficient, since it requires a single training of the algorithm for each configuration and does not require that data be left out for validation. The reader is referred to Supplementary Fig. S4 for a comparative illustration of cross-validation and bound selection.

In this work, both model selection approaches were used to train the CART and SCM algorithm, resulting in the CART_*b*_ and SCM_*b*_ algorithms (bound selection) and the CART_*cv*_ and SCM_*cv*_ algorithms (ten-fold cross-validation). It was observed that models obtained using bound selection were just as accurate as those obtained using cross-validation, but that they were significantly less complex and thus, more interpretable (see *Results*). The configurations selected for each dataset are provided as *Additional data*.

### Kover: a scalable disk-based implementation

The rule-based algorithms used in this work are implemented in Kover (https://github.com/aldro61/kover/). Kover is an open-source bioinformatics software that allows practitioners, with no prior knowledge of machine learning, to learn rule-based models of phenotypes from their data. It accepts genomes in the form of sequences (reads or contigs) or precomputed *k*-mer profiles. In the former case, the genomes are converted to *k*-mer profiles using the DSK *k*-mer counter. Kover automates the machine learning analysis (e.g., model selection, model evaluation), which ensures that proper protocols are followed. It produces detailed reports, which contain the learned models, along with several metrics assessing their accuracy. A detailed tutorial is provided as *Additional information*.

From a computational perspective, the particularity of Kover is that the learning algorithms are trained *out-of-core*, which means that the dataset is never entirely loaded into the computer’s memory. This is achieved through the careful use of HDF5 and data chunking patterns^17, 69^. In addition, Kover relies on the *popcount* atomic CPU instruction to train the algorithms directly from a compressed representation of the *k*-mer profiles, resulting in lesser memory requirements and faster computations^17^. These properties allow Kover to scale to datasets with sizes well beyond those encountered in this study and make it a tool of choice for large-scale genotype-to-phenotype studies based on machine learning.

### Comparison to state-of-the-art classifiers

#### Benchmark datasets

One dataset per species was included in the benchmark and the datasets were selected to have the largest possible sample size with low class imbalance. For each species, all datasets with less than 20% class imbalance, defined as:

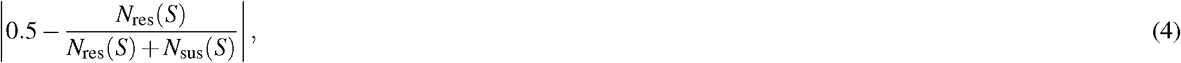

where *N*_res_(*S*) and *N*_sus_(*S*) are respectively the number of examples with the resistant and susceptible phenotype in *S*, were considered and the one with the most examples was selected. The only exception is *Salmonella enterica*, where the most balanced dataset had 22% class imbalance.

#### Selected algorithms

The benchmark includes a comparison to five learning algorithms and their choice is motivated hereafter.

#### Logistic regression

This algorithm produces linear classifiers that estimate the probability that an example belongs to each class. It can be used with an *L*1-norm regularizer to obtain sparse models that rely on a subset of *k*-mers (L1-logistic). It is interesting to compare the sparsity of these models to those of CART and SCM. Additionally, this algorithm can be used with an *L*2-norm regularizer to obtain dense models (L2-logistic) which serve to illustrate that sparsity does not have a detrimental effect on accuracy. The implementation in Scikit-Learn^70^ was used. In sharp contrast with Kover, it requires that the entire datasets be stored in the computer’s memory, which is intractable for the datasets used in this study. Hence, the one million features that were most associated with the phenotype were selected using a univariate feature selection with a *χ*^2^ test^71, 72^ and the others were discarded.

#### Polynomial Kernel Support Vector Machine

This kernel-based learning algorithms yields non-linear classifiers that, when trained with binary *k*-mer profiles, correspond to a majority vote of all possible conjunctions of *d k*-mer presence rules, where *d* is the degree of the polynomial. It is thus particularly relevant to compare this algorithm to SCM_*b*_, which seeks the single, most accurate, conjunction. The implementation in Scikit-Learn^70^ was used and the kernel was computed using powers of the pairwise similarity (dot product of *k*-mer profiles) matrix of genomes. No feature selection was used, since the memory requirements of this algorithm are small (*m*× *m* for *m* learning examples).

#### Naive Bayes

This baseline algorithm assumes that each input feature is statistically independent given the class (which is generally false). This approach is computationally efficient in high dimensions, since it assumes that class densities are simply given by the product of marginal densities; justifying its use in our context. A custom implementation was used and the code is available as *Additional information*.

#### Majority

This baseline algorithm is used to ensure that the algorithms successfully identify predictive patterns in the data. An algorithm that underperforms this baseline could have achieved better results without attempting to learn anything.

## Acknowledgements

The authors acknowledge Mathieu Blanchette, Christopher J.F. Cameron, Maia Kaplan, and Pier-Luc Plante for valuable comments and suggestions. This work was supported in part by an Alexander Graham Bell Canada Graduate Scholarship Doctoral Award of the Natural Sciences and Engineering Research Council of Canada (NSERC) to AD, an Alexander Graham Bell Canada Graduate Scholarship Master’s award (NSERC) to GL, the NSERC Discovery Grants (FL; 262067, MM; RGPIN-2016-05942), and the Canada Research Chair in Medical Genomics (JC). FR is associated with the Canada Research Excellence Chair in the Microbiome-Endocannabinoidome Axis in Metabolic Health. This research was enabled in part by support provided by Calcul Québec (www.calculquebec.ca) and Compute Canada (www.computecanada.ca). Computations were performed on the Colosse (Laval University) and Graham (University of Waterloo) supercomputers under resource allocation projects nne-790-af and agq-973-ac.

## Author contributions statement

A.D. and G.L. conceived and conducted the experiments, A.D., F.R., G.L., and J.C. analyzed the results, A.D., F.L., G.L., and M.M. did the theoretical analysis of the learning algorithms, A.D. and G.L. implemented the learning algorithms in Kover. All authors reviewed the manuscript.

## Additional information

### Additional data

The 2140 AMR prediction models learned with CART_*b*_ and SCM_*b*_ are provided with code for their visualization and guidelines for their interpretation at https://github.com/aldro61/kover2_paper.

### Tutorials

Detailed tutorials on using Kover for genotype-to-phenotype prediction are available at https://aldro61.github.io/kover

### Reproducibility

The code used to acquire the data and run the experiments, as well as detailed experimental results for each algorithm are available at https://github.com/aldro61/kover2_paper.

