## Supplementary Material for "Interpretable genotype-to-phenotype classifiers with performance guarantees"

### Supplementary information

**Alexandre Drouin[1,2,*], Gaël Letarte[1,2], Frédéric Raymond[3,4], Mario Marchand[1,2], Jacques Corbeil[2,5], and François Laviolette[1,2]**

[1]Department of Computer Science and Software Engineering, Université Laval, Quebec, Canada
[2]Big Data Research Centre, Université Laval, Quebec, Canada
[3]School of nutrition, Université Laval, Quebec, Canada
[4]Institute of Nutrition and Functional Foods, Université Laval, Quebec, Canada
[5]Infectious Disease Research Centre, Université Laval, Quebec, Canada
[*]

### Supplementary methods

#### A sample compression risk bound for decision trees

Based on the pioneering work of Littlestone and Warmuth (1986)[1] and Floyd and Warmuth (1995)[2], Marchand and Sokolova (2005)[3] obtained a general sample compression bound that can be used to upper bound the generalization error (see Equation (1) of main text) of any classifier $h$, such that $h = \mathbf{R}(Z, \sigma)$, where $\mathbf{R}$ is a reconstruction function that unambiguously reconstructs $h$ using a small subset $Z$ of the training examples (referred to as the *compression set*) and a message $\sigma$ of additional information. Their bound is as follows: for any data-generating distribution $D$, any compression set $Z$ and message $\sigma$, we have that, with probability at least $1-\delta$ (over the random draws of $S$ according to $D^m$), $R(h) \leq \varepsilon(h, S, \delta)$, with

$$\varepsilon(h,S,\delta) = 1 - \exp\left(\frac{-1}{m-|Z|-r}\left[\ln\binom{m}{|Z|} + \ln\binom{m-|Z|}{r} + \ln\left(\frac{1}{P_Z(\sigma)}\right) + \ln\left(\frac{1}{\xi(|Z|)\xi(r)\delta}\right)\right]\right), \quad \text{(S1)}$$

where $Z \subseteq S$, $P_Z(\sigma)$ is the prior probability assigned to the message $\sigma$ given that the compression set is $Z$, $|Z|$ denotes the number of examples in the compression set $Z$, $r$ is the number of prediction errors made by $h$ on $S \setminus Z$, and

$$\xi(a) \stackrel{\text{def}}{=} \frac{6}{\pi^2}(a+1)^{-2}. \quad \text{(S2)}$$

In order to use this result to obtain a sample compression bound for $k$-mer-based decision tree models used in this study, we must design a message $\sigma$, and a corresponding compression set $Z$, that jointly allow to unambiguously reconstruct any decision tree classifier $h$. Recall from the main text that $Z$ contains the genomes selected such that every $k$-mer in the model appears at least once in $Z$. Recall also that $N(Z)$ denotes the number of nucleotides contained in $Z$.

Our approach relies on the fact that any tree with $n$ inner nodes admits a unique preorder enumeration of its $2n+1$ nodes ($n$ inner nodes and $n+1$ leaves). We also consider that each message $\sigma$ is given by a tuple $(n, \mathbf{v_1}, \mathbf{v_2}, \mathbf{v_3})$, where

- $\mathbf{v_1} \in \{0,1\}^{2n+1}$ is a vector that gives the type of each node in the enumeration, such that $v_{1_i} \stackrel{\text{def}}{=} 1$ if the $i^{\text{th}}$ node is an inner node and $v_{1_i} \stackrel{\text{def}}{=} 0$ otherwise.
- $\mathbf{v_2} \in \{1,...,c\}^{n+1}$ is a vector indicating the class predicted by each leaf in the enumeration; $c$ is the number of classes.
- $\mathbf{v_3} \in \{1,...,N(Z)\}^n$ is a vector that specifies the $k$-mer used by each inner node (rule) in the enumeration, based on its position in the concatenated sequence of all genomes in $Z$.

Any decision tree $h$ can then be straightforwardly reconstructed from any compression set $Z$ and any such message tuple.

To obtain a generalization error bound, we must also define a prior probability distribution $P_Z(\sigma)$ over all possible values of $\sigma$, given a compression set $Z$. We start by attributing a probability of $\xi(n)$ to the number of inner nodes. Thus,

$$P_Z(\sigma) = P_Z(\sigma|n) \cdot \xi(n), \tag{S3}$$

which reflects our prior belief that smaller trees are more likely than large ones. We then assign equal probability to all messages specifying trees of $n$ inner nodes:

$$P_Z(\sigma|n) = P_1(\mathbf{v_1}) \cdot P_2(\mathbf{v_2}) \cdot P_3(\mathbf{v_3}), \tag{S4}$$

where $P_1$, $P_2$, and $P_3$ are chosen as follows. First,

$$P_1(\mathbf{v_1}) = \frac{1}{\binom{2n+1}{n}}, \tag{S5}$$

which assigns equal probability to all vectors $\mathbf{v_1}$ with $n$ elements equal to 1 and $n+1$ elements equal to 0. Then

$$P_2(\mathbf{v_2}) = \left(\frac{1}{c}\right)^{n+1}, \tag{S6}$$

which assigns equal probability to each class for the $n+1$ leaves. Finally,

$$P_3(\mathbf{v_3}) = \left(\frac{1}{N(Z)}\right)^{n}, \tag{S7}$$

which assigns equal probability over all positions in the combined sequence of all genomes in $Z$ for every inner node. Hence, we obtain a prior $P_Z(\sigma)$, where

$$P_Z(\sigma) = \frac{6}{\pi^2}(n+1)^{-2}\binom{2n+1}{n}^{-1}\left(\frac{1}{N(Z)}\right)^{n}\left(\frac{1}{c}\right)^{n+1}. \tag{S8}$$

By inserting this prior $P_Z(\sigma)$ into Equation S1, we obtain a sample compression risk bound $\varepsilon_{CART}(h,S,\delta)$, which valid for any decision tree $h$ based on rules that detect the presence of $k$-mers:

$$\varepsilon_{CART}(h,S,\delta) = 1 - \exp\left(\frac{-1}{m-|Z|-r}\left[\ln\binom{m}{|Z|} + \ln\binom{m-|Z|}{r} + \ln\binom{2n+1}{n} + n\cdot\ln(N(Z)) + (n+1)\ln(c) + \ln\left(\frac{\pi^6(n+1)^2(r+1)^2(|Z|+1)^2}{216\cdot\delta}\right)\right]\right). \tag{S9}$$

In the main text, we use $|h|$, instead of $n$, for the number of rules (i.e., internal nodes) in the decision tree $h$.

### Related work on generalization bounds for decision trees

Several theoretical upper bounds on the risk of decision trees exist in the literature. Most of them are either based on the Vapnik-Chervonenkis dimension[4] (VC-dim) or the Rademacher complexity[5]. However, the tightness of these bounds is challenged in our setting due to the extremely high dimensionality of the input space. Given that our goal is to derive a generalization bound that is tight enough to guide model selection, including the pruning process of decision trees, such bounds are of limited interest. In fact, the VC-dim and Rademacher complexity are only defined for data-independent sets of classifiers. Given that we consider $k$-mers of length $k = 31$, we have to consider all decision trees of $n$ nodes (for small $n$) that can be constructed over $d = 4^{31}$ boolean variables (i.e., the presence or absence of a $k$-mer). According to the recent work of Yıldız (2015)[6], the VC-dim of decision trees of height $p$ is a least $2^{(p-1)}(1+\lfloor\log_2(d-p+2)\rfloor)$. Since $p$ is at least $\log_2(n)$, this gives a linear increase in $n$ with a large multiplier, since $d = 4^{31}$. Similarly, bounds based on Rademacher complexities exhibit the same difficulties[7].

One way to obtain tighter bounds for data-independent sets of classifiers is to use part of the training set to build the model and the remaining data to calculate a generalization bound. For instance, Kääriäinen et al. (2004)[8] use a fraction of the training set to build a (possibly very large) decision tree $T$ and then use the remaining data to prune it based on the Rademacher complexity of the set of subtrees of $T$. However, the problem with this approach is that fewer examples are used to grow the tree and then prune it.

Consequently, in order to use the full training set to grow and prune the tree, while achieving a tight bound, we have decided to investigate risk bounds for data-dependent sets of classifiers. Such sets of classifiers can be significantly more concise than their data-independent counterparts and thus, lead to tighter bounds. In our case, we consider the set of all decision trees composed exclusively of $k$-mers that are present in the data set (and not the full set of $4^{31}$ $k$-mers). In this sense, our best prospect was the sample compression bounds proposed by Floyd and Warmuth (1995)[2] and then later by Marchand and Sokolova (2005)[3] for conjunctions, disjunctions, and decision lists. Such bounds were recently explored by Drouin et al. (2016)[9] to obtain tight generalization bounds for conjunctions and disjunctions of $k$-mers learned using Set Covering Machines[10]. In the case of decision trees, the only sample compression bound that we are aware of is the one proposed by Shah (2007)[11]. However, this bound is too generic and no pruning algorithm based on it was ever proposed. Instead, following the work of Drouin et al. (2016)[9], we have decided to propose a tighter bound, specialized to decision trees of $k$-mers, that takes into account the particularities of this representation, i.e., that $k$-mers are substrings of genome sequences.

In fact, the main difference between our bound and the one of Shah (2007)[11] is in the way that the decision tree is encoded into a compression set and a message string. In their bound, the $k$-mers are considered as generic features and the message string contains the index of the $k$-mer on which each rule (inner node) relies. The compression set serves only to specify each rule's threshold (i.e., 0 or 1) based on the value of the feature in the corresponding example. In contrast, we exploit the fact that the $k$-mers are substrings of genomes and define the compression set as being the smallest set of genomes that contains all the $k$-mers in the model. We then use the message string to specify the index of the $k$-mers in the concatenated sequences of the genomes of the compression set. We also account for the fact that multiple $k$-mers can be found in the same genome, which leads to smaller compression sets and thus a tighter bound.

More formally, for a decision tree with $n$ inner nodes (rules) the bound of Shah (2007)[11] contains an additive term of $n \cdot \ln(d)$, where $d$ is the total number of k-mers (i.e., $4^k$), whereas, in our bound, this term is replaced by $n \cdot \ln(N(Z))$, where $N(Z)$ is the number of nucleotides in all the genomes of the compression set. Given that $|Z| \leq n$ and that we seek simple trees with few inner nodes (i.e., small $n$), $N(Z)$ is of the order of a few millions and is bound to be smaller than the total number of $k$-mers ($d$) when $k$ is sufficiently large (e.g., for $k = 31$, $d = 4^k \approx 4.6116\text{e}18$), resulting in a tighter bound.

### Supplementary figures

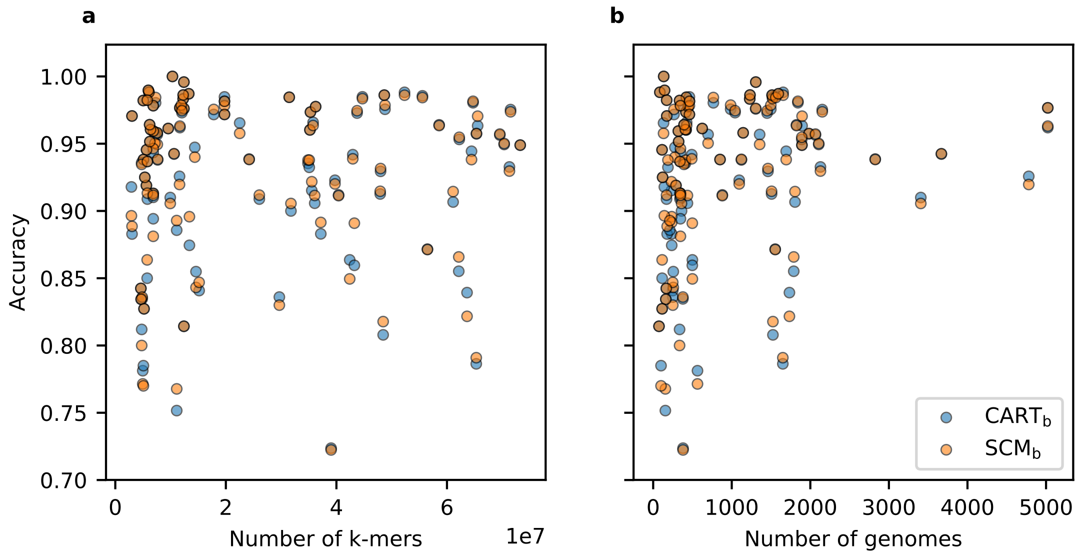


**Figure S1.** Accuracy of the CART$_b$ and SCM$_b$ models with respect to a) the number of $k$-mers and b) the number of genomes in each of the 107 datasets (shown as dots). Clearly, small numbers of genomes are not associated with poor accuracies. The same is true for large numbers of $k$-mers. These results emphasize the ability of these algorithms to achieve good generalization despite small samples sizes and extremely high dimensional data.

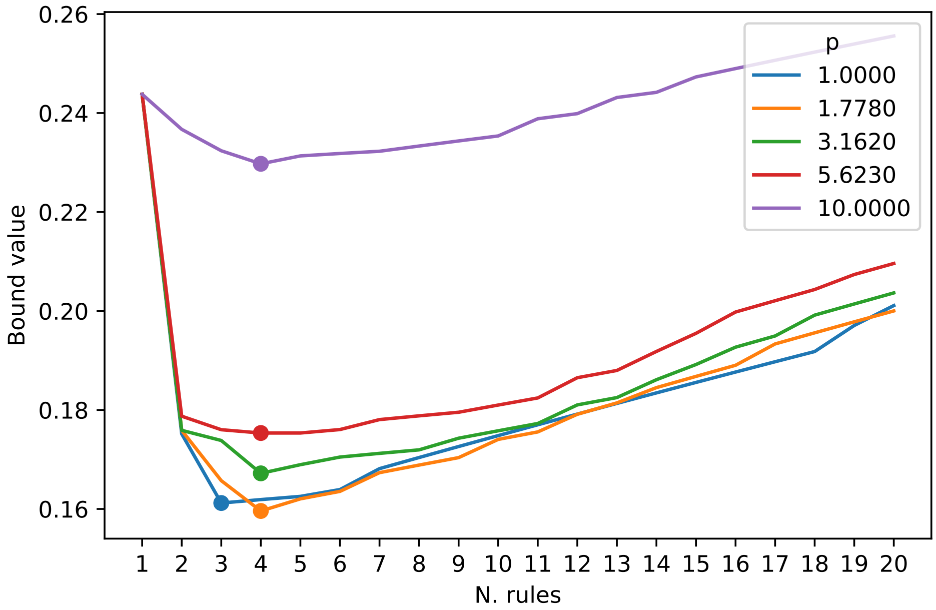


**Figure S2.** Value of the sample compression bound of the SCM$_b$ algorithm (Equation (2)) with respect to the number of rules in the model for the *M. tuberculosis* benchmark dataset. Each of the colored lines corresponds to a different value of the $p$ hyperparameter, which controls the importance of the positive and negative classes in the greedy optimization algorithm (see Marchand and Shawe-Taylor (2002)[10]). The minimum of each line is marked by a dot. For clarity, we only show results for disjunction models (logical-OR). Clearly, there is a well-distinguishable set of hyperparameter values that yield a smaller bound value than the others and the bound allows for model selection.

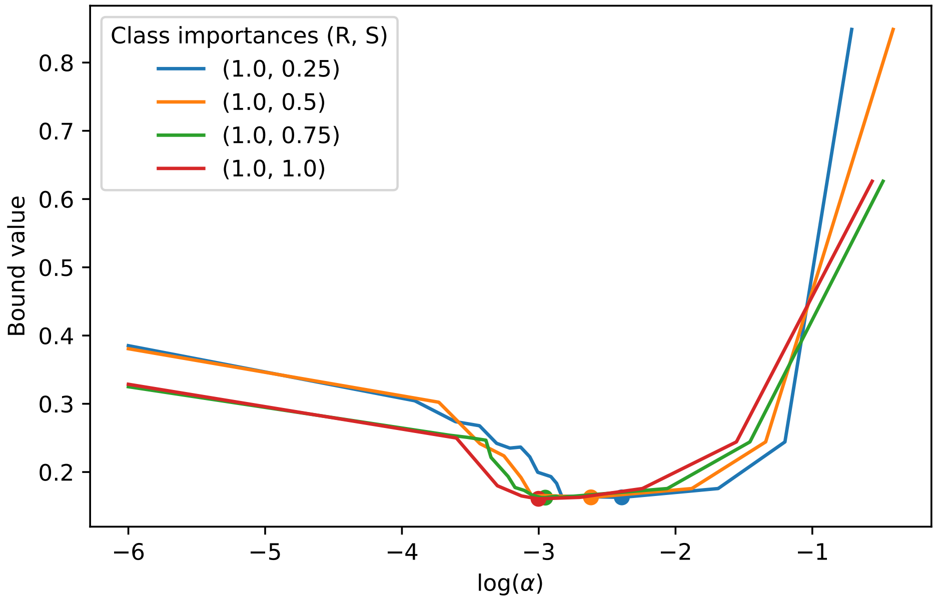


**Figure S3.** Value of the sample compression bound of the $\text{CART}_b$ algorithm (Equation (3)) with respect to the alpha hyperparameter of the minimum cost-complexity pruning algorithm of Breiman et al. (1984)[12], which controls the size of the resulting tree, for the *M. tuberculosis* benchmark dataset. Each of the colored lines corresponds to a different class importance ratio, which serves to increase the importance of making errors on any of the classes. The minimum of each line is marked by a dot. For clarity, we only show results while varying the importance of the susceptible (S) class. Clearly, there is a well-distinguishable set of hyperparameter values that yield a smaller bound value than the others and the bound allows for model selection.

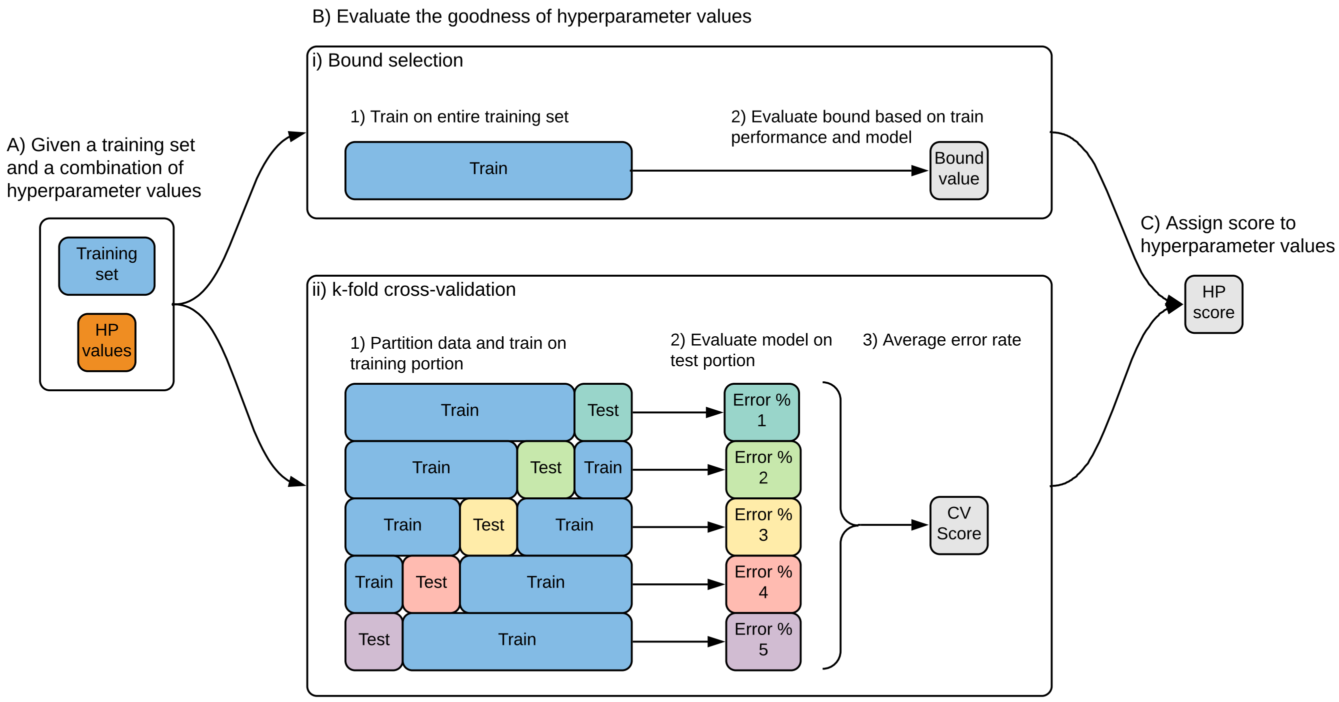


**Figure S4.** Illustration of the bound selection and cross-validation model selection methods. A) Both methods are given a combination of hyperparameter values to score, as well as a set of training data. B) The methods differ in the strategy that they use to compute a score. (i) Bound selection trains the algorithm on the entire training set and scores the hyperparameters based on the expression of a generalization bound. In this study, the bound depends on some properties of the model (e.g., complexity) and its performance on the training data (e.g., number of prediction errors). This requires a single training of the algorithm and all the data is used for training. (ii) In contrast, $k$-fold cross-validation creates $k$ partitions of the data and trains $k$ distinct models that are evaluated on $k$ testing sets (folds). This approach is less computationally efficient than bound selection and requires that some data be left out for testing. C) The score estimated by both methods is assigned to the combination of hyperparameter values and the combination with the best score (e.g., minimum value) is retained.

### Supplementary tables

**Table S1.** Detailed results for all datasets and the methods compared in Table 2 and Supplementary Table S2. For each dataset (species-antibiotic pair), the number of genomes (total, resistant, and susceptible) and *k*-mers is shown, along with the accuracy, sensitivity, specificity, F1 score, and the complexity of the models learned by each algorithm (average ± standard deviation for ten repetitions – see main text). The complexity is the number of *k*-mers used by the models, with all* indicating that feature selection was performed (see main text) and that the one million selected features were used. Missing F1 score values indicate that, in at least one repetition, the value of this metric was *nan* or infinite, which (in our case) can occur if no examples are predicted as positive or there are no true positive predictions.

| Species | Antibiotic | Genomes | Resistant | Susceptible | *k*-mers (millions) | Method | Accuracy | Sensitivity | Specificity | F1 score | Complexity |
|---|---|---|---|---|---|---|---|---|---|---|---|
| *A. baumannii* | amikacin | 256 | 195 | 61 | 14.6 | L1-logistic | 0.835 ± 0.064 | 0.880 ± 0.072 | 0.663 ± 0.192 | 0.893 ± 0.048 | 4575.7 ± 6046.6 |
| | | | | | | L2-logistic | **0.861 ± 0.051** | 0.890 ± 0.058 | **0.740 ± 0.155** | **0.909 ± 0.038** | all* |
| | | | | | | Majority | 0.790 ± 0.045 | **1.000 ± 0.000** | 0.000 ± 0.000 | 0.882 ± 0.029 | – |
| | | | | | | Naive Bayes | 0.725 ± 0.049 | 0.780 ± 0.051 | 0.533 ± 0.095 | 0.817 ± 0.036 | all |
| | | | | | | Poly-SVM | **0.865 ± 0.051** | 0.902 ± 0.056 | 0.717 ± 0.173 | **0.912 ± 0.038** | all |
| | | | | | | RBF-SVM | 0.851 ± 0.069 | 0.902 ± 0.058 | 0.660 ± 0.170 | 0.904 ± 0.048 | all |
| | | | | | | Random Forests | 0.843 ± 0.054 | 0.895 ± 0.060 | 0.635 ± 0.112 | 0.899 ± 0.039 | 6762.8 ± 7422.5 |
| | | | | | | CART$_b$ | 0.855 ± 0.041 | 0.898 ± 0.037 | 0.698 ± 0.179 | **0.907 ± 0.027** | 2.5 ± 0.5 |
| | | | | | | CART$_{cv}$ | **0.867 ± 0.041** | 0.905 ± 0.045 | 0.719 ± 0.188 | **0.914 ± 0.029** | 4.1 ± 1.7 |
| | | | | | | SCM$_b$ | 0.843 ± 0.044 | 0.878 ± 0.035 | 0.725 ± 0.129 | 0.898 ± 0.032 | 2.1 ± 0.3 |
| | | | | | | SCM$_{cv}$ | 0.837 ± 0.043 | 0.886 ± 0.048 | 0.615 ± 0.228 | 0.895 ± 0.030 | 5.5 ± 2.6 |
| | ampicillin/sulbactam | 155 | 111 | 44 | 11.1 | L1-logistic | 0.823 ± 0.067 | 0.842 ± 0.075 | **0.756 ± 0.190** | 0.876 ± 0.046 | 106216.1 ± 233553.8 |
| | | | | | | L2-logistic | 0.835 ± 0.056 | 0.867 ± 0.069 | 0.743 ± 0.160 | 0.887 ± 0.039 | all* |
| | | | | | | Majority | 0.748 ± 0.042 | **1.000 ± 0.000** | 0.000 ± 0.000 | 0.855 ± 0.028 | – |
| | | | | | | Naive Bayes | 0.797 ± 0.071 | 0.902 ± 0.082 | 0.492 ± 0.175 | 0.868 ± 0.050 | all |
| | | | | | | Poly-SVM | 0.829 ± 0.053 | 0.867 ± 0.059 | 0.722 ± 0.140 | 0.883 ± 0.040 | all |
| | | | | | | RBF-SVM | **0.848 ± 0.034** | 0.893 ± 0.063 | 0.706 ± 0.187 | **0.898 ± 0.024** | all |
| | | | | | | Random Forests | **0.839 ± 0.059** | 0.881 ± 0.081 | 0.706 ± 0.135 | **0.890 ± 0.042** | 1910.0 ± 3035.0 |
| | | | | | | CART$_b$ | 0.752 ± 0.046 | 0.811 ± 0.076 | 0.586 ± 0.285 | 0.830 ± 0.030 | 1.0 ± 0.0 |
| | | | | | | CART$_{cv}$ | 0.810 ± 0.084 | 0.846 ± 0.100 | 0.707 ± 0.141 | 0.867 ± 0.063 | 7.3 ± 2.9 |
| | | | | | | SCM$_b$ | 0.768 ± 0.060 | 0.841 ± 0.073 | 0.562 ± 0.227 | 0.843 ± 0.042 | 1.0 ± 0.0 |
| | | | | | | SCM$_{cv}$ | 0.787 ± 0.049 | 0.842 ± 0.067 | 0.624 ± 0.218 | 0.855 ± 0.031 | 5.6 ± 2.4 |
| | carbapenem | 232 | 122 | 110 | 35.5 | L1-logistic | 0.943 ± 0.040 | 0.937 ± 0.043 | 0.948 ± 0.047 | 0.949 ± 0.037 | 1075.4 ± 627.9 |
| | | | | | | L2-logistic | 0.943 ± 0.046 | 0.945 ± 0.048 | 0.942 ± 0.049 | 0.950 ± 0.038 | all* |
| | | | | | | Majority | 0.520 ± 0.094 | 0.900 ± 0.316 | 0.100 ± 0.316 | – | – |
| | | | | | | Naive Bayes | 0.904 ± 0.026 | **0.977 ± 0.032** | 0.810 ± 0.056 | 0.918 ± 0.025 | all |
| | | | | | | Poly-SVM | 0.948 ± 0.040 | 0.949 ± 0.046 | 0.940 ± 0.055 | 0.954 ± 0.036 | all |
| | | | | | | RBF-SVM | 0.946 ± 0.039 | 0.945 ± 0.050 | 0.946 ± 0.042 | 0.950 ± 0.035 | all |
| | | | | | | Random Forests | **0.965 ± 0.031** | **0.968 ± 0.038** | **0.958 ± 0.047** | **0.969 ± 0.029** | 1637.7 ± 2268.4 |
| | | | | | | CART$_b$ | 0.915 ± 0.052 | 0.905 ± 0.054 | 0.925 ± 0.078 | 0.922 ± 0.049 | 2.0 ± 0.0 |
| | | | | | | CART$_{cv}$ | 0.915 ± 0.054 | 0.918 ± 0.061 | 0.910 ± 0.070 | 0.923 ± 0.049 | 2.2 ± 1.5 |
| | | | | | | SCM$_b$ | 0.922 ± 0.047 | 0.914 ± 0.042 | 0.929 ± 0.078 | 0.929 ± 0.041 | 2.0 ± 0.0 |
| | | | | | | SCM$_{cv}$ | 0.924 ± 0.040 | 0.917 ± 0.050 | 0.923 ± 0.062 | 0.928 ± 0.040 | 3.5 ± 1.1 |
| | ceftazidime | 277 | 249 | 28 | 14.4 | L1-logistic | **0.944 ± 0.034** | 0.990 ± 0.017 | 0.511 ± 0.288 | **0.969 ± 0.018** | 153574.8 ± 295176.4 |
| | | | | | | L2-logistic | 0.927 ± 0.049 | 0.968 ± 0.047 | 0.548 ± 0.310 | 0.960 ± 0.029 | all* |
| | | | | | | Majority | 0.907 ± 0.035 | **1.000 ± 0.000** | 0.000 ± 0.000 | 0.951 ± 0.019 | – |
| | | | | | | Naive Bayes | 0.871 ± 0.039 | 0.858 ± 0.041 | **1.000 ± 0.000** | 0.923 ± 0.024 | all |

Continued on next page

**Table S1.** (Continued)

| Species | Antibiotic | Genomes | Resistant | Susceptible | $k$-mers (millions) | Method | Accuracy | Sensitivity | Specificity | F1 score | Complexity |
|---|---|---|---|---|---|---|---|---|---|---|---|
| | | | | | | Poly-SVM | **0.942 ± 0.032** | 0.982 ± 0.029 | 0.562 ± 0.284 | **0.968 ± 0.017** | all |
| | | | | | | RBF-SVM | **0.942 ± 0.033** | 0.980 ± 0.030 | 0.577 ± 0.289 | **0.968 ± 0.018** | all |
| | | | | | | Random Forests | 0.940 ± 0.021 | 0.982 ± 0.022 | 0.546 ± 0.260 | **0.967 ± 0.012** | 1389.7 ± 2594.2 |
| | | | | | | $CART_b$ | **0.947 ± 0.026** | 0.984 ± 0.022 | 0.593 ± 0.229 | **0.971 ± 0.014** | 1.1 ± 0.3 |
| | | | | | | $CART_{cv}$ | **0.951 ± 0.047** | 0.974 ± 0.042 | 0.736 ± 0.280 | **0.973 ± 0.027** | 2.3 ± 1.3 |
| | | | | | | $SCM_b$ | 0.940 ± 0.043 | 0.976 ± 0.043 | 0.593 ± 0.229 | **0.967 ± 0.024** | 1.2 ± 0.4 |
| | | | | | | $SCM_{cv}$ | 0.935 ± 0.042 | 0.967 ± 0.042 | 0.660 ± 0.277 | **0.964 ± 0.024** | 1.7 ± 1.1 |
| | imipenem | 499 | 325 | 174 | 42.4 | L1-logistic | 0.880 ± 0.029 | 0.915 ± 0.031 | 0.819 ± 0.059 | 0.907 ± 0.025 | 3980.5 ± 4676.0 |
| | | | | | | L2-logistic | **0.885 ± 0.034** | 0.906 ± 0.038 | **0.851 ± 0.070** | **0.909 ± 0.029** | all* |
| | | | | | | Majority | 0.644 ± 0.037 | **1.000 ± 0.000** | 0.000 ± 0.000 | 0.783 ± 0.028 | – |
| | | | | | | Naive Bayes | 0.822 ± 0.027 | 0.912 ± 0.031 | 0.661 ± 0.057 | 0.868 ± 0.023 | all |
| | | | | | | Poly-SVM | **0.886 ± 0.031** | 0.917 ± 0.028 | 0.832 ± 0.072 | **0.912 ± 0.024** | all |
| | | | | | | RBF-SVM | 0.880 ± 0.031 | 0.912 ± 0.041 | 0.824 ± 0.063 | 0.907 ± 0.024 | all |
| | | | | | | Random Forests | **0.892 ± 0.024** | 0.937 ± 0.023 | 0.812 ± 0.052 | **0.917 ± 0.020** | 6314.6 ± 7055.6 |
| | | | | | | $CART_b$ | 0.864 ± 0.042 | 0.915 ± 0.039 | 0.773 ± 0.110 | 0.896 ± 0.032 | 3.4 ± 0.7 |
| | | | | | | $CART_{cv}$ | 0.863 ± 0.041 | 0.910 ± 0.035 | 0.780 ± 0.085 | 0.894 ± 0.033 | 9.6 ± 5.0 |
| | | | | | | $SCM_b$ | 0.849 ± 0.031 | 0.926 ± 0.020 | 0.711 ± 0.078 | 0.888 ± 0.023 | 2.7 ± 0.5 |
| | | | | | | $SCM_{cv}$ | 0.857 ± 0.039 | 0.912 ± 0.046 | 0.759 ± 0.075 | 0.890 ± 0.033 | 10.6 ± 5.2 |
| | meropenem | 236 | 203 | 33 | 13.4 | L1-logistic | 0.896 ± 0.048 | 0.933 ± 0.042 | **0.628 ± 0.208** | 0.940 ± 0.028 | 167820.2 ± 298168.7 |
| | | | | | | L2-logistic | 0.887 ± 0.049 | 0.932 ± 0.046 | 0.560 ± 0.240 | 0.935 ± 0.030 | all* |
| | | | | | | Majority | 0.881 ± 0.025 | **1.000 ± 0.000** | 0.000 ± 0.000 | 0.936 ± 0.014 | – |
| | | | | | | Naive Bayes | 0.791 ± 0.048 | 0.836 ± 0.054 | 0.451 ± 0.269 | 0.876 ± 0.030 | all |
| | | | | | | Poly-SVM | 0.900 ± 0.050 | 0.945 ± 0.019 | 0.580 ± 0.364 | 0.944 ± 0.027 | all |
| | | | | | | RBF-SVM | 0.906 ± 0.042 | 0.954 ± 0.021 | 0.566 ± 0.355 | **0.948 ± 0.022** | all |
| | | | | | | Random Forests | **0.921 ± 0.027** | 0.974 ± 0.017 | 0.541 ± 0.269 | **0.956 ± 0.014** | 2282.4 ± 3017.3 |
| | | | | | | $CART_b$ | 0.874 ± 0.032 | 0.964 ± 0.040 | 0.245 ± 0.334 | 0.931 ± 0.017 | 0.9 ± 0.9 |
| | | | | | | $CART_{cv}$ | 0.900 ± 0.032 | 0.943 ± 0.036 | 0.583 ± 0.220 | 0.943 ± 0.018 | 6.5 ± 2.9 |
| | | | | | | $SCM_b$ | 0.896 ± 0.038 | 0.947 ± 0.029 | 0.499 ± 0.327 | 0.941 ± 0.021 | 1.5 ± 0.5 |
| | | | | | | $SCM_{cv}$ | 0.889 ± 0.036 | 0.943 ± 0.041 | 0.507 ± 0.255 | 0.938 ± 0.020 | 4.5 ± 2.5 |
| | tobramycin | 249 | | 46 | 15.2 | L1-logistic | 0.863 ± 0.019 | 0.905 ± 0.047 | 0.648 ± 0.159 | 0.918 ± 0.012 | 70944.3 ± 159229.9 |
| | | | | | | L2-logistic | 0.857 ± 0.049 | 0.882 ± 0.063 | **0.741 ± 0.140** | 0.912 ± 0.030 | all* |
| | | | | | | Majority | 0.849 ± 0.034 | **1.000 ± 0.000** | 0.000 ± 0.000 | 0.918 ± 0.020 | – |
| | | | | | | Naive Bayes | 0.733 ± 0.049 | 0.784 ± 0.067 | 0.457 ± 0.208 | 0.831 ± 0.038 | all |
| | | | | | | Poly-SVM | **0.873 ± 0.054** | 0.921 ± 0.055 | 0.606 ± 0.226 | **0.925 ± 0.032** | all |
| | | | | | | RBF-SVM | **0.873 ± 0.043** | 0.935 ± 0.051 | 0.520 ± 0.177 | **0.926 ± 0.027** | all |
| | | | | | | Random Forests | **0.882 ± 0.039** | 0.945 ± 0.039 | 0.539 ± 0.183 | **0.931 ± 0.024** | 1034.8 ± 901.9 |
| | | | | | | $CART_b$ | 0.841 ± 0.041 | 0.936 ± 0.050 | 0.342 ± 0.307 | 0.909 ± 0.024 | 1.5 ± 0.7 |
| | | | | | | $CART_{cv}$ | 0.841 ± 0.051 | 0.881 ± 0.059 | 0.627 ± 0.138 | 0.903 ± 0.032 | 6.8 ± 2.3 |
| | | | | | | $SCM_b$ | 0.847 ± 0.040 | 0.924 ± 0.042 | 0.441 ± 0.272 | 0.911 ± 0.023 | 1.7 ± 0.5 |
| | | | | | | $SCM_{cv}$ | 0.841 ± 0.049 | 0.871 ± 0.061 | 0.698 ± 0.187 | 0.902 ± 0.031 | 6.0 ± 2.9 |
| *E. coli* | amoxicillin | 1095 | 661 | 434 | 39.7 | L1-logistic | 0.900 ± 0.029 | 0.874 ± 0.028 | 0.942 ± 0.054 | 0.914 ± 0.026 | 1861.0 ± 4505.5 |
| | | | | | | L2-logistic | 0.888 ± 0.022 | 0.873 ± 0.028 | 0.912 ± 0.040 | 0.905 ± 0.020 | all* |
| | | | | | | Majority | 0.614 ± 0.025 | **1.000 ± 0.000** | 0.000 ± 0.000 | 0.761 ± 0.020 | – |
| | | | | | | Naive Bayes | 0.603 ± 0.025 | 0.552 ± 0.025 | 0.685 ± 0.043 | 0.630 ± 0.027 | all |
| | | | | | | Poly-SVM | 0.869 ± 0.032 | 0.888 ± 0.029 | 0.842 ± 0.058 | 0.893 ± 0.027 | all |

Continued on next page

**Table S1.** (Continued)

| Species | Antibiotic | Genomes | Resistant | Susceptible | *k*-mers (millions) | Method | Accuracy | Sensitivity | Specificity | F1 score | Complexity |
|---|---|---|---|---|---|---|---|---|---|---|---|
| | | | | | | RBF-SVM | 0.864±0.039 | 0.878±0.034 | 0.844±0.054 | 0.888±0.034 | all |
| | | | | | | Random Forests | 0.909±0.022 | 0.893±0.026 | 0.934±0.029 | 0.923±0.019 | 17109.2 ± 13709.9 |
| | | | | | | CART$_b$ | **0.923±0.018** | 0.891±0.026 | **0.973±0.011** | **0.934±0.016** | 3.6 ± 0.5 |
| | | | | | | CART$_{cv}$ | **0.919±0.022** | 0.889±0.024 | **0.966±0.027** | **0.930±0.019** | 4.1 ± 1.7 |
| | | | | | | SCM$_b$ | **0.920±0.016** | 0.893±0.025 | 0.962±0.014 | **0.932±0.015** | 4.1 ± 0.7 |
| | | | | | | SCM$_{cv}$ | **0.920±0.021** | 0.891±0.023 | **0.966±0.028** | **0.932±0.019** | 4.0 ± 1.2 |
| | amoxicillin/clavulan-ic acid | 1524 | 464 | 1060 | 48.5 | L1-logistic | 0.792±0.018 | **0.746±0.075** | 0.812±0.026 | **0.683±0.040** | 3727.2 ± 5890.3 |
| | | | | | | L2-logistic | 0.789±0.022 | 0.684±0.078 | 0.835±0.042 | 0.661±0.035 | all* |
| | | | | | | Majority | 0.697±0.014 | 0.000±0.000 | **1.000±0.000** | – | – |
| | | | | | | Naive Bayes | 0.634±0.026 | 0.596±0.035 | 0.652±0.054 | 0.497±0.017 | all |
| | | | | | | Poly-SVM | 0.779±0.022 | 0.604±0.070 | 0.856±0.020 | 0.622±0.043 | all |
| | | | | | | RBF-SVM | 0.776±0.021 | 0.597±0.073 | 0.855±0.016 | 0.616±0.046 | all |
| | | | | | | Random Forests | 0.812±0.021 | 0.598±0.060 | 0.906±0.023 | 0.657±0.037 | 39289.6 ± 29690.9 |
| | | | | | | CART$_b$ | 0.808±0.021 | 0.563±0.075 | 0.915±0.041 | 0.638±0.040 | 7.0 ± 0.7 |
| | | | | | | CART$_{cv}$ | 0.812±0.019 | 0.533±0.101 | 0.933±0.052 | 0.627±0.047 | 13.3 ± 7.7 |
| | | | | | | SCM$_b$ | 0.818±0.019 | 0.464±0.050 | 0.972±0.014 | 0.606±0.041 | 4.6 ± 1.1 |
| | | | | | | SCM$_{cv}$ | **0.830±0.023** | 0.467±0.059 | 0.988±0.010 | 0.623±0.054 | 6.2 ± 1.9 |
| | ampicillin | 436 | 271 | 165 | 36.1 | L1-logistic | **0.926±0.029** | 0.905±0.052 | **0.964±0.031** | **0.937±0.027** | 3006.9 ± 2011.3 |
| | | | | | | L2-logistic | 0.908±0.038 | 0.900±0.051 | 0.920±0.049 | 0.922±0.034 | all* |
| | | | | | | Majority | 0.610±0.040 | **1.000±0.000** | 0.000±0.000 | 0.757±0.031 | – |
| | | | | | | Naive Bayes | 0.629±0.036 | 0.615±0.058 | 0.651±0.075 | 0.668±0.035 | all |
| | | | | | | Poly-SVM | 0.826±0.029 | 0.839±0.046 | 0.808±0.035 | 0.855±0.025 | all |
| | | | | | | RBF-SVM | 0.824±0.027 | 0.839±0.040 | 0.803±0.042 | 0.853±0.023 | all |
| | | | | | | Random Forests | 0.923±0.042 | 0.913±0.051 | 0.938±0.051 | **0.935±0.038** | 2720.0 ± 6356.9 |
| | | | | | | CART$_b$ | 0.906±0.037 | 0.902±0.044 | 0.910±0.062 | 0.921±0.033 | 2.2 ± 0.6 |
| | | | | | | CART$_{cv}$ | 0.916±0.036 | 0.912±0.042 | 0.922±0.065 | 0.930±0.029 | 3.1 ± 1.7 |
| | | | | | | SCM$_b$ | 0.911±0.040 | 0.912±0.048 | 0.911±0.063 | 0.926±0.035 | 2.2 ± 0.6 |
| | | | | | | SCM$_{cv}$ | **0.933±0.040** | 0.933±0.043 | 0.936±0.065 | **0.944±0.033** | 3.5 ± 1.4 |
| | aztreonam | 419 | 39 | 380 | 35.2 | L1-logistic | **0.954±0.011** | 0.673±0.148 | 0.984±0.010 | 0.725±0.088 | 13896.1 ± 22296.9 |
| | | | | | | L2-logistic | **0.953±0.024** | 0.649±0.173 | 0.985±0.011 | 0.713±0.153 | all* |
| | | | | | | Majority | 0.906±0.018 | 0.000±0.000 | **1.000±0.000** | – | – |
| | | | | | | Naive Bayes | 0.765±0.086 | **0.854±0.135** | 0.755±0.089 | 0.418±0.128 | all |
| | | | | | | Poly-SVM | 0.934±0.024 | 0.497±0.223 | 0.980±0.025 | 0.556±0.207 | all |
| | | | | | | RBF-SVM | 0.931±0.027 | 0.432±0.304 | 0.984±0.019 | – | all |
| | | | | | | Random Forests | **0.959±0.021** | 0.628±0.133 | **0.995±0.009** | 0.745±0.113 | 4960.1 ± 4877.3 |
| | | | | | | CART$_b$ | **0.960±0.021** | 0.696±0.118 | 0.988±0.012 | **0.768±0.105** | 1.0 ± 0.0 |
| | | | | | | CART$_{cv}$ | **0.958±0.023** | 0.696±0.118 | 0.985±0.015 | **0.759±0.108** | 1.6 ± 1.3 |
| | | | | | | SCM$_b$ | **0.960±0.021** | 0.696±0.118 | 0.988±0.012 | **0.768±0.105** | 1.0 ± 0.0 |
| | | | | | | SCM$_{cv}$ | **0.958±0.021** | 0.710±0.110 | 0.984±0.012 | **0.760±0.101** | 1.8 ± 1.0 |
| | cefalotin | 250 | 59 | 191 | 29.7 | L1-logistic | 0.804±0.056 | 0.548±0.149 | 0.884±0.050 | 0.571±0.124 | 592.7 ± 753.2 |
| | | | | | | L2-logistic | 0.812±0.074 | **0.565±0.157** | 0.893±0.058 | 0.594±0.150 | all* |
| | | | | | | Majority | 0.752±0.056 | 0.000±0.000 | **1.000±0.000** | – | – |
| | | | | | | Naive Bayes | 0.834±0.045 | **0.558±0.156** | 0.919±0.046 | 0.613±0.126 | all |
| | | | | | | Poly-SVM | **0.840±0.037** | 0.485±0.114 | 0.955±0.025 | 0.589±0.118 | all |

Continued on next page

**Table S1.** (Continued)

| Species | Antibiotic | Genomes | Resistant | Susceptible | $k$-mers (millions) | Method | Accuracy | Sensitivity | Specificity | F1 score | Complexity |
|---|---|---|---|---|---|---|---|---|---|---|---|
| | | | | | | RBF-SVM | 0.836±0.030 | 0.492±0.112 | 0.947±0.028 | 0.586±0.111 | all |
| | | | | | | Random Forests | **0.846±0.042** | 0.502±0.138 | 0.956±0.035 | 0.604±0.145 | 1334.6 ± 2911.6 |
| | | | | | | $CART_b$ | 0.836±0.048 | 0.530±0.131 | 0.933±0.041 | 0.608±0.128 | 1.0 ± 0.0 |
| | | | | | | $CART_{cv}$ | 0.836±0.044 | 0.524±0.120 | 0.936±0.041 | 0.601±0.124 | 2.0 ± 1.5 |
| | | | | | | $SCM_b$ | 0.830±0.047 | 0.520±0.137 | 0.927±0.034 | 0.594±0.130 | 1.0 ± 0.0 |
| | | | | | | $SCM_{cv}$ | **0.840±0.057** | 0.538±0.092 | 0.941±0.045 | **0.626±0.106** | 3.2 ± 1.4 |
| | cefepime | 426 | 32 | 394 | 35.8 | L1-logistic | **0.971±0.022** | 0.825±0.163 | 0.981±0.021 | 0.795±0.133 | 3594.5 ± 4097.0 |
| | | | | | | L2-logistic | **0.975±0.019** | 0.754±0.215 | 0.990±0.013 | 0.786±0.205 | all* |
| | | | | | | Majority | 0.934±0.019 | 0.000±0.000 | **1.000±0.000** | – | – |
| | | | | | | Naive Bayes | 0.782±0.049 | 0.679±0.197 | 0.790±0.052 | 0.293±0.115 | all |
| | | | | | | Poly-SVM | 0.966±0.015 | 0.576±0.259 | **0.994±0.009** | 0.657±0.188 | all |
| | | | | | | RBF-SVM | 0.966±0.016 | 0.565±0.258 | **0.995±0.009** | 0.657±0.186 | all |
| | | | | | | Random Forests | **0.979±0.013** | 0.754±0.173 | **0.995±0.011** | **0.823±0.103** | 1626.6 ± 2320.9 |
| | | | | | | $CART_b$ | 0.966±0.016 | 0.728±0.144 | 0.982±0.017 | 0.736±0.083 | 1.4 ± 0.7 |
| | | | | | | $CART_{cv}$ | **0.973±0.014** | **0.841±0.153** | 0.982±0.012 | 0.800±0.100 | 3.4 ± 1.2 |
| | | | | | | $SCM_b$ | 0.964±0.017 | 0.705±0.199 | 0.981±0.017 | 0.709±0.130 | 1.5 ± 0.5 |
| | | | | | | $SCM_{cv}$ | 0.965±0.018 | 0.768±0.299 | 0.978±0.018 | – | 2.1 ± 0.3 |
| | cefotaxime | 1450 | 139 | 1311 | 43.7 | L1-logistic | **0.976±0.009** | **0.860±0.088** | 0.988±0.006 | 0.873±0.060 | 67531.0 ± 138221.3 |
| | | | | | | L2-logistic | **0.973±0.010** | **0.860±0.086** | 0.985±0.006 | 0.859±0.062 | all* |
| | | | | | | Majority | 0.898±0.018 | 0.000±0.000 | **1.000±0.000** | – | – |
| | | | | | | Naive Bayes | 0.830±0.048 | 0.754±0.105 | 0.837±0.061 | 0.477±0.060 | all |
| | | | | | | Poly-SVM | **0.971±0.011** | 0.816±0.101 | 0.988±0.006 | 0.846±0.067 | all |
| | | | | | | RBF-SVM | 0.969±0.010 | 0.803±0.091 | 0.987±0.007 | 0.833±0.065 | all |
| | | | | | | Random Forests | **0.979±0.011** | 0.850±0.089 | **0.993±0.005** | **0.890±0.059** | 12400.7 ± 11836.2 |
| | | | | | | $CART_b$ | **0.973±0.009** | 0.782±0.089 | **0.994±0.004** | 0.849±0.058 | 2.8 ± 0.9 |
| | | | | | | $CART_{cv}$ | **0.979±0.008** | **0.858±0.092** | **0.993±0.003** | **0.890±0.051** | 6.0 ± 2.4 |
| | | | | | | $SCM_b$ | **0.974±0.007** | 0.786±0.074 | **0.995±0.004** | 0.858±0.044 | 3.0 ± 1.1 |
| | | | | | | $SCM_{cv}$ | **0.980±0.009** | **0.860±0.094** | **0.993±0.003** | **0.892±0.053** | 5.2 ± 1.2 |
| | cefoxitin | 417 | 27 | 390 | 35.3 | L1-logistic | 0.964±0.025 | 0.682±0.232 | 0.982±0.017 | 0.686±0.198 | 1584.0 ± 3147.2 |
| | | | | | | L2-logistic | 0.945±0.021 | 0.489±0.219 | 0.974±0.018 | 0.500±0.175 | all* |
| | | | | | | Majority | 0.940±0.015 | 0.000±0.000 | **1.000±0.000** | – | – |
| | | | | | | Naive Bayes | 0.810±0.038 | 0.381±0.234 | 0.836±0.036 | – | all |
| | | | | | | Poly-SVM | 0.965±0.012 | 0.400±0.258 | **0.999±0.004** | – | all |
| | | | | | | RBF-SVM | 0.967±0.013 | 0.450±0.247 | **0.999±0.004** | – | all |
| | | | | | | Random Forests | 0.961±0.017 | 0.407±0.274 | **0.996±0.009** | – | 788.2 ± 1579.9 |
| | | | | | | $CART_b$ | **0.973±0.018** | **0.794±0.130** | 0.985±0.013 | 0.782±0.131 | 1.0 ± 0.0 |
| | | | | | | $CART_{cv}$ | 0.971±0.017 | **0.799±0.129** | 0.982±0.015 | 0.769±0.123 | 1.7 ± 0.9 |
| | | | | | | $SCM_b$ | **0.973±0.018** | **0.794±0.130** | 0.985±0.013 | 0.782±0.131 | 1.0 ± 0.0 |
| | | | | | | $SCM_{cv}$ | **0.982±0.014** | 0.744±0.182 | **0.996±0.012** | **0.816±0.143** | 2.0 ± 0.0 |
| | cefuroxime | 1507 | 241 | 1266 | 47.9 | L1-logistic | 0.833±0.021 | 0.333±0.289 | 0.928±0.065 | – | 11536.9 ± 14007.2 |
| | | | | | | L2-logistic | 0.830±0.028 | 0.504±0.111 | 0.893±0.036 | 0.486±0.085 | all* |
| | | | | | | Majority | 0.838±0.020 | 0.000±0.000 | **1.000±0.000** | – | – |
| | | | | | | Naive Bayes | 0.793±0.022 | **0.578±0.096** | 0.834±0.031 | 0.472±0.054 | all |
| | | | | | | Poly-SVM | 0.880±0.018 | 0.395±0.073 | 0.974±0.013 | 0.514±0.083 | all |
| | | | | | | RBF-SVM | 0.875±0.011 | 0.392±0.062 | 0.968±0.012 | 0.500±0.059 | all |

Continued on next page

**Table S1.** (Continued)

| Species | Antibiotic | Genomes | Resistant | Susceptible | *k*-mers (millions) | Method | Accuracy | Sensitivity | Specificity | F1 score | Complexity |
|---|---|---|---|---|---|---|---|---|---|---|---|
| | | | | | | Random Forests | 0.904 ± 0.012 | 0.439 ± 0.063 | **0.994 ± 0.005** | 0.594 ± 0.056 | 8889.5 ± 13173.8 |
| | | | | | | CART$_b$ | **0.913 ± 0.015** | 0.502 ± 0.055 | **0.992 ± 0.004** | 0.650 ± 0.053 | 3.0 ± 0.5 |
| | | | | | | CART$_{cv}$ | **0.916 ± 0.011** | 0.534 ± 0.058 | 0.990 ± 0.006 | **0.672 ± 0.047** | 5.5 ± 1.4 |
| | | | | | | SCM$_b$ | **0.915 ± 0.015** | 0.515 ± 0.052 | **0.992 ± 0.004** | 0.660 ± 0.051 | 3.4 ± 0.8 |
| | | | | | | SCM$_{cv}$ | **0.917 ± 0.012** | 0.545 ± 0.064 | 0.989 ± 0.004 | **0.679 ± 0.052** | 5.5 ± 1.2 |
| | ceftazidime | 1497 | 99 | 1398 | 48.8 | L1-logistic | 0.976 ± 0.010 | 0.794 ± 0.136 | 0.989 ± 0.007 | 0.812 ± 0.093 | 47016.4 ± 110544.7 |
| | | | | | | L2-logistic | 0.974 ± 0.008 | 0.743 ± 0.126 | 0.990 ± 0.008 | 0.786 ± 0.087 | all* |
| | | | | | | Majority | 0.932 ± 0.010 | 0.000 ± 0.000 | **1.000 ± 0.000** | – | – |
| | | | | | | Naive Bayes | 0.768 ± 0.023 | **0.905 ± 0.114** | 0.758 ± 0.025 | 0.343 ± 0.047 | all |
| | | | | | | Poly-SVM | 0.966 ± 0.008 | 0.636 ± 0.097 | 0.990 ± 0.009 | 0.710 ± 0.080 | all |
| | | | | | | RBF-SVM | 0.966 ± 0.008 | 0.639 ± 0.088 | 0.989 ± 0.008 | 0.712 ± 0.074 | all |
| | | | | | | Random Forests | **0.988 ± 0.007** | 0.840 ± 0.108 | **0.998 ± 0.002** | **0.895 ± 0.064** | 7770.6 ± 8004.7 |
| | | | | | | CART$_b$ | 0.976 ± 0.010 | 0.723 ± 0.140 | **0.994 ± 0.004** | 0.789 ± 0.105 | 2.8 ± 0.4 |
| | | | | | | CART$_{cv}$ | **0.983 ± 0.005** | 0.826 ± 0.082 | **0.994 ± 0.006** | 0.867 ± 0.045 | 5.8 ± 1.9 |
| | | | | | | SCM$_b$ | **0.979 ± 0.010** | 0.730 ± 0.128 | **0.996 ± 0.004** | 0.814 ± 0.101 | 2.8 ± 0.6 |
| | | | | | | SCM$_{cv}$ | **0.985 ± 0.005** | 0.830 ± 0.065 | **0.996 ± 0.005** | 0.882 ± 0.043 | 3.8 ± 0.6 |
| | ciprofloxacin | 1519 | 289 | 1230 | 44.7 | L1-logistic | **0.986 ± 0.006** | **0.949 ± 0.028** | **0.995 ± 0.005** | **0.962 ± 0.018** | 432.7 ± 1043.6 |
| | | | | | | L2-logistic | 0.963 ± 0.012 | 0.868 ± 0.065 | 0.986 ± 0.010 | 0.900 ± 0.031 | all* |
| | | | | | | Majority | 0.806 ± 0.021 | 0.000 ± 0.000 | **1.000 ± 0.000** | – | – |
| | | | | | | Naive Bayes | 0.835 ± 0.029 | 0.911 ± 0.052 | 0.817 ± 0.043 | 0.682 ± 0.048 | all |
| | | | | | | Poly-SVM | 0.965 ± 0.014 | 0.845 ± 0.061 | **0.994 ± 0.006** | 0.902 ± 0.035 | all |
| | | | | | | RBF-SVM | 0.965 ± 0.014 | 0.850 ± 0.060 | **0.993 ± 0.008** | 0.903 ± 0.034 | all |
| | | | | | | Random Forests | 0.975 ± 0.011 | 0.892 ± 0.051 | **0.996 ± 0.006** | 0.932 ± 0.029 | 6291.3 ± 11483.4 |
| | | | | | | CART$_b$ | **0.985 ± 0.005** | 0.938 ± 0.022 | **0.996 ± 0.005** | **0.960 ± 0.013** | 2.0 ± 0.0 |
| | | | | | | CART$_{cv}$ | **0.983 ± 0.005** | 0.935 ± 0.022 | **0.996 ± 0.005** | **0.957 ± 0.012** | 2.2 ± 0.4 |
| | | | | | | SCM$_b$ | **0.983 ± 0.005** | 0.938 ± 0.022 | **0.995 ± 0.005** | **0.956 ± 0.013** | 2.0 ± 0.0 |
| | | | | | | SCM$_{cv}$ | **0.983 ± 0.004** | 0.939 ± 0.023 | **0.995 ± 0.005** | **0.957 ± 0.010** | 2.7 ± 1.1 |
| | gentamicin | 1513 | 115 | 1398 | 48.7 | L1-logistic | **0.983 ± 0.007** | **0.896 ± 0.053** | **0.991 ± 0.005** | 0.891 ± 0.043 | 5673.3 ± 13968.4 |
| | | | | | | L2-logistic | **0.979 ± 0.007** | 0.851 ± 0.068 | **0.991 ± 0.005** | 0.864 ± 0.045 | all* |
| | | | | | | Majority | 0.923 ± 0.011 | 0.000 ± 0.000 | **1.000 ± 0.000** | – | – |
| | | | | | | Naive Bayes | 0.687 ± 0.037 | 0.824 ± 0.048 | 0.676 ± 0.043 | 0.289 ± 0.038 | all |
| | | | | | | Poly-SVM | 0.956 ± 0.014 | 0.614 ± 0.152 | 0.986 ± 0.006 | 0.676 ± 0.104 | all |
| | | | | | | RBF-SVM | 0.957 ± 0.013 | 0.619 ± 0.134 | 0.986 ± 0.005 | 0.686 ± 0.096 | all |
| | | | | | | Random Forests | **0.987 ± 0.007** | **0.900 ± 0.070** | **0.995 ± 0.005** | **0.914 ± 0.055** | 4242.2 ± 5881.0 |
| | | | | | | CART$_b$ | **0.986 ± 0.006** | **0.898 ± 0.061** | **0.994 ± 0.004** | **0.907 ± 0.046** | 2.0 ± 0.0 |
| | | | | | | CART$_{cv}$ | **0.985 ± 0.009** | **0.893 ± 0.064** | **0.993 ± 0.007** | 0.898 ± 0.063 | 2.4 ± 1.3 |
| | | | | | | SCM$_b$ | **0.986 ± 0.006** | **0.898 ± 0.061** | **0.994 ± 0.004** | **0.907 ± 0.046** | 2.0 ± 0.0 |
| | | | | | | SCM$_{cv}$ | **0.986 ± 0.007** | **0.898 ± 0.061** | **0.993 ± 0.005** | **0.905 ± 0.049** | 2.2 ± 0.6 |
| | meropenem | 446 | 28 | 418 | 36.2 | L1-logistic | **0.982 ± 0.013** | **0.963 ± 0.078** | 0.983 ± 0.015 | 0.844 ± 0.093 | 445.5 ± 394.3 |
| | | | | | | L2-logistic | 0.976 ± 0.011 | 0.787 ± 0.189 | 0.987 ± 0.009 | 0.761 ± 0.118 | all* |
| | | | | | | Majority | 0.949 ± 0.012 | 0.000 ± 0.000 | **1.000 ± 0.000** | – | – |
| | | | | | | Naive Bayes | 0.811 ± 0.046 | 0.838 ± 0.121 | 0.809 ± 0.047 | 0.316 ± 0.101 | all |
| | | | | | | Poly-SVM | **0.984 ± 0.009** | 0.770 ± 0.184 | **0.996 ± 0.006** | 0.823 ± 0.098 | all |
| | | | | | | RBF-SVM | **0.983 ± 0.010** | 0.745 ± 0.203 | **0.996 ± 0.008** | 0.806 ± 0.105 | all |
| | | | | | | Random Forests | **0.990 ± 0.006** | 0.863 ± 0.127 | **0.996 ± 0.006** | **0.883 ± 0.093** | 2669.0 ± 2582.8 |

Continued on next page

**Table S1.** (Continued)

| Species | Antibiotic | Genomes | Resistant | Susceptible | $k$-mers (millions) | Method | Accuracy | Sensitivity | Specificity | F1 score | Complexity |
|---|---|---|---|---|---|---|---|---|---|---|---|
| | | | | | | CART$_b$ | 0.978 ± 0.013 | 0.922 ± 0.130 | 0.981 ± 0.014 | 0.802 ± 0.097 | 1.4 ± 0.5 |
| | | | | | | CART$_{cv}$ | 0.976 ± 0.013 | 0.907 ± 0.158 | 0.981 ± 0.014 | 0.791 ± 0.103 | 1.6 ± 1.1 |
| | | | | | | SCM$_b$ | 0.978 ± 0.013 | 0.922 ± 0.130 | 0.981 ± 0.014 | 0.802 ± 0.097 | 1.1 ± 0.3 |
| | | | | | | SCM$_{cv}$ | 0.973 ± 0.012 | 0.840 ± 0.152 | 0.981 ± 0.016 | 0.758 ± 0.069 | 1.8 ± 0.8 |
| | piperacillin/tazobactam | 1461 | 99 | 1362 | 48.0 | L1-logistic | **0.929 ± 0.012** | 0.000 ± 0.000 | **1.000 ± 0.000** | – | 0.0 ± 0.0 |
| | | | | | | L2-logistic | 0.899 ± 0.019 | 0.176 ± 0.101 | 0.953 ± 0.016 | – | all* |
| | | | | | | Majority | **0.929 ± 0.012** | 0.000 ± 0.000 | **1.000 ± 0.000** | – | – |
| | | | | | | Naive Bayes | 0.675 ± 0.034 | **0.481 ± 0.139** | 0.689 ± 0.038 | **0.173 ± 0.053** | all |
| | | | | | | Poly-SVM | **0.933 ± 0.010** | 0.066 ± 0.053 | **0.999 ± 0.002** | – | all |
| | | | | | | RBF-SVM | **0.931 ± 0.010** | 0.054 ± 0.045 | **0.998 ± 0.003** | – | all |
| | | | | | | Random Forests | **0.935 ± 0.010** | 0.100 ± 0.060 | **0.998 ± 0.004** | – | 1164.6 ± 2422.5 |
| | | | | | | CART$_b$ | **0.929 ± 0.010** | 0.036 ± 0.049 | **0.997 ± 0.004** | – | 0.5 ± 0.5 |
| | | | | | | CART$_{cv}$ | **0.930 ± 0.012** | 0.104 ± 0.060 | **0.993 ± 0.006** | – | 1.4 ± 0.7 |
| | | | | | | SCM$_b$ | **0.932 ± 0.012** | 0.095 ± 0.047 | **0.995 ± 0.006** | – | 1.0 ± 0.0 |
| | | | | | | SCM$_{cv}$ | **0.933 ± 0.010** | 0.099 ± 0.052 | **0.996 ± 0.003** | – | 1.8 ± 0.4 |
| | tobramycin | 422 | 50 | 372 | 31.5 | L1-logistic | 0.974 ± 0.014 | 0.892 ± 0.115 | 0.985 ± 0.015 | 0.883 ± 0.062 | 1686.7 ± 1223.2 |
| | | | | | | L2-logistic | 0.964 ± 0.018 | 0.824 ± 0.124 | 0.982 ± 0.018 | 0.840 ± 0.082 | all* |
| | | | | | | Majority | 0.886 ± 0.025 | 0.000 ± 0.000 | **1.000 ± 0.000** | – | – |
| | | | | | | Naive Bayes | 0.790 ± 0.074 | 0.802 ± 0.166 | 0.787 ± 0.097 | 0.473 ± 0.074 | all |
| | | | | | | Poly-SVM | 0.915 ± 0.027 | 0.536 ± 0.088 | 0.965 ± 0.022 | 0.593 ± 0.100 | all |
| | | | | | | RBF-SVM | 0.912 ± 0.033 | 0.546 ± 0.117 | 0.960 ± 0.029 | 0.589 ± 0.124 | all |
| | | | | | | Random Forests | **0.981 ± 0.018** | 0.867 ± 0.135 | **0.996 ± 0.006** | 0.908 ± 0.095 | 574.1 ± 917.1 |
| | | | | | | CART$_b$ | **0.985 ± 0.014** | **0.924 ± 0.106** | **0.992 ± 0.009** | **0.927 ± 0.072** | 2.0 ± 0.0 |
| | | | | | | CART$_{cv}$ | **0.985 ± 0.014** | **0.924 ± 0.106** | **0.992 ± 0.009** | **0.927 ± 0.072** | 2.0 ± 0.0 |
| | | | | | | SCM$_b$ | **0.985 ± 0.014** | **0.924 ± 0.106** | **0.992 ± 0.009** | **0.927 ± 0.072** | 2.0 ± 0.0 |
| | | | | | | SCM$_{cv}$ | **0.981 ± 0.014** | 0.899 ± 0.115 | **0.992 ± 0.009** | 0.913 ± 0.070 | 2.0 ± 0.0 |
| | trimethoprim | 411 | 147 | 264 | 34.9 | L1-logistic | **0.933 ± 0.025** | **0.904 ± 0.060** | 0.949 ± 0.033 | **0.900 ± 0.036** | 14801.2 ± 14384.9 |
| | | | | | | L2-logistic | 0.911 ± 0.026 | 0.890 ± 0.059 | 0.923 ± 0.037 | 0.870 ± 0.039 | all* |
| | | | | | | Majority | 0.662 ± 0.038 | 0.000 ± 0.000 | **1.000 ± 0.000** | – | – |
| | | | | | | Naive Bayes | 0.711 ± 0.038 | 0.730 ± 0.085 | 0.700 ± 0.050 | 0.627 ± 0.064 | all |
| | | | | | | Poly-SVM | 0.839 ± 0.044 | 0.753 ± 0.120 | 0.885 ± 0.049 | 0.755 ± 0.075 | all |
| | | | | | | RBF-SVM | 0.832 ± 0.050 | 0.746 ± 0.124 | 0.877 ± 0.042 | 0.745 ± 0.081 | all |
| | | | | | | Random Forests | **0.929 ± 0.021** | **0.898 ± 0.057** | 0.946 ± 0.027 | 0.893 ± 0.037 | 7838.8 ± 8352.5 |
| | | | | | | CART$_b$ | **0.935 ± 0.021** | 0.880 ± 0.058 | 0.963 ± 0.031 | **0.901 ± 0.033** | 2.0 ± 0.0 |
| | | | | | | CART$_{cv}$ | 0.926 ± 0.032 | 0.870 ± 0.072 | 0.955 ± 0.052 | 0.887 ± 0.046 | 2.8 ± 0.9 |
| | | | | | | SCM$_b$ | **0.938 ± 0.022** | 0.887 ± 0.061 | 0.963 ± 0.031 | **0.905 ± 0.035** | 2.0 ± 0.0 |
| | | | | | | SCM$_{cv}$ | **0.935 ± 0.024** | 0.887 ± 0.054 | 0.960 ± 0.033 | **0.901 ± 0.039** | 2.8 ± 1.4 |
| *E. faecium* | vancomycin | 134 | 51 | 83 | 10.3 | L1-logistic | **1.000 ± 0.000** | **1.000 ± 0.000** | **1.000 ± 0.000** | **1.000 ± 0.000** | 142.0 ± 45.2 |
| | | | | | | L2-logistic | **1.000 ± 0.000** | **1.000 ± 0.000** | **1.000 ± 0.000** | **1.000 ± 0.000** | all* |
| | | | | | | Majority | 0.588 ± 0.112 | 0.000 ± 0.000 | **1.000 ± 0.000** | – | – |
| | | | | | | Naive Bayes | 0.808 ± 0.110 | 0.589 ± 0.189 | 0.976 ± 0.043 | 0.707 ± 0.159 | all |
| | | | | | | Poly-SVM | **0.996 ± 0.012** | **0.992 ± 0.024** | **1.000 ± 0.000** | **0.996 ± 0.013** | all |
| | | | | | | RBF-SVM | **0.992 ± 0.016** | 0.980 ± 0.044 | **1.000 ± 0.000** | 0.989 ± 0.023 | all |
| | | | | | | Random Forests | **1.000 ± 0.000** | **1.000 ± 0.000** | **1.000 ± 0.000** | **1.000 ± 0.000** | 202.6 ± 491.7 |

Continued on next page

**Table S1.** (Continued)

| Species | Antibiotic | Genomes | Resistant | Susceptible | $k$-mers (millions) | Method | Accuracy | Sensitivity | Specificity | F1 score | Complexity |
|---|---|---|---|---|---|---|---|---|---|---|---|
| | | | | | | CART$_b$ | **1.000 ± 0.000** | **1.000 ± 0.000** | **1.000 ± 0.000** | **1.000 ± 0.000** | 1.0 ± 0.0 |
| | | | | | | CART$_{cv}$ | **1.000 ± 0.000** | **1.000 ± 0.000** | **1.000 ± 0.000** | **1.000 ± 0.000** | 1.0 ± 0.0 |
| | | | | | | SCM$_b$ | **1.000 ± 0.000** | **1.000 ± 0.000** | **1.000 ± 0.000** | **1.000 ± 0.000** | 1.0 ± 0.0 |
| | | | | | | SCM$_{cv}$ | **1.000 ± 0.000** | **1.000 ± 0.000** | **1.000 ± 0.000** | **1.000 ± 0.000** | 1.0 ± 0.0 |
| *K. pneumoniae* | amikacin | 1893 | 180 | 1713 | 73.2 | L1-logistic | **0.951 ± 0.010** | 0.740 ± 0.060 | 0.974 ± 0.008 | 0.744 ± 0.038 | 37210.7 ± 42390.4 |
| | | | | | | L2-logistic | 0.942 ± 0.010 | 0.694 ± 0.068 | 0.968 ± 0.010 | 0.693 ± 0.051 | all* |
| | | | | | | Majority | 0.904 ± 0.014 | 0.000 ± 0.000 | **1.000 ± 0.000** | – | – |
| | | | | | | Naive Bayes | 0.875 ± 0.020 | **0.910 ± 0.046** | 0.872 ± 0.020 | 0.583 ± 0.057 | all |
| | | | | | | Poly-SVM | **0.958 ± 0.010** | 0.762 ± 0.086 | 0.980 ± 0.006 | **0.776 ± 0.050** | all |
| | | | | | | RBF-SVM | **0.957 ± 0.010** | 0.764 ± 0.075 | 0.978 ± 0.005 | **0.773 ± 0.042** | all |
| | | | | | | Random Forests | **0.954 ± 0.013** | 0.706 ± 0.076 | 0.981 ± 0.008 | 0.750 ± 0.055 | 10355.2 ± 16634.8 |
| | | | | | | CART$_b$ | **0.949 ± 0.011** | 0.673 ± 0.086 | 0.978 ± 0.007 | 0.715 ± 0.061 | 5.2 ± 1.5 |
| | | | | | | CART$_{cv}$ | **0.951 ± 0.015** | 0.699 ± 0.127 | 0.977 ± 0.009 | 0.726 ± 0.096 | 11.1 ± 4.0 |
| | | | | | | SCM$_b$ | **0.949 ± 0.015** | 0.643 ± 0.140 | 0.981 ± 0.005 | 0.698 ± 0.120 | 4.2 ± 0.4 |
| | | | | | | SCM$_{cv}$ | **0.952 ± 0.011** | 0.642 ± 0.062 | 0.985 ± 0.008 | 0.720 ± 0.058 | 11.3 ± 5.2 |
| | amoxicillin/clavulan-ic acid | 236 | 120 | 116 | 37.2 | L1-logistic | **0.921 ± 0.020** | 0.937 ± 0.042 | **0.904 ± 0.059** | **0.926 ± 0.021** | 69783.1 ± 171583.6 |
| | | | | | | L2-logistic | **0.926 ± 0.037** | **0.953 ± 0.043** | **0.898 ± 0.061** | **0.930 ± 0.038** | all* |
| | | | | | | Majority | 0.457 ± 0.067 | 0.600 ± 0.516 | 0.400 ± 0.516 | – | – |
| | | | | | | Naive Bayes | 0.653 ± 0.100 | 0.932 ± 0.045 | 0.343 ± 0.174 | 0.740 ± 0.080 | all |
| | | | | | | Poly-SVM | 0.885 ± 0.040 | 0.884 ± 0.068 | 0.894 ± 0.054 | 0.890 ± 0.042 | all |
| | | | | | | RBF-SVM | 0.891 ± 0.052 | 0.888 ± 0.071 | **0.901 ± 0.053** | 0.896 ± 0.048 | all |
| | | | | | | Random Forests | 0.911 ± 0.026 | 0.937 ± 0.052 | 0.880 ± 0.065 | 0.916 ± 0.028 | 2255.7 ± 3172.6 |
| | | | | | | CART$_b$ | 0.883 ± 0.046 | 0.904 ± 0.084 | 0.855 ± 0.101 | 0.890 ± 0.050 | 1.8 ± 0.6 |
| | | | | | | CART$_{cv}$ | 0.872 ± 0.040 | 0.883 ± 0.074 | 0.863 ± 0.082 | 0.879 ± 0.043 | 2.2 ± 1.1 |
| | | | | | | SCM$_b$ | 0.891 ± 0.039 | 0.929 ± 0.069 | 0.850 ± 0.097 | 0.902 ± 0.029 | 1.9 ± 0.6 |
| | | | | | | SCM$_{cv}$ | 0.872 ± 0.043 | 0.908 ± 0.075 | 0.833 ± 0.101 | 0.883 ± 0.038 | 2.2 ± 0.9 |
| | ampicillin/sulbactam | 1654 | 1563 | 91 | 52.3 | L1-logistic | **0.982 ± 0.006** | 0.989 ± 0.005 | 0.843 ± 0.116 | **0.990 ± 0.003** | 2168.7 ± 787.6 |
| | | | | | | L2-logistic | 0.970 ± 0.010 | 0.984 ± 0.006 | 0.709 ± 0.202 | 0.984 ± 0.005 | all* |
| | | | | | | Majority | 0.952 ± 0.011 | **1.000 ± 0.000** | 0.000 ± 0.000 | 0.975 ± 0.006 | – |
| | | | | | | Naive Bayes | 0.802 ± 0.019 | 0.794 ± 0.021 | **0.955 ± 0.056** | 0.884 ± 0.012 | all |
| | | | | | | Poly-SVM | 0.974 ± 0.009 | 0.989 ± 0.005 | 0.695 ± 0.128 | **0.986 ± 0.005** | all |
| | | | | | | RBF-SVM | 0.975 ± 0.008 | 0.988 ± 0.005 | 0.706 ± 0.117 | **0.987 ± 0.004** | all |
| | | | | | | Random Forests | **0.984 ± 0.009** | **0.996 ± 0.005** | 0.748 ± 0.159 | **0.991 ± 0.005** | 3295.4 ± 5665.1 |
| | | | | | | CART$_b$ | **0.988 ± 0.008** | **0.997 ± 0.004** | 0.810 ± 0.117 | **0.994 ± 0.004** | 3.0 ± 0.0 |
| | | | | | | CART$_{cv}$ | **0.983 ± 0.006** | **0.991 ± 0.007** | 0.824 ± 0.109 | **0.991 ± 0.003** | 4.5 ± 3.3 |
| | | | | | | SCM$_b$ | **0.986 ± 0.008** | **0.994 ± 0.006** | 0.829 ± 0.110 | **0.993 ± 0.004** | 3.0 ± 0.5 |
| | | | | | | SCM$_{cv}$ | **0.985 ± 0.008** | **0.993 ± 0.007** | 0.829 ± 0.110 | **0.992 ± 0.004** | 3.3 ± 0.8 |
| | aztreonam | 1805 | 1582 | 223 | 61.1 | L1-logistic | 0.853 ± 0.021 | 0.908 ± 0.015 | 0.464 ± 0.086 | 0.915 ± 0.013 | 12926.6 ± 2079.5 |
| | | | | | | L2-logistic | 0.884 ± 0.014 | 0.953 ± 0.016 | 0.394 ± 0.093 | 0.935 ± 0.009 | all* |
| | | | | | | Majority | 0.876 ± 0.020 | **1.000 ± 0.000** | 0.000 ± 0.000 | 0.934 ± 0.011 | – |
| | | | | | | Naive Bayes | 0.693 ± 0.029 | 0.692 ± 0.028 | **0.703 ± 0.083** | 0.798 ± 0.020 | all |
| | | | | | | Poly-SVM | 0.904 ± 0.016 | 0.971 ± 0.012 | 0.431 ± 0.065 | **0.947 ± 0.009** | all |
| | | | | | | RBF-SVM | **0.908 ± 0.014** | 0.973 ± 0.010 | 0.446 ± 0.050 | **0.949 ± 0.009** | all |
| | | | | | | Random Forests | **0.916 ± 0.013** | 0.982 ± 0.008 | 0.452 ± 0.060 | **0.953 ± 0.008** | 28793.8 ± 20797.3 |

Continued on next page

**Table S1.** (Continued)

| Species | Antibiotic | Genomes | Resistant | Susceptible | $k$-mers (millions) | Method | Accuracy | Sensitivity | Specificity | F1 score | Complexity |
|---|---|---|---|---|---|---|---|---|---|---|---|
| | | | | | | $CART_b$ | **0.907±0.016** | 0.977±0.016 | 0.407±0.090 | **0.948±0.010** | 5.0 ± 0.7 |
| | | | | | | $CART_{cv}$ | 0.901±0.013 | 0.966±0.014 | 0.434±0.106 | **0.944±0.008** | 7.4 ± 3.3 |
| | | | | | | $SCM_b$ | **0.914±0.013** | 0.971±0.007 | 0.510±0.074 | **0.952±0.008** | 6.4 ± 1.4 |
| | | | | | | $SCM_{cv}$ | **0.911±0.011** | 0.980±0.006 | 0.418±0.082 | **0.951±0.007** | 12.4 ± 3.2 |
| | cefazolin | 1895 | 1706 | 189 | 65.6 | L1-logistic | 0.941±0.014 | 0.967±0.012 | 0.705±0.069 | 0.967±0.008 | 7038.1 ± 5605.9 |
| | | | | | | L2-logistic | 0.939±0.016 | 0.970±0.013 | 0.667±0.097 | 0.966±0.009 | all* |
| | | | | | | Majority | 0.901±0.014 | **1.000±0.000** | 0.000±0.000 | 0.948±0.008 | – |
| | | | | | | Naive Bayes | 0.890±0.026 | 0.908±0.034 | 0.727±0.104 | 0.937±0.017 | all |
| | | | | | | Poly-SVM | **0.967±0.010** | 0.979±0.008 | **0.852±0.043** | **0.982±0.006** | all |
| | | | | | | RBF-SVM | **0.967±0.010** | 0.980±0.009 | **0.852±0.038** | **0.982±0.006** | all |
| | | | | | | Random Forests | 0.944±0.014 | 0.975±0.010 | 0.657±0.091 | 0.969±0.008 | 22077.8 ± 15353.5 |
| | | | | | | $CART_b$ | **0.963±0.010** | 0.977±0.008 | **0.846±0.071** | **0.980±0.006** | 5.1 ± 0.6 |
| | | | | | | $CART_{cv}$ | 0.960±0.010 | 0.975±0.008 | 0.831±0.098 | **0.978±0.005** | 8.1 ± 3.4 |
| | | | | | | $SCM_b$ | **0.970±0.006** | 0.983±0.008 | **0.853±0.061** | **0.984±0.004** | 6.5 ± 1.0 |
| | | | | | | $SCM_{cv}$ | **0.969±0.005** | 0.983±0.008 | 0.835±0.060 | **0.983±0.003** | 7.1 ± 1.4 |
| | cefepime | 1650 | 1098 | 552 | 65.3 | L1-logistic | 0.766±0.017 | 0.778±0.025 | 0.742±0.047 | 0.817±0.015 | 5579.4 ± 9607.8 |
| | | | | | | L2-logistic | 0.776±0.023 | 0.785±0.032 | **0.758±0.035** | 0.825±0.017 | all* |
| | | | | | | Majority | 0.672±0.023 | **1.000±0.000** | 0.000±0.000 | 0.804±0.017 | – |
| | | | | | | Naive Bayes | 0.682±0.031 | 0.653±0.069 | 0.747±0.070 | 0.732±0.035 | all |
| | | | | | | Poly-SVM | 0.788±0.021 | 0.874±0.050 | 0.618±0.102 | 0.847±0.017 | all |
| | | | | | | RBF-SVM | **0.797±0.026** | 0.876±0.030 | 0.642±0.085 | **0.853±0.019** | all |
| | | | | | | Random Forests | **0.806±0.017** | 0.878±0.020 | 0.659±0.049 | **0.859±0.012** | 35037.5 ± 30125.9 |
| | | | | | | $CART_b$ | 0.786±0.017 | 0.899±0.046 | 0.561±0.064 | 0.849±0.014 | 6.6 ± 1.8 |
| | | | | | | $CART_{cv}$ | 0.788±0.020 | 0.897±0.056 | 0.569±0.095 | 0.850±0.017 | 9.4 ± 3.8 |
| | | | | | | $SCM_b$ | 0.791±0.023 | 0.942±0.026 | 0.483±0.044 | **0.858±0.018** | 3.0 ± 0.5 |
| | | | | | | $SCM_{cv}$ | 0.795±0.020 | 0.952±0.020 | 0.475±0.040 | **0.862±0.015** | 4.7 ± 1.7 |
| | cefoxitin | 1789 | 964 | 825 | 62.1 | L1-logistic | 0.857±0.017 | 0.825±0.044 | 0.897±0.050 | 0.864±0.019 | 9038.3 ± 10488.3 |
| | | | | | | L2-logistic | 0.858±0.025 | 0.840±0.030 | 0.880±0.025 | 0.867±0.026 | all* |
| | | | | | | Majority | 0.552±0.016 | **1.000±0.000** | 0.000±0.000 | 0.711±0.014 | – |
| | | | | | | Naive Bayes | 0.767±0.026 | 0.810±0.034 | 0.714±0.042 | 0.793±0.024 | all |
| | | | | | | Poly-SVM | **0.866±0.018** | 0.835±0.032 | 0.903±0.023 | **0.872±0.020** | all |
| | | | | | | RBF-SVM | **0.871±0.018** | 0.833±0.031 | 0.917±0.026 | **0.877±0.019** | all |
| | | | | | | Random Forests | **0.869±0.020** | 0.829±0.040 | 0.918±0.018 | **0.874±0.022** | 47666.8 ± 49259.7 |
| | | | | | | $CART_b$ | 0.855±0.018 | 0.778±0.030 | 0.949±0.030 | 0.855±0.021 | 5.2 ± 1.5 |
| | | | | | | $CART_{cv}$ | **0.869±0.014** | 0.820±0.026 | 0.929±0.025 | **0.873±0.015** | 20.6 ± 5.7 |
| | | | | | | $SCM_b$ | **0.866±0.020** | 0.789±0.029 | **0.961±0.017** | 0.866±0.022 | 6.5 ± 1.2 |
| | | | | | | $SCM_{cv}$ | **0.872±0.017** | 0.819±0.031 | 0.937±0.020 | **0.876±0.020** | 13.5 ± 3.9 |
| | cefuroxime/sodium | 1560 | 1469 | 91 | 55.6 | L1-logistic | **0.984±0.009** | **0.991±0.006** | 0.861±0.075 | **0.992±0.005** | 2833.1 ± 2999.7 |
| | | | | | | L2-logistic | **0.979±0.008** | 0.989±0.007 | 0.804±0.075 | **0.989±0.004** | all* |
| | | | | | | Majority | 0.948±0.011 | **1.000±0.000** | 0.000±0.000 | 0.973±0.006 | – |
| | | | | | | Naive Bayes | 0.777±0.021 | 0.766±0.022 | **0.974±0.035** | 0.867±0.015 | all |
| | | | | | | Poly-SVM | 0.970±0.007 | 0.986±0.005 | 0.653±0.114 | **0.984±0.004** | all |
| | | | | | | RBF-SVM | 0.971±0.006 | 0.987±0.006 | 0.662±0.097 | **0.985±0.004** | all |
| | | | | | | Random Forests | **0.983±0.006** | **0.994±0.006** | 0.780±0.069 | **0.991±0.003** | 1828.9 ± 3685.9 |
| | | | | | | $CART_b$ | **0.986±0.006** | **0.996±0.003** | 0.788±0.080 | **0.992±0.003** | 1.0 ± 0.0 |

Continued on next page

**Table S1.** (Continued)

| Species | Antibiotic | Genomes | Resistant | Susceptible | $k$-mers (millions) | Method | Accuracy | Sensitivity | Specificity | F1 score | Complexity |
|---|---|---|---|---|---|---|---|---|---|---|---|
| | | | | | | CART$_{cv}$ | **0.986 ± 0.006** | **0.996 ± 0.003** | 0.788 ± 0.080 | **0.992 ± 0.003** | 1.0 ± 0.0 |
| | | | | | | SCM$_{b}$ | **0.984 ± 0.006** | **0.995 ± 0.005** | 0.788 ± 0.080 | **0.992 ± 0.003** | 1.2 ± 0.4 |
| | | | | | | SCM$_{cv}$ | **0.984 ± 0.007** | **0.995 ± 0.006** | 0.788 ± 0.080 | **0.992 ± 0.004** | 2.1 ± 2.2 |
| | ceftazidime | 1983 | 1835 | 148 | 65.3 | L1-logistic | 0.956 ± 0.010 | 0.976 ± 0.005 | 0.690 ± 0.073 | **0.976 ± 0.006** | 7469.9 ± 4105.8 |
| | | | | | | L2-logistic | 0.956 ± 0.011 | 0.979 ± 0.007 | 0.673 ± 0.126 | **0.977 ± 0.006** | all* |
| | | | | | | Majority | 0.930 ± 0.014 | **1.000 ± 0.000** | 0.000 ± 0.000 | 0.963 ± 0.007 | – |
| | | | | | | Naive Bayes | 0.776 ± 0.025 | 0.765 ± 0.025 | **0.916 ± 0.060** | 0.863 ± 0.016 | all |
| | | | | | | Poly-SVM | **0.966 ± 0.009** | 0.988 ± 0.005 | 0.675 ± 0.114 | **0.982 ± 0.005** | all |
| | | | | | | RBF-SVM | **0.966 ± 0.009** | 0.987 ± 0.005 | 0.683 ± 0.108 | **0.982 ± 0.005** | all |
| | | | | | | Random Forests | **0.965 ± 0.009** | **0.993 ± 0.003** | 0.595 ± 0.105 | **0.981 ± 0.005** | 23866.7 ± 16647.8 |
| | | | | | | CART$_{b}$ | **0.957 ± 0.008** | 0.988 ± 0.007 | 0.549 ± 0.129 | **0.977 ± 0.004** | 4.2 ± 1.1 |
| | | | | | | CART$_{cv}$ | 0.951 ± 0.006 | 0.980 ± 0.013 | 0.554 ± 0.178 | **0.974 ± 0.004** | 7.4 ± 2.6 |
| | | | | | | SCM$_{b}$ | **0.958 ± 0.008** | 0.989 ± 0.008 | 0.549 ± 0.146 | **0.977 ± 0.004** | 4.8 ± 2.1 |
| | | | | | | SCM$_{cv}$ | **0.959 ± 0.014** | 0.984 ± 0.009 | 0.636 ± 0.138 | **0.978 ± 0.008** | 8.2 ± 3.3 |
| | ceftriaxone | 1842 | 1670 | 172 | 64.7 | L1-logistic | 0.972 ± 0.013 | 0.982 ± 0.009 | 0.878 ± 0.079 | **0.984 ± 0.007** | 3401.8 ± 2110.9 |
| | | | | | | L2-logistic | 0.969 ± 0.011 | 0.982 ± 0.008 | 0.838 ± 0.068 | **0.983 ± 0.006** | all* |
| | | | | | | Majority | 0.910 ± 0.016 | **1.000 ± 0.000** | 0.000 ± 0.000 | 0.953 ± 0.009 | – |
| | | | | | | Naive Bayes | 0.927 ± 0.005 | 0.941 ± 0.012 | 0.789 ± 0.115 | 0.959 ± 0.003 | all |
| | | | | | | Poly-SVM | **0.978 ± 0.007** | 0.985 ± 0.007 | **0.900 ± 0.033** | **0.988 ± 0.004** | all |
| | | | | | | RBF-SVM | **0.976 ± 0.008** | 0.985 ± 0.007 | 0.886 ± 0.057 | **0.987 ± 0.005** | all |
| | | | | | | Random Forests | **0.975 ± 0.009** | 0.986 ± 0.006 | 0.864 ± 0.061 | **0.986 ± 0.005** | 7079.7 ± 9997.1 |
| | | | | | | CART$_{b}$ | **0.980 ± 0.007** | 0.990 ± 0.006 | 0.881 ± 0.041 | **0.989 ± 0.004** | 4.9 ± 0.7 |
| | | | | | | CART$_{cv}$ | **0.978 ± 0.006** | 0.988 ± 0.007 | 0.875 ± 0.042 | **0.988 ± 0.003** | 8.2 ± 2.0 |
| | | | | | | SCM$_{b}$ | **0.982 ± 0.007** | **0.993 ± 0.007** | 0.870 ± 0.049 | **0.990 ± 0.004** | 5.3 ± 0.5 |
| | | | | | | SCM$_{cv}$ | **0.981 ± 0.006** | **0.992 ± 0.008** | 0.872 ± 0.043 | **0.989 ± 0.003** | 6.3 ± 0.8 |
| | ciprofloxacin | 2152 | 1817 | 335 | 71.5 | L1-logistic | 0.952 ± 0.013 | 0.972 ± 0.010 | 0.846 ± 0.038 | 0.972 ± 0.008 | 6018.8 ± 1859.5 |
| | | | | | | L2-logistic | 0.951 ± 0.012 | 0.965 ± 0.011 | 0.873 ± 0.054 | 0.971 ± 0.008 | all* |
| | | | | | | Majority | 0.846 ± 0.021 | **1.000 ± 0.000** | 0.000 ± 0.000 | 0.916 ± 0.012 | – |
| | | | | | | Naive Bayes | 0.884 ± 0.009 | 0.872 ± 0.010 | **0.949 ± 0.025** | 0.927 ± 0.006 | all |
| | | | | | | Poly-SVM | 0.960 ± 0.010 | 0.975 ± 0.008 | 0.879 ± 0.036 | **0.977 ± 0.006** | all |
| | | | | | | RBF-SVM | 0.960 ± 0.011 | 0.976 ± 0.008 | 0.875 ± 0.046 | **0.976 ± 0.006** | all |
| | | | | | | Random Forests | 0.957 ± 0.010 | 0.976 ± 0.009 | 0.854 ± 0.046 | 0.975 ± 0.006 | 8979.5 ± 12331.8 |
| | | | | | | CART$_{b}$ | **0.975 ± 0.009** | 0.988 ± 0.006 | 0.906 ± 0.046 | **0.985 ± 0.005** | 5.0 ± 0.9 |
| | | | | | | CART$_{cv}$ | **0.972 ± 0.010** | 0.983 ± 0.008 | 0.913 ± 0.046 | **0.983 ± 0.006** | 4.9 ± 1.7 |
| | | | | | | SCM$_{b}$ | **0.974 ± 0.008** | 0.984 ± 0.005 | 0.914 ± 0.038 | **0.984 ± 0.005** | 3.2 ± 0.4 |
| | | | | | | SCM$_{cv}$ | **0.974 ± 0.007** | 0.985 ± 0.005 | 0.916 ± 0.037 | **0.985 ± 0.004** | 3.6 ± 0.8 |
| | ertapenem | 361 | 288 | 73 | 31.8 | L1-logistic | **0.965 ± 0.022** | 0.969 ± 0.031 | **0.959 ± 0.054** | **0.978 ± 0.014** | 1748.0 ± 2154.3 |
| | | | | | | L2-logistic | 0.939 ± 0.013 | 0.946 ± 0.018 | 0.923 ± 0.081 | 0.960 ± 0.009 | all* |
| | | | | | | Majority | 0.783 ± 0.032 | **1.000 ± 0.000** | 0.000 ± 0.000 | 0.878 ± 0.020 | – |
| | | | | | | Naive Bayes | 0.831 ± 0.041 | 0.825 ± 0.034 | 0.851 ± 0.106 | 0.884 ± 0.028 | all |
| | | | | | | Poly-SVM | **0.956 ± 0.016** | 0.967 ± 0.019 | 0.919 ± 0.071 | **0.972 ± 0.010** | all |
| | | | | | | RBF-SVM | **0.957 ± 0.015** | 0.967 ± 0.017 | 0.925 ± 0.052 | **0.972 ± 0.010** | all |
| | | | | | | Random Forests | 0.940 ± 0.020 | 0.965 ± 0.022 | 0.862 ± 0.099 | 0.962 ± 0.013 | 6013.6 ± 4539.2 |
| | | | | | | CART$_{b}$ | 0.900 ± 0.019 | 0.937 ± 0.031 | 0.777 ± 0.107 | 0.936 ± 0.013 | 2.7 ± 0.8 |
| | | | | | | CART$_{cv}$ | 0.924 ± 0.028 | 0.958 ± 0.018 | 0.807 ± 0.130 | 0.952 ± 0.017 | 6.2 ± 1.2 |

Continued on next page

**Table S1.** (Continued)

| Species | Antibiotic | Genomes | Resistant | Susceptible | $k$-mers (millions) | Method | Accuracy | Sensitivity | Specificity | F1 score | Complexity |
|---|---|---|---|---|---|---|---|---|---|---|---|
| | | | | | | SCM$_b$ | 0.906 ± 0.022 | 0.958 ± 0.031 | 0.721 ± 0.072 | 0.941 ± 0.014 | 2.8 ± 0.6 |
| | | | | | | SCM$_{cv}$ | 0.904 ± 0.031 | 0.951 ± 0.044 | 0.740 ± 0.084 | 0.939 ± 0.021 | 4.5 ± 1.3 |
| | gentamicin | 2107 | 906 | 1201 | 70.3 | L1-logistic | **0.952 ± 0.010** | **0.926 ± 0.019** | 0.971 ± 0.017 | **0.943 ± 0.011** | 7607.4 ± 7145.7 |
| | | | | | | L2-logistic | **0.948 ± 0.008** | **0.933 ± 0.015** | 0.960 ± 0.011 | **0.939 ± 0.008** | all* |
| | | | | | | Majority | 0.571 ± 0.015 | 0.000 ± 0.000 | **1.000 ± 0.000** | – | – |
| | | | | | | Naive Bayes | 0.760 ± 0.020 | 0.783 ± 0.027 | 0.743 ± 0.028 | 0.737 ± 0.018 | all |
| | | | | | | Poly-SVM | 0.943 ± 0.006 | 0.922 ± 0.011 | 0.959 ± 0.012 | 0.933 ± 0.008 | all |
| | | | | | | RBF-SVM | 0.943 ± 0.005 | **0.925 ± 0.013** | 0.957 ± 0.011 | 0.933 ± 0.007 | all |
| | | | | | | Random Forests | **0.956 ± 0.007** | **0.932 ± 0.020** | 0.974 ± 0.010 | **0.948 ± 0.008** | 42856.8 ± 31470.4 |
| | | | | | | CART$_b$ | **0.949 ± 0.007** | 0.920 ± 0.025 | 0.972 ± 0.011 | **0.940 ± 0.009** | 4.3 ± 1.2 |
| | | | | | | CART$_{cv}$ | **0.948 ± 0.008** | **0.931 ± 0.025** | 0.961 ± 0.014 | **0.939 ± 0.009** | 8.8 ± 3.6 |
| | | | | | | SCM$_b$ | **0.950 ± 0.007** | **0.924 ± 0.022** | 0.970 ± 0.012 | **0.941 ± 0.009** | 3.9 ± 0.7 |
| | | | | | | SCM$_{cv}$ | **0.953 ± 0.009** | **0.931 ± 0.021** | 0.970 ± 0.017 | **0.945 ± 0.011** | 7.9 ± 2.7 |
| | imipenem | 1891 | 660 | 1231 | 62.2 | L1-logistic | **0.949 ± 0.009** | 0.920 ± 0.018 | 0.964 ± 0.012 | 0.927 ± 0.012 | 4562.0 ± 8919.4 |
| | | | | | | L2-logistic | 0.943 ± 0.011 | 0.926 ± 0.021 | 0.953 ± 0.010 | 0.920 ± 0.015 | all* |
| | | | | | | Majority | 0.647 ± 0.023 | 0.000 ± 0.000 | **1.000 ± 0.000** | – | – |
| | | | | | | Naive Bayes | 0.771 ± 0.021 | 0.611 ± 0.034 | 0.858 ± 0.019 | 0.652 ± 0.035 | all |
| | | | | | | Poly-SVM | **0.951 ± 0.008** | 0.925 ± 0.015 | 0.964 ± 0.010 | **0.930 ± 0.012** | all |
| | | | | | | RBF-SVM | **0.951 ± 0.008** | 0.927 ± 0.015 | 0.964 ± 0.010 | **0.930 ± 0.013** | all |
| | | | | | | Random Forests | **0.949 ± 0.008** | 0.923 ± 0.018 | 0.964 ± 0.011 | **0.928 ± 0.013** | 38326.0 ± 22322.1 |
| | | | | | | CART$_b$ | **0.953 ± 0.008** | **0.933 ± 0.019** | 0.964 ± 0.009 | **0.934 ± 0.010** | 2.3 ± 0.5 |
| | | | | | | CART$_{cv}$ | **0.954 ± 0.009** | **0.934 ± 0.020** | 0.966 ± 0.010 | **0.935 ± 0.012** | 3.0 ± 1.1 |
| | | | | | | SCM$_b$ | **0.955 ± 0.009** | **0.937 ± 0.014** | 0.964 ± 0.010 | **0.936 ± 0.011** | 2.0 ± 0.0 |
| | | | | | | SCM$_{cv}$ | **0.956 ± 0.009** | **0.939 ± 0.018** | 0.964 ± 0.010 | **0.937 ± 0.011** | 2.6 ± 0.8 |
| | levofloxacin | 1824 | 1462 | 362 | 58.6 | L1-logistic | **0.964 ± 0.007** | 0.962 ± 0.006 | 0.974 ± 0.029 | **0.978 ± 0.004** | 969.6 ± 1700.7 |
| | | | | | | L2-logistic | 0.955 ± 0.010 | 0.959 ± 0.015 | 0.939 ± 0.038 | **0.972 ± 0.007** | all* |
| | | | | | | Majority | 0.807 ± 0.023 | **1.000 ± 0.000** | 0.000 ± 0.000 | 0.893 ± 0.014 | – |
| | | | | | | Naive Bayes | 0.843 ± 0.029 | 0.808 ± 0.036 | **0.990 ± 0.012** | 0.892 ± 0.023 | all |
| | | | | | | Poly-SVM | **0.960 ± 0.011** | 0.967 ± 0.011 | 0.930 ± 0.035 | **0.975 ± 0.007** | all |
| | | | | | | RBF-SVM | **0.961 ± 0.011** | 0.967 ± 0.013 | 0.931 ± 0.024 | **0.975 ± 0.007** | all |
| | | | | | | Random Forests | **0.960 ± 0.011** | 0.965 ± 0.011 | 0.937 ± 0.042 | **0.975 ± 0.007** | 15165.3 ± 19326.5 |
| | | | | | | CART$_b$ | **0.964 ± 0.006** | 0.977 ± 0.008 | 0.908 ± 0.026 | **0.978 ± 0.004** | 3.1 ± 1.0 |
| | | | | | | CART$_{cv}$ | **0.965 ± 0.007** | 0.977 ± 0.008 | 0.915 ± 0.037 | **0.978 ± 0.004** | 3.1 ± 1.1 |
| | | | | | | SCM$_b$ | **0.963 ± 0.006** | 0.976 ± 0.009 | 0.911 ± 0.031 | **0.977 ± 0.004** | 2.1 ± 0.3 |
| | | | | | | SCM$_{cv}$ | **0.967 ± 0.007** | 0.969 ± 0.009 | 0.957 ± 0.019 | **0.979 ± 0.004** | 3.0 ± 2.9 |
| | meropenem | 2065 | 684 | 1381 | 69.6 | L1-logistic | **0.953 ± 0.010** | **0.924 ± 0.037** | 0.967 ± 0.006 | **0.928 ± 0.019** | 968.5 ± 67.9 |
| | | | | | | L2-logistic | **0.949 ± 0.008** | **0.918 ± 0.030** | 0.964 ± 0.010 | 0.922 ± 0.015 | all* |
| | | | | | | Majority | 0.671 ± 0.017 | 0.000 ± 0.000 | **1.000 ± 0.000** | – | – |
| | | | | | | Naive Bayes | 0.813 ± 0.036 | 0.704 ± 0.091 | 0.866 ± 0.020 | 0.710 ± 0.070 | all |
| | | | | | | Poly-SVM | **0.953 ± 0.011** | 0.908 ± 0.030 | 0.975 ± 0.009 | **0.926 ± 0.020** | all |
| | | | | | | RBF-SVM | **0.951 ± 0.012** | 0.908 ± 0.031 | 0.972 ± 0.009 | 0.923 ± 0.022 | all |
| | | | | | | Random Forests | **0.953 ± 0.010** | 0.912 ± 0.031 | 0.973 ± 0.007 | **0.927 ± 0.018** | 20415.8 ± 24643.6 |
| | | | | | | CART$_b$ | **0.956 ± 0.011** | 0.912 ± 0.030 | 0.978 ± 0.005 | **0.932 ± 0.018** | 3.0 ± 0.0 |
| | | | | | | CART$_{cv}$ | **0.957 ± 0.010** | 0.914 ± 0.031 | 0.977 ± 0.006 | **0.932 ± 0.019** | 2.9 ± 0.6 |
| | | | | | | SCM$_b$ | **0.957 ± 0.008** | **0.924 ± 0.026** | 0.973 ± 0.006 | **0.934 ± 0.015** | 2.0 ± 0.0 |

Continued on next page

**Table S1.** (Continued)

| Species | Antibiotic | Genomes | Resistant | Susceptible | $k$-mers (millions) | Method | Accuracy | Sensitivity | Specificity | F1 score | Complexity |
|---|---|---|---|---|---|---|---|---|---|---|---|
| | | | | | | $SCM_{cv}$ | **0.957±0.009** | **0.925±0.026** | 0.973±0.006 | **0.934±0.016** | 2.1 ± 0.3 |
| | nitrofurantoin | 880 | 790 | 90 | 40.4 | L1-logistic | 0.894±0.010 | 0.940±0.015 | 0.491±0.148 | 0.940±0.006 | 152112.5 ± 285018.2 |
| | | | | | | L2-logistic | 0.907±0.010 | 0.957±0.013 | 0.482±0.100 | 0.949±0.005 | all* |
| | | | | | | Majority | 0.894±0.015 | **1.000±0.000** | 0.000±0.000 | 0.944±0.008 | – |
| | | | | | | Naive Bayes | 0.911±0.013 | 0.969±0.019 | 0.416±0.156 | 0.951±0.008 | all |
| | | | | | | Poly-SVM | **0.926±0.026** | 0.975±0.015 | 0.512±0.136 | **0.959±0.015** | all |
| | | | | | | RBF-SVM | **0.929±0.028** | 0.976±0.023 | **0.525±0.144** | **0.961±0.016** | all |
| | | | | | | Random Forests | **0.923±0.019** | 0.982±0.010 | 0.412±0.141 | **0.958±0.010** | 2203.2 ± 3773.1 |
| | | | | | | $CART_b$ | 0.912±0.025 | 0.965±0.018 | 0.457±0.116 | 0.951±0.015 | 2.8 ± 0.8 |
| | | | | | | $CART_{cv}$ | 0.914±0.022 | 0.974±0.014 | 0.405±0.179 | **0.953±0.012** | 2.8 ± 1.5 |
| | | | | | | $SCM_b$ | 0.911±0.018 | 0.972±0.009 | 0.393±0.128 | 0.951±0.010 | 2.3 ± 0.5 |
| | | | | | | $SCM_{cv}$ | 0.911±0.017 | 0.970±0.018 | 0.404±0.135 | 0.951±0.010 | 5.5 ± 4.3 |
| | ofloxacin | 74 | 47 | 27 | 12.4 | L1-logistic | 0.821±0.069 | 0.856±0.087 | 0.803±0.178 | 0.840±0.081 | 46375.4 ± 143625.1 |
| | | | | | | L2-logistic | 0.793±0.109 | 0.792±0.176 | 0.811±0.199 | 0.801±0.133 | all* |
| | | | | | | Majority | 0.600±0.155 | **1.000±0.000** | 0.000±0.000 | 0.739±0.123 | – |
| | | | | | | Naive Bayes | 0.671±0.136 | 0.641±0.201 | 0.778±0.186 | 0.685±0.149 | all |
| | | | | | | Poly-SVM | 0.807±0.126 | 0.847±0.116 | 0.764±0.208 | 0.827±0.119 | all |
| | | | | | | RBF-SVM | **0.850±0.119** | 0.884±0.104 | **0.825±0.185** | **0.870±0.101** | all |
| | | | | | | Random Forests | 0.829±0.102 | 0.867±0.113 | 0.789±0.207 | 0.853±0.105 | 192.3 ± 192.2 |
| | | | | | | $CART_b$ | 0.814±0.090 | 0.878±0.107 | 0.760±0.202 | 0.836±0.096 | 1.4 ± 0.5 |
| | | | | | | $CART_{cv}$ | 0.786±0.075 | 0.836±0.114 | 0.740±0.171 | 0.814±0.075 | 2.0 ± 0.8 |
| | | | | | | $SCM_b$ | 0.814±0.090 | 0.878±0.107 | 0.760±0.202 | 0.836±0.096 | 1.4 ± 0.5 |
| | | | | | | $SCM_{cv}$ | 0.786±0.101 | 0.848±0.140 | 0.738±0.160 | 0.811±0.107 | 2.3 ± 0.9 |
| | piperacillin/tazobac-tam | 1734 | 1184 | 550 | 63.6 | L1-logistic | 0.862±0.016 | 0.879±0.028 | 0.822±0.061 | 0.897±0.012 | 19247.3 ± 14467.8 |
| | | | | | | L2-logistic | 0.864±0.011 | 0.884±0.024 | 0.819±0.034 | 0.899±0.009 | all* |
| | | | | | | Majority | 0.688±0.022 | **1.000±0.000** | 0.000±0.000 | 0.815±0.016 | – |
| | | | | | | Naive Bayes | 0.766±0.015 | 0.759±0.015 | 0.782±0.025 | 0.817±0.014 | all |
| | | | | | | Poly-SVM | **0.886±0.014** | 0.921±0.015 | 0.807±0.030 | **0.917±0.011** | all |
| | | | | | | RBF-SVM | **0.884±0.014** | 0.921±0.014 | 0.804±0.027 | **0.916±0.011** | all |
| | | | | | | Random Forests | 0.876±0.008 | 0.900±0.019 | 0.822±0.036 | **0.909±0.008** | 47680.8 ± 43218.2 |
| | | | | | | $CART_b$ | 0.839±0.012 | 0.865±0.025 | 0.783±0.046 | 0.881±0.011 | 9.8 ± 2.3 |
| | | | | | | $CART_{cv}$ | 0.842±0.014 | 0.873±0.029 | 0.775±0.080 | 0.884±0.010 | 19.7 ± 7.7 |
| | | | | | | $SCM_b$ | 0.822±0.019 | 0.876±0.022 | 0.702±0.047 | 0.871±0.015 | 4.5 ± 1.0 |
| | | | | | | $SCM_{cv}$ | 0.829±0.009 | 0.817±0.027 | **0.854±0.067** | 0.868±0.010 | 15.0 ± 4.5 |
| | tetracycline | 1553 | 799 | 754 | 56.5 | L1-logistic | **0.877±0.019** | 0.798±0.028 | **0.966±0.018** | **0.872±0.022** | 636.6 ± 66.6 |
| | | | | | | L2-logistic | 0.852±0.039 | 0.817±0.027 | 0.889±0.081 | 0.854±0.032 | all* |
| | | | | | | Majority | 0.526±0.013 | **1.000±0.000** | 0.000±0.000 | 0.689±0.011 | – |
| | | | | | | Naive Bayes | 0.670±0.017 | 0.880±0.016 | 0.435±0.048 | 0.737±0.010 | all |
| | | | | | | Poly-SVM | 0.857±0.019 | 0.818±0.025 | 0.900±0.022 | 0.857±0.019 | all |
| | | | | | | RBF-SVM | 0.855±0.015 | 0.819±0.028 | 0.895±0.020 | 0.856±0.017 | all |
| | | | | | | Random Forests | **0.873±0.022** | 0.796±0.027 | 0.958±0.029 | **0.868±0.022** | 31688.8 ± 26213.0 |
| | | | | | | $CART_b$ | **0.872±0.017** | 0.791±0.026 | **0.961±0.026** | **0.866±0.018** | 3.9 ± 0.3 |
| | | | | | | $CART_{cv}$ | **0.878±0.013** | 0.795±0.029 | **0.969±0.017** | **0.872±0.016** | 7.2 ± 7.8 |
| | | | | | | $SCM_b$ | **0.871±0.018** | 0.788±0.026 | **0.963±0.026** | **0.865±0.020** | 4.0 ± 0.0 |

Continued on next page

**Table S1.** (Continued)

| Species | Antibiotic | Genomes | Resistant | Susceptible | $k$-mers (millions) | Method | Accuracy | Sensitivity | Specificity | F1 score | Complexity |
|---|---|---|---|---|---|---|---|---|---|---|---|
| | | | | | | SCM$_{cv}$ | **0.875±0.016** | 0.796±0.029 | **0.962±0.019** | **0.870±0.019** | 8.0 ± 4.1 |
| | ticarcillin/clavulanic acid | 170 | 75 | 95 | 26.1 | L1-logistic | 0.953±0.021 | **0.952±0.045** | 0.959±0.039 | 0.948±0.029 | 109218.1 ± 254118.1 |
| | | | | | | L2-logistic | **0.965±0.030** | **0.952±0.060** | 0.979±0.027 | **0.961±0.035** | all* |
| | | | | | | Majority | 0.526±0.070 | 0.000±0.000 | **1.000±0.000** | – | – |
| | | | | | | Naive Bayes | 0.935±0.052 | **0.951±0.053** | 0.924±0.083 | 0.932±0.054 | all |
| | | | | | | Poly-SVM | **0.962±0.028** | **0.945±0.050** | 0.979±0.027 | **0.958±0.032** | all |
| | | | | | | RBF-SVM | **0.962±0.028** | **0.945±0.050** | 0.979±0.027 | **0.958±0.032** | all |
| | | | | | | Random Forests | 0.950±0.020 | **0.946±0.041** | 0.957±0.045 | 0.947±0.022 | 198.2 ± 190.5 |
| | | | | | | CART$_b$ | 0.909±0.040 | 0.896±0.086 | 0.930±0.063 | 0.903±0.041 | 1.0 ± 0.0 |
| | | | | | | CART$_{cv}$ | 0.921±0.034 | 0.935±0.061 | 0.912±0.055 | 0.917±0.037 | 2.3 ± 1.3 |
| | | | | | | SCM$_b$ | 0.912±0.039 | 0.907±0.075 | 0.924±0.058 | 0.907±0.040 | 1.3 ± 0.5 |
| | | | | | | SCM$_{cv}$ | 0.918±0.033 | 0.917±0.067 | 0.925±0.049 | 0.912±0.038 | 2.2 ± 0.6 |
| | tobramycin | 1693 | 964 | 729 | 64.4 | L1-logistic | **0.941±0.011** | 0.931±0.016 | **0.955±0.021** | **0.948±0.010** | 9322.4 ± 7525.6 |
| | | | | | | L2-logistic | **0.941±0.013** | 0.926±0.024 | **0.962±0.014** | **0.948±0.013** | all* |
| | | | | | | Majority | 0.583±0.017 | **1.000±0.000** | 0.000±0.000 | 0.737±0.014 | – |
| | | | | | | Naive Bayes | 0.822±0.022 | 0.755±0.037 | 0.916±0.031 | 0.831±0.020 | all |
| | | | | | | Poly-SVM | 0.935±0.013 | 0.939±0.015 | 0.930±0.019 | 0.944±0.011 | all |
| | | | | | | RBF-SVM | 0.934±0.012 | 0.936±0.017 | 0.932±0.018 | 0.943±0.011 | all |
| | | | | | | Random Forests | **0.949±0.013** | 0.944±0.016 | **0.957±0.018** | **0.956±0.011** | 10760.1 ± 16650.0 |
| | | | | | | CART$_b$ | **0.944±0.009** | 0.943±0.013 | 0.946±0.016 | **0.952±0.008** | 5.8 ± 1.8 |
| | | | | | | CART$_{cv}$ | **0.947±0.011** | 0.954±0.016 | 0.938±0.015 | **0.955±0.010** | 11.0 ± 4.3 |
| | | | | | | SCM$_b$ | 0.938±0.009 | 0.941±0.020 | 0.934±0.013 | **0.947±0.009** | 3.8 ± 0.6 |
| | | | | | | SCM$_{cv}$ | **0.940±0.012** | 0.951±0.020 | 0.925±0.020 | **0.949±0.011** | 9.1 ± 2.6 |
| | trimethoprim | 188 | 81 | 107 | 35.1 | L1-logistic | **0.954±0.036** | 0.906±0.090 | 0.990±0.022 | **0.943±0.051** | 156.7 ± 20.9 |
| | | | | | | L2-logistic | 0.927±0.042 | **0.914±0.061** | 0.938±0.043 | 0.915±0.059 | all* |
| | | | | | | Majority | 0.551±0.061 | 0.000±0.000 | **1.000±0.000** | – | – |
| | | | | | | Naive Bayes | 0.611±0.120 | **0.918±0.069** | 0.374±0.192 | 0.681±0.089 | all |
| | | | | | | Poly-SVM | 0.857±0.054 | 0.805±0.089 | 0.902±0.064 | 0.829±0.085 | all |
| | | | | | | RBF-SVM | 0.854±0.051 | 0.793±0.094 | 0.907±0.062 | 0.824±0.082 | all |
| | | | | | | Random Forests | 0.932±0.032 | 0.872±0.084 | 0.981±0.025 | 0.918±0.044 | 1702.8 ± 2378.2 |
| | | | | | | CART$_b$ | 0.932±0.034 | 0.879±0.117 | 0.971±0.034 | 0.914±0.063 | 1.1 ± 0.3 |
| | | | | | | CART$_{cv}$ | 0.932±0.029 | 0.873±0.096 | 0.974±0.036 | 0.915±0.049 | 1.6 ± 1.0 |
| | | | | | | SCM$_b$ | 0.938±0.026 | 0.902±0.074 | 0.967±0.032 | 0.926±0.038 | 1.0 ± 0.0 |
| | | | | | | SCM$_{cv}$ | 0.932±0.026 | 0.869±0.086 | 0.979±0.027 | 0.917±0.040 | 1.8 ± 0.8 |
| | trimethoprim/sul-famethoxazole | 2129 | 1587 | 542 | 71.3 | L1-logistic | **0.935±0.011** | 0.949±0.013 | **0.893±0.018** | **0.956±0.008** | 669.6 ± 86.5 |
| | | | | | | L2-logistic | 0.924±0.013 | 0.942±0.018 | 0.871±0.041 | 0.949±0.010 | all* |
| | | | | | | Majority | 0.752±0.026 | **1.000±0.000** | 0.000±0.000 | 0.858±0.017 | – |
| | | | | | | Naive Bayes | 0.803±0.018 | 0.902±0.019 | 0.504±0.053 | 0.873±0.013 | all |
| | | | | | | Poly-SVM | **0.933±0.013** | 0.963±0.013 | 0.843±0.031 | **0.955±0.010** | all |
| | | | | | | RBF-SVM | **0.932±0.013** | 0.963±0.017 | 0.842±0.034 | **0.955±0.010** | all |
| | | | | | | Random Forests | **0.937±0.016** | 0.968±0.014 | 0.846±0.030 | **0.959±0.011** | 25734.7 ± 20327.6 |
| | | | | | | CART$_b$ | **0.933±0.015** | 0.973±0.012 | 0.813±0.032 | **0.956±0.011** | 3.3 ± 0.5 |
| | | | | | | CART$_{cv}$ | **0.932±0.015** | 0.971±0.016 | 0.814±0.034 | **0.955±0.011** | 5.4 ± 2.9 |

Continued on next page

**Table S1.** (Continued)

| Species | Antibiotic | Genomes | Resistant | Susceptible | $k$-mers (millions) | Method | Accuracy | Sensitivity | Specificity | F1 score | Complexity |
|---|---|---|---|---|---|---|---|---|---|---|---|
| | | | | | | $SCM_b$ | **0.930 ± 0.015** | 0.958 ± 0.017 | 0.845 ± 0.056 | **0.953 ± 0.011** | 4.0 ± 1.6 |
| | | | | | | $SCM_{cv}$ | **0.930 ± 0.012** | 0.952 ± 0.010 | 0.863 ± 0.031 | **0.953 ± 0.009** | 9.3 ± 3.5 |
| *M. tuberculosis* | amikacin | 1145 | 208 | 937 | 7.6 | L1-logistic | **0.951 ± 0.011** | **0.802 ± 0.064** | 0.987 ± 0.010 | 0.862 ± 0.039 | 17781.3 ± 15688.3 |
| | | | | | | L2-logistic | 0.918 ± 0.019 | 0.711 ± 0.064 | 0.970 ± 0.026 | 0.771 ± 0.061 | all* |
| | | | | | | Majority | 0.803 ± 0.024 | 0.000 ± 0.000 | **1.000 ± 0.000** | – | – |
| | | | | | | Naive Bayes | 0.752 ± 0.042 | 0.668 ± 0.098 | 0.773 ± 0.063 | 0.512 ± 0.063 | all |
| | | | | | | Poly-SVM | 0.903 ± 0.021 | 0.613 ± 0.077 | 0.974 ± 0.019 | 0.709 ± 0.074 | all |
| | | | | | | RBF-SVM | 0.902 ± 0.026 | 0.613 ± 0.083 | 0.973 ± 0.020 | 0.708 ± 0.085 | all |
| | | | | | | Random Forests | 0.941 ± 0.010 | 0.748 ± 0.047 | 0.989 ± 0.012 | 0.832 ± 0.034 | 17558.2 ± 19697.7 |
| | | | | | | $CART_b$ | **0.958 ± 0.009** | **0.808 ± 0.056** | **0.994 ± 0.009** | **0.881 ± 0.035** | 1.0 ± 0.0 |
| | | | | | | $CART_{cv}$ | **0.958 ± 0.009** | **0.808 ± 0.056** | **0.994 ± 0.009** | **0.881 ± 0.035** | 1.0 ± 0.0 |
| | | | | | | $SCM_b$ | **0.958 ± 0.009** | **0.808 ± 0.056** | **0.994 ± 0.009** | **0.881 ± 0.035** | 1.0 ± 0.0 |
| | | | | | | $SCM_{cv}$ | **0.958 ± 0.009** | **0.808 ± 0.056** | **0.994 ± 0.009** | **0.881 ± 0.035** | 1.0 ± 0.0 |
| | amoxicillin | 766 | 25 | 741 | 7.3 | L1-logistic | **0.981 ± 0.012** | 0.585 ± 0.322 | **0.993 ± 0.007** | – | 1345.2 ± 331.0 |
| | | | | | | L2-logistic | 0.974 ± 0.012 | 0.261 ± 0.243 | **0.993 ± 0.005** | – | all* |
| | | | | | | Majority | 0.974 ± 0.012 | 0.000 ± 0.000 | **1.000 ± 0.000** | – | – |
| | | | | | | Naive Bayes | 0.962 ± 0.014 | 0.000 ± 0.000 | 0.988 ± 0.007 | – | all |
| | | | | | | Poly-SVM | **0.975 ± 0.014** | 0.212 ± 0.322 | **0.996 ± 0.006** | – | all |
| | | | | | | RBF-SVM | **0.975 ± 0.015** | 0.187 ± 0.328 | **0.996 ± 0.006** | – | all |
| | | | | | | Random Forests | 0.972 ± 0.012 | 0.245 ± 0.249 | **0.992 ± 0.009** | – | 3351.2 ± 4778.6 |
| | | | | | | $CART_b$ | **0.980 ± 0.013** | 0.643 ± 0.370 | **0.991 ± 0.006** | – | 0.9 ± 0.3 |
| | | | | | | $CART_{cv}$ | **0.983 ± 0.008** | 0.693 ± 0.301 | **0.991 ± 0.006** | – | 1.1 ± 0.3 |
| | | | | | | $SCM_b$ | **0.984 ± 0.007** | **0.718 ± 0.294** | **0.991 ± 0.006** | – | 1.0 ± 0.0 |
| | | | | | | $SCM_{cv}$ | **0.984 ± 0.007** | 0.618 ± 0.352 | **0.993 ± 0.006** | – | 1.4 ± 0.7 |
| | capreomycin | 1123 | 204 | 919 | 7.7 | L1-logistic | **0.932 ± 0.020** | 0.772 ± 0.056 | 0.971 ± 0.013 | 0.813 ± 0.044 | 34525.0 ± 17931.7 |
| | | | | | | L2-logistic | 0.902 ± 0.022 | 0.640 ± 0.074 | 0.965 ± 0.019 | 0.712 ± 0.057 | all* |
| | | | | | | Majority | 0.810 ± 0.026 | 0.000 ± 0.000 | **1.000 ± 0.000** | – | – |
| | | | | | | Naive Bayes | 0.783 ± 0.027 | 0.617 ± 0.083 | 0.823 ± 0.023 | 0.517 ± 0.058 | all |
| | | | | | | Poly-SVM | 0.889 ± 0.021 | 0.608 ± 0.063 | 0.955 ± 0.014 | 0.674 ± 0.050 | all |
| | | | | | | RBF-SVM | 0.891 ± 0.022 | 0.601 ± 0.070 | 0.960 ± 0.012 | 0.676 ± 0.059 | all |
| | | | | | | Random Forests | 0.909 ± 0.026 | 0.612 ± 0.077 | 0.980 ± 0.016 | 0.719 ± 0.072 | 15086.7 ± 18855.0 |
| | | | | | | $CART_b$ | **0.938 ± 0.014** | **0.796 ± 0.065** | 0.972 ± 0.011 | **0.829 ± 0.035** | 1.5 ± 0.5 |
| | | | | | | $CART_{cv}$ | **0.938 ± 0.014** | **0.793 ± 0.075** | 0.972 ± 0.014 | **0.828 ± 0.040** | 1.9 ± 1.0 |
| | | | | | | $SCM_b$ | **0.938 ± 0.014** | **0.787 ± 0.062** | 0.975 ± 0.010 | **0.829 ± 0.034** | 1.8 ± 0.4 |
| | | | | | | $SCM_{cv}$ | **0.937 ± 0.014** | **0.791 ± 0.058** | 0.972 ± 0.010 | **0.826 ± 0.031** | 3.5 ± 3.2 |
| | ciprofloxacin | 336 | 35 | 301 | 5.1 | L1-logistic | 0.973 ± 0.020 | 0.902 ± 0.132 | 0.979 ± 0.022 | 0.854 ± 0.095 | 734.6 ± 627.2 |
| | | | | | | L2-logistic | 0.940 ± 0.031 | 0.650 ± 0.194 | 0.968 ± 0.022 | 0.631 ± 0.157 | all* |
| | | | | | | Majority | 0.921 ± 0.022 | 0.000 ± 0.000 | **1.000 ± 0.000** | – | – |
| | | | | | | Naive Bayes | 0.912 ± 0.022 | 0.039 ± 0.087 | 0.987 ± 0.010 | – | all |
| | | | | | | Poly-SVM | 0.951 ± 0.027 | 0.675 ± 0.208 | 0.977 ± 0.020 | 0.684 ± 0.160 | all |
| | | | | | | RBF-SVM | 0.936 ± 0.037 | 0.559 ± 0.289 | 0.972 ± 0.027 | – | all |
| | | | | | | Random Forests | 0.969 ± 0.027 | 0.694 ± 0.248 | **0.995 ± 0.008** | 0.769 ± 0.211 | 932.2 ± 572.8 |
| | | | | | | $CART_b$ | **0.982 ± 0.009** | **0.935 ± 0.106** | 0.985 ± 0.009 | 0.888 ± 0.065 | 1.1 ± 0.3 |
| | | | | | | $CART_{cv}$ | **0.984 ± 0.011** | 0.918 ± 0.107 | 0.989 ± 0.011 | **0.901 ± 0.072** | 1.9 ± 0.9 |
| | | | | | | $SCM_b$ | **0.982 ± 0.009** | 0.918 ± 0.107 | 0.987 ± 0.010 | 0.886 ± 0.064 | 1.3 ± 0.5 |

Continued on next page

**Table S1.** (Continued)

| Species | Antibiotic | Genomes | Resistant | Susceptible | $k$-mers (millions) | Method | Accuracy | Sensitivity | Specificity | F1 score | Complexity |
|---|---|---|---|---|---|---|---|---|---|---|---|
| | | | | | | SCM$_{cv}$ | **0.981±0.010** | 0.885±0.129 | 0.987±0.010 | 0.867±0.095 | 1.5 ± 0.5 |
| | cycloserine | | 72 | 264 | 4.8 | L1-logistic | 0.842±0.028 | **0.618±0.109** | 0.893±0.035 | **0.582±0.094** | 318306.1 ± 239638.1 |
| | | | | | | L2-logistic | 0.839±0.034 | 0.461±0.188 | 0.919±0.043 | 0.491±0.172 | all* |
| | | | | | | Majority | 0.815±0.045 | 0.000±0.000 | **1.000±0.000** | – | – |
| | | | | | | Naive Bayes | 0.813±0.047 | 0.005±0.017 | **0.996±0.009** | – | all |
| | | | | | | Poly-SVM | 0.828±0.030 | 0.527±0.094 | 0.898±0.041 | 0.525±0.092 | all |
| | | | | | | RBF-SVM | **0.858±0.041** | 0.356±0.108 | 0.970±0.023 | 0.475±0.128 | all |
| | | | | | | Random Forests | **0.860±0.027** | 0.417±0.141 | 0.960±0.029 | 0.507±0.115 | 8661.1 ± 11376.0 |
| | | | | | | CART$_b$ | 0.812±0.043 | 0.020±0.063 | **0.993±0.022** | – | 0.3 ± 0.9 |
| | | | | | | CART$_{cv}$ | 0.830±0.043 | 0.418±0.130 | 0.922±0.058 | 0.468±0.150 | 12.7 ± 9.5 |
| | | | | | | SCM$_b$ | 0.800±0.039 | 0.137±0.083 | 0.953±0.043 | 0.190±0.083 | 1.5 ± 0.5 |
| | | | | | | SCM$_{cv}$ | 0.822±0.040 | 0.302±0.152 | 0.942±0.056 | 0.365±0.104 | 6.5 ± 5.4 |
| | ethambutol | 4780 | 748 | 4032 | 11.6 | L1-logistic | **0.924±0.007** | 0.760±0.059 | 0.955±0.015 | 0.761±0.024 | 54872.7 ± 37237.7 |
| | | | | | | L2-logistic | **0.924±0.010** | **0.773±0.070** | 0.952±0.013 | 0.762±0.033 | all* |
| | | | | | | Majority | 0.841±0.006 | 0.000±0.000 | **1.000±0.000** | – | – |
| | | | | | | Naive Bayes | 0.823±0.013 | **0.769±0.053** | 0.833±0.010 | 0.579±0.035 | all |
| | | | | | | Poly-SVM | **0.925±0.011** | 0.722±0.055 | 0.963±0.005 | 0.752±0.041 | all |
| | | | | | | RBF-SVM | 0.922±0.011 | 0.705±0.068 | 0.963±0.005 | 0.740±0.045 | all |
| | | | | | | Random Forests | **0.933±0.011** | 0.752±0.045 | 0.967±0.006 | **0.781±0.037** | 55934.9 ± 51887.2 |
| | | | | | | CART$_b$ | **0.926±0.010** | 0.764±0.053 | 0.956±0.007 | 0.765±0.036 | 13.2 ± 2.1 |
| | | | | | | CART$_{cv}$ | **0.924±0.012** | **0.774±0.067** | 0.952±0.008 | 0.762±0.045 | 20.7 ± 8.0 |
| | | | | | | SCM$_b$ | 0.920±0.006 | 0.743±0.056 | 0.953±0.013 | 0.745±0.020 | 5.7 ± 1.2 |
| | | | | | | SCM$_{cv}$ | 0.923±0.007 | **0.766±0.040** | 0.952±0.011 | 0.758±0.018 | 10.2 ± 2.3 |
| | ethionamide | 564 | 210 | 354 | 5.0 | L1-logistic | 0.781±0.043 | 0.695±0.058 | 0.836±0.046 | **0.709±0.052** | 7671.6 ± 13150.6 |
| | | | | | | L2-logistic | 0.739±0.047 | **0.726±0.079** | 0.746±0.104 | 0.681±0.043 | all* |
| | | | | | | Majority | 0.616±0.032 | 0.000±0.000 | **1.000±0.000** | – | – |
| | | | | | | Naive Bayes | 0.688±0.060 | 0.570±0.115 | 0.762±0.134 | 0.581±0.062 | all |
| | | | | | | Poly-SVM | 0.779±0.028 | 0.623±0.057 | 0.875±0.031 | 0.682±0.042 | all |
| | | | | | | RBF-SVM | 0.786±0.029 | 0.630±0.067 | 0.882±0.033 | 0.691±0.048 | all |
| | | | | | | Random Forests | **0.798±0.036** | 0.592±0.073 | 0.925±0.041 | 0.691±0.054 | 29885.4 ± 18585.0 |
| | | | | | | CART$_b$ | 0.781±0.024 | 0.616±0.062 | 0.884±0.035 | 0.682±0.042 | 2.5 ± 0.5 |
| | | | | | | CART$_{cv}$ | 0.782±0.046 | 0.674±0.078 | 0.849±0.069 | **0.703±0.062** | 25.3 ± 14.0 |
| | | | | | | SCM$_b$ | 0.771±0.026 | 0.605±0.090 | 0.876±0.072 | 0.667±0.043 | 2.2 ± 0.4 |
| | | | | | | SCM$_{cv}$ | 0.762±0.032 | 0.601±0.092 | 0.863±0.091 | 0.658±0.042 | 5.0 ± 1.9 |
| | isoniazid | 5022 | 1719 | 3303 | 11.7 | L1-logistic | **0.962±0.004** | 0.921±0.012 | 0.984±0.005 | **0.944±0.006** | 2242.2 ± 202.1 |
| | | | | | | L2-logistic | 0.941±0.005 | 0.865±0.016 | 0.981±0.006 | 0.910±0.007 | all* |
| | | | | | | Majority | 0.658±0.011 | 0.000±0.000 | **1.000±0.000** | – | – |
| | | | | | | Naive Bayes | 0.789±0.011 | 0.697±0.033 | 0.837±0.010 | 0.693±0.024 | all |
| | | | | | | Poly-SVM | 0.934±0.007 | 0.845±0.019 | 0.980±0.005 | 0.897±0.011 | all |
| | | | | | | RBF-SVM | 0.930±0.007 | 0.849±0.016 | 0.973±0.004 | 0.893±0.010 | all |
| | | | | | | Random Forests | **0.962±0.006** | 0.920±0.016 | 0.984±0.004 | **0.944±0.009** | 78761.3 ± 44953.9 |
| | | | | | | CART$_b$ | **0.962±0.004** | **0.935±0.011** | 0.976±0.008 | **0.944±0.005** | 4.7 ± 1.2 |
| | | | | | | CART$_{cv}$ | **0.963±0.004** | **0.943±0.010** | 0.973±0.007 | **0.945±0.006** | 5.9 ± 2.6 |
| | | | | | | SCM$_b$ | **0.963±0.005** | **0.936±0.016** | 0.977±0.009 | **0.945±0.007** | 4.5 ± 0.5 |
| | | | | | | SCM$_{cv}$ | **0.963±0.004** | **0.941±0.013** | 0.975±0.008 | **0.946±0.006** | 5.0 ± 1.7 |

Continued on next page

**Table S1.** (Continued)

| Species | Antibiotic | Genomes | Resistant | Susceptible | $k$-mers (millions) | Method | Accuracy | Sensitivity | Specificity | F1 score | Complexity |
|---|---|---|---|---|---|---|---|---|---|---|---|
| | kanamycin | 1355 | 297 | 1058 | 7.6 | L1-logistic | 0.947 ± 0.010 | **0.842 ± 0.037** | 0.976 ± 0.007 | 0.874 ± 0.024 | 9464.5 ± 18157.1 |
| | | | | | | L2-logistic | 0.895 ± 0.029 | 0.774 ± 0.060 | 0.929 ± 0.050 | 0.766 ± 0.040 | all* |
| | | | | | | Majority | 0.781 ± 0.012 | 0.000 ± 0.000 | **1.000 ± 0.000** | – | – |
| | | | | | | Naive Bayes | 0.764 ± 0.022 | 0.773 ± 0.027 | 0.762 ± 0.025 | 0.590 ± 0.033 | all |
| | | | | | | Poly-SVM | 0.907 ± 0.010 | 0.714 ± 0.033 | 0.961 ± 0.012 | 0.770 ± 0.023 | all |
| | | | | | | RBF-SVM | 0.911 ± 0.013 | 0.722 ± 0.026 | 0.965 ± 0.014 | 0.782 ± 0.026 | all |
| | | | | | | Random Forests | 0.928 ± 0.010 | 0.754 ± 0.031 | 0.977 ± 0.009 | 0.822 ± 0.025 | 45035.8 ± 13637.5 |
| | | | | | | CART$_b$ | **0.957 ± 0.011** | **0.844 ± 0.040** | 0.989 ± 0.006 | **0.895 ± 0.025** | 3.0 ± 0.0 |
| | | | | | | CART$_{cv}$ | **0.957 ± 0.011** | **0.844 ± 0.040** | 0.989 ± 0.006 | **0.895 ± 0.025** | 3.0 ± 0.0 |
| | | | | | | SCM$_b$ | **0.949 ± 0.012** | **0.844 ± 0.040** | 0.979 ± 0.008 | 0.880 ± 0.028 | 2.0 ± 0.0 |
| | | | | | | SCM$_{cv}$ | **0.949 ± 0.013** | **0.845 ± 0.038** | 0.978 ± 0.010 | 0.879 ± 0.029 | 2.1 ± 0.3 |
| | moxifloxacin | 699 | 57 | 642 | 7.2 | L1-logistic | **0.953 ± 0.017** | 0.722 ± 0.123 | 0.976 ± 0.017 | 0.729 ± 0.097 | 6834.6 ± 7712.5 |
| | | | | | | L2-logistic | 0.933 ± 0.021 | 0.457 ± 0.131 | 0.980 ± 0.013 | 0.543 ± 0.143 | all* |
| | | | | | | Majority | 0.911 ± 0.018 | 0.000 ± 0.000 | **1.000 ± 0.000** | – | – |
| | | | | | | Naive Bayes | 0.883 ± 0.025 | 0.110 ± 0.107 | 0.961 ± 0.024 | – | all |
| | | | | | | Poly-SVM | 0.924 ± 0.012 | 0.231 ± 0.145 | 0.990 ± 0.009 | – | all |
| | | | | | | RBF-SVM | 0.927 ± 0.014 | 0.249 ± 0.142 | **0.991 ± 0.008** | – | all |
| | | | | | | Random Forests | 0.932 ± 0.014 | 0.328 ± 0.117 | **0.991 ± 0.015** | 0.453 ± 0.137 | 9679.0 ± 9024.3 |
| | | | | | | CART$_b$ | **0.957 ± 0.020** | 0.844 ± 0.162 | 0.968 ± 0.014 | 0.769 ± 0.117 | 1.1 ± 0.3 |
| | | | | | | CART$_{cv}$ | **0.960 ± 0.014** | **0.860 ± 0.149** | 0.969 ± 0.015 | **0.782 ± 0.099** | 1.1 ± 0.3 |
| | | | | | | SCM$_b$ | 0.950 ± 0.017 | 0.771 ± 0.185 | 0.969 ± 0.014 | 0.725 ± 0.109 | 1.4 ± 0.5 |
| | | | | | | SCM$_{cv}$ | **0.960 ± 0.014** | **0.854 ± 0.118** | 0.970 ± 0.016 | **0.785 ± 0.075** | 1.4 ± 1.3 |
| | nicotinamide | 167 | 84 | 83 | 4.6 | L1-logistic | 0.803 ± 0.098 | 0.724 ± 0.119 | 0.888 ± 0.123 | 0.791 ± 0.095 | 17764.3 ± 54250.8 |
| | | | | | | L2-logistic | 0.730 ± 0.097 | 0.671 ± 0.079 | 0.782 ± 0.155 | 0.725 ± 0.076 | all* |
| | | | | | | Majority | 0.433 ± 0.038 | 0.400 ± 0.516 | 0.600 ± 0.516 | – | – |
| | | | | | | Naive Bayes | 0.618 ± 0.128 | 0.455 ± 0.271 | 0.826 ± 0.153 | – | all |
| | | | | | | Poly-SVM | 0.752 ± 0.114 | **0.783 ± 0.131** | 0.715 ± 0.140 | 0.767 ± 0.096 | all |
| | | | | | | RBF-SVM | 0.758 ± 0.111 | **0.774 ± 0.137** | 0.742 ± 0.113 | 0.769 ± 0.094 | all |
| | | | | | | Random Forests | 0.736 ± 0.078 | 0.648 ± 0.108 | 0.839 ± 0.083 | 0.718 ± 0.076 | 4379.0 ± 5821.1 |
| | | | | | | CART$_b$ | **0.842 ± 0.057** | 0.746 ± 0.090 | **0.952 ± 0.045** | **0.828 ± 0.066** | 1.0 ± 0.0 |
| | | | | | | CART$_{cv}$ | **0.836 ± 0.063** | 0.746 ± 0.090 | 0.939 ± 0.058 | **0.823 ± 0.069** | 1.3 ± 0.7 |
| | | | | | | SCM$_b$ | **0.842 ± 0.057** | 0.746 ± 0.090 | **0.952 ± 0.045** | **0.828 ± 0.066** | 1.0 ± 0.0 |
| | | | | | | SCM$_{cv}$ | 0.821 ± 0.063 | 0.734 ± 0.099 | 0.919 ± 0.082 | 0.807 ± 0.070 | 2.0 ± 1.2 |
| | ofloxacin | 851 | 307 | 544 | 5.1 | L1-logistic | **0.935 ± 0.017** | **0.888 ± 0.019** | 0.963 ± 0.025 | **0.912 ± 0.018** | 193.9 ± 24.5 |
| | | | | | | L2-logistic | 0.828 ± 0.029 | 0.802 ± 0.055 | 0.844 ± 0.037 | 0.776 ± 0.029 | all* |
| | | | | | | Majority | 0.628 ± 0.031 | 0.000 ± 0.000 | **1.000 ± 0.000** | – | – |
| | | | | | | Naive Bayes | 0.672 ± 0.031 | 0.275 ± 0.155 | 0.907 ± 0.067 | 0.357 ± 0.166 | all |
| | | | | | | Poly-SVM | 0.848 ± 0.026 | 0.791 ± 0.046 | 0.883 ± 0.027 | 0.795 ± 0.033 | all |
| | | | | | | RBF-SVM | 0.844 ± 0.025 | 0.782 ± 0.043 | 0.881 ± 0.026 | 0.788 ± 0.029 | all |
| | | | | | | Random Forests | 0.891 ± 0.029 | 0.808 ± 0.052 | 0.940 ± 0.017 | 0.846 ± 0.041 | 33826.7 ± 23226.3 |
| | | | | | | CART$_b$ | **0.938 ± 0.019** | **0.895 ± 0.020** | 0.964 ± 0.026 | **0.916 ± 0.021** | 1.0 ± 0.0 |
| | | | | | | CART$_{cv}$ | **0.938 ± 0.019** | **0.895 ± 0.020** | 0.964 ± 0.026 | **0.916 ± 0.021** | 1.0 ± 0.0 |
| | | | | | | SCM$_b$ | **0.938 ± 0.019** | **0.895 ± 0.020** | 0.964 ± 0.026 | **0.916 ± 0.021** | 1.0 ± 0.0 |
| | | | | | | SCM$_{cv}$ | **0.937 ± 0.018** | **0.895 ± 0.020** | 0.962 ± 0.026 | **0.914 ± 0.020** | 1.4 ± 0.8 |
| | para-aminosalicylic acid | 378 | 80 | 298 | 4.9 | L1-logistic | **0.883 ± 0.055** | 0.720 ± 0.108 | 0.925 ± 0.051 | **0.712 ± 0.127** | 2944.3 ± 3479.3 |

Continued on next page

**Table S1.** (Continued)

| Species | Antibiotic | Genomes | Resistant | Susceptible | *k*-mers (millions) | Method | Accuracy | Sensitivity | Specificity | F1 score | Complexity |
|---|---|---|---|---|---|---|---|---|---|---|---|
| | | | | | | L2-logistic | 0.843 ± 0.040 | **0.789 ± 0.113** | 0.856 ± 0.065 | 0.666 ± 0.070 | all* |
| | | | | | | Majority | 0.797 ± 0.047 | 0.000 ± 0.000 | **1.000 ± 0.000** | – | – |
| | | | | | | Naive Bayes | 0.856 ± 0.031 | 0.562 ± 0.096 | 0.932 ± 0.031 | 0.607 ± 0.085 | all |
| | | | | | | Poly-SVM | 0.845 ± 0.033 | 0.468 ± 0.109 | 0.942 ± 0.044 | 0.543 ± 0.083 | all |
| | | | | | | RBF-SVM | 0.863 ± 0.033 | 0.562 ± 0.133 | 0.940 ± 0.029 | 0.614 ± 0.101 | all |
| | | | | | | Random Forests | 0.852 ± 0.029 | 0.550 ± 0.126 | 0.932 ± 0.036 | 0.592 ± 0.077 | 4906.8 ± 8827.5 |
| | | | | | | CART$_b$ | 0.835 ± 0.029 | 0.362 ± 0.101 | 0.957 ± 0.033 | 0.459 ± 0.090 | 1.0 ± 0.0 |
| | | | | | | CART$_{cv}$ | 0.823 ± 0.028 | 0.454 ± 0.116 | 0.918 ± 0.032 | 0.499 ± 0.063 | 10.3 ± 9.2 |
| | | | | | | SCM$_b$ | 0.836 ± 0.037 | 0.402 ± 0.127 | 0.948 ± 0.041 | 0.487 ± 0.110 | 1.1 ± 0.3 |
| | | | | | | SCM$_{cv}$ | 0.825 ± 0.035 | 0.418 ± 0.192 | 0.935 ± 0.063 | 0.469 ± 0.115 | 3.0 ± 2.7 |
| | pyrazinamide | 3668 | 377 | 3291 | 10.6 | L1-logistic | **0.944 ± 0.009** | **0.696 ± 0.064** | 0.971 ± 0.007 | **0.707 ± 0.043** | 63589.6 ± 18666.8 |
| | | | | | | L2-logistic | **0.938 ± 0.008** | **0.695 ± 0.069** | 0.965 ± 0.005 | 0.685 ± 0.036 | all* |
| | | | | | | Majority | 0.903 ± 0.009 | 0.000 ± 0.000 | **1.000 ± 0.000** | – | – |
| | | | | | | Naive Bayes | 0.842 ± 0.016 | 0.673 ± 0.070 | 0.860 ± 0.017 | 0.451 ± 0.039 | all |
| | | | | | | Poly-SVM | **0.942 ± 0.008** | 0.665 ± 0.062 | 0.972 ± 0.005 | 0.689 ± 0.036 | all |
| | | | | | | RBF-SVM | **0.941 ± 0.008** | 0.658 ± 0.057 | 0.971 ± 0.005 | 0.682 ± 0.033 | all |
| | | | | | | Random Forests | **0.944 ± 0.009** | 0.633 ± 0.072 | 0.977 ± 0.007 | 0.685 ± 0.047 | 43384.9 ± 32114.6 |
| | | | | | | CART$_b$ | **0.942 ± 0.012** | 0.609 ± 0.067 | 0.978 ± 0.008 | 0.671 ± 0.054 | 11.3 ± 2.0 |
| | | | | | | CART$_{cv}$ | **0.945 ± 0.009** | 0.584 ± 0.060 | 0.984 ± 0.009 | 0.671 ± 0.038 | 17.4 ± 9.4 |
| | | | | | | SCM$_b$ | **0.943 ± 0.008** | 0.571 ± 0.056 | 0.983 ± 0.006 | 0.657 ± 0.038 | 7.6 ± 1.6 |
| | | | | | | SCM$_{cv}$ | **0.941 ± 0.010** | 0.613 ± 0.046 | 0.977 ± 0.009 | 0.669 ± 0.038 | 13.4 ± 4.0 |
| | rifabutin | 161 | 72 | 89 | 4.7 | L1-logistic | **0.828 ± 0.045** | 0.795 ± 0.073 | 0.848 ± 0.094 | 0.814 ± 0.041 | 47.9 ± 11.6 |
| | | | | | | L2-logistic | 0.619 ± 0.062 | 0.621 ± 0.191 | 0.616 ± 0.123 | 0.593 ± 0.125 | all* |
| | | | | | | Majority | 0.522 ± 0.078 | 0.000 ± 0.000 | **1.000 ± 0.000** | – | – |
| | | | | | | Naive Bayes | 0.575 ± 0.082 | 0.584 ± 0.148 | 0.574 ± 0.096 | 0.559 ± 0.111 | all |
| | | | | | | Poly-SVM | 0.641 ± 0.068 | 0.589 ± 0.114 | 0.690 ± 0.064 | 0.605 ± 0.082 | all |
| | | | | | | RBF-SVM | 0.631 ± 0.073 | 0.572 ± 0.103 | 0.684 ± 0.089 | 0.593 ± 0.087 | all |
| | | | | | | Random Forests | 0.678 ± 0.096 | 0.555 ± 0.139 | 0.793 ± 0.132 | 0.616 ± 0.114 | 7461.1 ± 7864.9 |
| | | | | | | CART$_b$ | **0.834 ± 0.047** | **0.819 ± 0.071** | 0.835 ± 0.085 | **0.824 ± 0.047** | 1.0 ± 0.0 |
| | | | | | | CART$_{cv}$ | **0.828 ± 0.054** | **0.813 ± 0.067** | 0.829 ± 0.088 | **0.818 ± 0.052** | 1.7 ± 1.6 |
| | | | | | | SCM$_b$ | **0.834 ± 0.047** | **0.819 ± 0.071** | 0.835 ± 0.085 | **0.824 ± 0.047** | 1.0 ± 0.0 |
| | | | | | | SCM$_{cv}$ | **0.825 ± 0.040** | **0.811 ± 0.054** | 0.822 ± 0.095 | **0.815 ± 0.035** | 1.5 ± 0.7 |
| | rifampin | 5022 | 1396 | 3626 | 11.7 | L1-logistic | **0.974 ± 0.005** | **0.962 ± 0.013** | 0.979 ± 0.005 | **0.954 ± 0.009** | 1376.3 ± 164.7 |
| | | | | | | L2-logistic | 0.958 ± 0.008 | 0.902 ± 0.014 | 0.979 ± 0.007 | 0.922 ± 0.014 | all* |
| | | | | | | Majority | 0.724 ± 0.011 | 0.000 ± 0.000 | **1.000 ± 0.000** | – | – |
| | | | | | | Naive Bayes | 0.828 ± 0.011 | 0.821 ± 0.026 | 0.831 ± 0.011 | 0.725 ± 0.021 | all |
| | | | | | | Poly-SVM | 0.950 ± 0.007 | 0.883 ± 0.014 | 0.976 ± 0.006 | 0.907 ± 0.013 | all |
| | | | | | | RBF-SVM | 0.948 ± 0.009 | 0.885 ± 0.015 | 0.972 ± 0.007 | 0.904 ± 0.015 | all |
| | | | | | | Random Forests | 0.965 ± 0.005 | 0.932 ± 0.011 | 0.978 ± 0.006 | 0.937 ± 0.008 | 77974.3 ± 44091.8 |
| | | | | | | CART$_b$ | **0.977 ± 0.005** | **0.963 ± 0.014** | 0.982 ± 0.005 | **0.958 ± 0.009** | 4.0 ± 0.9 |
| | | | | | | CART$_{cv}$ | **0.978 ± 0.005** | **0.966 ± 0.014** | 0.982 ± 0.006 | **0.960 ± 0.008** | 4.6 ± 1.1 |
| | | | | | | SCM$_b$ | **0.977 ± 0.005** | **0.963 ± 0.014** | 0.982 ± 0.005 | **0.958 ± 0.009** | 3.4 ± 0.5 |
| | | | | | | SCM$_{cv}$ | **0.977 ± 0.005** | **0.966 ± 0.013** | 0.982 ± 0.006 | **0.960 ± 0.008** | 4.2 ± 1.0 |
| | streptomycin | 3406 | 1084 | 2322 | 9.9 | L1-logistic | **0.907 ± 0.004** | **0.865 ± 0.015** | 0.926 ± 0.007 | **0.854 ± 0.007** | 1926.9 ± 130.3 |
| | | | | | | L2-logistic | 0.895 ± 0.008 | 0.817 ± 0.017 | 0.931 ± 0.009 | 0.830 ± 0.015 | all* |

Continued on next page

**Table S1.** (Continued)

| Species | Antibiotic | Genomes | Resistant | Susceptible | $k$-mers (millions) | Method | Accuracy | Sensitivity | Specificity | F1 score | Complexity |
|---|---|---|---|---|---|---|---|---|---|---|---|
| | | | | | | Majority | 0.687±0.009 | 0.000±0.000 | **1.000±0.000** | – | – |
| | | | | | | Naive Bayes | 0.761±0.020 | 0.713±0.030 | 0.783±0.025 | 0.652±0.026 | all |
| | | | | | | Poly-SVM | 0.896±0.008 | 0.797±0.025 | 0.941±0.010 | 0.827±0.016 | all |
| | | | | | | RBF-SVM | 0.892±0.009 | 0.780±0.025 | 0.943±0.012 | 0.818±0.018 | all |
| | | | | | | Random Forests | **0.906±0.008** | 0.805±0.024 | 0.952±0.007 | 0.843±0.016 | 68247.2 ± 46223.9 |
| | | | | | | $CART_b$ | **0.910±0.006** | 0.805±0.027 | 0.958±0.011 | **0.848±0.011** | 10.0 ± 1.4 |
| | | | | | | $CART_{cv}$ | **0.907±0.006** | 0.807±0.032 | 0.953±0.014 | **0.845±0.012** | 17.5 ± 11.8 |
| | | | | | | $SCM_b$ | **0.906±0.010** | 0.783±0.037 | 0.961±0.012 | 0.838±0.019 | 6.8 ± 0.9 |
| | | | | | | $SCM_{cv}$ | **0.908±0.011** | 0.777±0.029 | 0.968±0.009 | 0.841±0.021 | 11.2 ± 3.5 |
| *N. gonorrhoeae* | azithromycin | 392 | 214 | 178 | 4.8 | L1-logistic | **0.942±0.024** | 0.939±0.036 | **0.945±0.039** | **0.945±0.025** | 6095.6 ± 9342.0 |
| | | | | | | L2-logistic | 0.915±0.031 | 0.903±0.048 | 0.928±0.032 | 0.918±0.030 | all* |
| | | | | | | Majority | 0.529±0.035 | **1.000±0.000** | 0.000±0.000 | 0.692±0.030 | – |
| | | | | | | Naive Bayes | 0.736±0.055 | 0.596±0.086 | 0.894±0.045 | 0.702±0.072 | all |
| | | | | | | Poly-SVM | 0.906±0.038 | 0.902±0.057 | 0.910±0.046 | 0.909±0.038 | all |
| | | | | | | RBF-SVM | 0.905±0.035 | 0.902±0.053 | 0.907±0.040 | 0.908±0.035 | all |
| | | | | | | Random Forests | 0.895±0.040 | 0.893±0.049 | 0.897±0.045 | 0.899±0.039 | 4571.7 ± 7185.8 |
| | | | | | | $CART_b$ | **0.936±0.039** | 0.969±0.028 | 0.899±0.057 | **0.942±0.035** | 3.3 ± 0.5 |
| | | | | | | $CART_{cv}$ | 0.929±0.031 | 0.962±0.030 | 0.894±0.047 | **0.935±0.028** | 6.1 ± 3.8 |
| | | | | | | $SCM_b$ | **0.935±0.030** | 0.974±0.023 | 0.891±0.047 | **0.941±0.026** | 3.0 ± 0.0 |
| | | | | | | $SCM_{cv}$ | **0.935±0.033** | 0.972±0.024 | 0.894±0.048 | **0.940±0.029** | 3.5 ± 0.8 |
| | ciprofloxacin | 173 | 106 | 67 | 3.0 | L1-logistic | **0.971±0.024** | 0.974±0.037 | 0.967±0.060 | **0.977±0.018** | 9440.5 ± 24435.6 |
| | | | | | | L2-logistic | **0.968±0.017** | 0.950±0.024 | **1.000±0.000** | **0.974±0.013** | all* |
| | | | | | | Majority | 0.638±0.048 | **1.000±0.000** | 0.000±0.000 | 0.778±0.036 | – |
| | | | | | | Naive Bayes | 0.935±0.053 | 0.899±0.086 | **1.000±0.000** | 0.945±0.049 | all |
| | | | | | | Poly-SVM | **0.971±0.020** | 0.955±0.029 | **1.000±0.000** | **0.977±0.015** | all |
| | | | | | | RBF-SVM | **0.971±0.014** | 0.954±0.021 | **1.000±0.000** | **0.977±0.011** | all |
| | | | | | | Random Forests | **0.968±0.035** | 0.965±0.050 | 0.976±0.039 | **0.975±0.027** | 1026.2 ± 1402.6 |
| | | | | | | $CART_b$ | **0.971±0.031** | **0.991±0.029** | 0.935±0.065 | **0.977±0.024** | 1.0 ± 0.0 |
| | | | | | | $CART_{cv}$ | 0.956±0.040 | 0.977±0.038 | 0.917±0.070 | 0.966±0.030 | 1.1 ± 0.3 |
| | | | | | | $SCM_b$ | **0.971±0.031** | **0.991±0.029** | 0.935±0.065 | **0.977±0.024** | 1.0 ± 0.0 |
| | | | | | | $SCM_{cv}$ | **0.965±0.030** | 0.982±0.032 | 0.935±0.065 | **0.973±0.024** | 1.1 ± 0.3 |
| | erythromycin | 178 | 97 | 81 | | L1-logistic | 0.869±0.041 | 0.887±0.064 | **0.838±0.082** | 0.882±0.041 | 130.7 ± 13.7 |
| | | | | | | L2-logistic | 0.849±0.036 | 0.866±0.071 | 0.818±0.074 | 0.864±0.039 | all* |
| | | | | | | Majority | 0.566±0.061 | **1.000±0.000** | 0.000±0.000 | 0.721±0.051 | – |
| | | | | | | Naive Bayes | 0.843±0.078 | 0.831±0.116 | **0.846±0.076** | 0.850±0.086 | all |
| | | | | | | Poly-SVM | 0.869±0.047 | 0.902±0.064 | 0.818±0.097 | 0.885±0.041 | all |
| | | | | | | RBF-SVM | 0.866±0.049 | 0.902±0.064 | 0.811±0.090 | 0.882±0.044 | all |
| | | | | | | Random Forests | 0.871±0.036 | 0.909±0.038 | 0.818±0.066 | 0.889±0.029 | 413.5 ± 822.4 |
| | | | | | | $CART_b$ | **0.883±0.041** | 0.919±0.047 | 0.831±0.075 | **0.898±0.034** | 1.0 ± 0.0 |
| | | | | | | $CART_{cv}$ | **0.886±0.038** | 0.925±0.047 | 0.831±0.075 | **0.901±0.030** | 1.0 ± 0.0 |
| | | | | | | $SCM_b$ | **0.889±0.044** | 0.925±0.047 | **0.838±0.082** | **0.904±0.035** | 1.0 ± 0.0 |
| | | | | | | $SCM_{cv}$ | 0.874±0.049 | 0.908±0.055 | 0.825±0.079 | 0.889±0.045 | 1.2 ± 0.6 |
| | tetracycline | 142 | 109 | 33 | 2.9 | L1-logistic | 0.929±0.038 | 0.972±0.048 | 0.758±0.180 | 0.954±0.026 | 40683.8 ± 76894.8 |
| | | | | | | L2-logistic | 0.904±0.058 | 0.929±0.065 | **0.801±0.170** | 0.938±0.036 | all* |
| | | | | | | Majority | 0.775±0.073 | **1.000±0.000** | 0.000±0.000 | 0.872±0.047 | – |

Continued on next page

**Table S1.** (Continued)

| Species | Antibiotic | Genomes | Resistant | Susceptible | $k$-mers (millions) | Method | Accuracy | Sensitivity | Specificity | F1 score | Complexity |
|---|---|---|---|---|---|---|---|---|---|---|---|
| | | | | | | Naive Bayes | 0.896±0.064 | 0.920±0.058 | **0.801±0.170** | 0.933±0.039 | all |
| | | | | | | Poly-SVM | **0.950±0.038** | **0.996±0.013** | 0.770±0.177 | **0.969±0.023** | all |
| | | | | | | RBF-SVM | **0.950±0.038** | **0.996±0.013** | 0.770±0.177 | **0.969±0.023** | all |
| | | | | | | Random Forests | 0.936±0.060 | 0.969±0.053 | **0.801±0.170** | 0.959±0.036 | 1010.6 ± 1560.9 |
| | | | | | | CART$_b$ | 0.918±0.053 | 0.966±0.044 | 0.747±0.190 | 0.949±0.033 | 1.0 ± 0.0 |
| | | | | | | CART$_{cv}$ | 0.896±0.043 | 0.940±0.074 | 0.736±0.184 | 0.932±0.034 | 1.5 ± 0.8 |
| | | | | | | SCM$_b$ | 0.896±0.059 | 0.942±0.072 | 0.735±0.175 | 0.933±0.040 | 1.0 ± 0.0 |
| | | | | | | SCM$_{cv}$ | 0.907±0.042 | 0.950±0.062 | 0.747±0.190 | 0.940±0.029 | 1.3 ± 0.5 |
| *P. aeruginosa* | amikacin | 498 | 90 | 408 | 43.2 | L1-logistic | 0.879±0.029 | 0.576±0.095 | 0.942±0.024 | 0.620±0.097 | 33987.3 ± 66238.1 |
| | | | | | | L2-logistic | 0.845±0.030 | 0.553±0.127 | 0.908±0.026 | 0.550±0.092 | all* |
| | | | | | | Majority | 0.824±0.031 | 0.000±0.000 | **1.000±0.000** | – | – |
| | | | | | | Naive Bayes | 0.802±0.030 | **0.630±0.090** | 0.838±0.027 | 0.523±0.088 | all |
| | | | | | | Poly-SVM | 0.848±0.031 | 0.417±0.136 | 0.941±0.024 | 0.479±0.122 | all |
| | | | | | | RBF-SVM | 0.864±0.028 | 0.414±0.101 | 0.960±0.024 | 0.509±0.108 | all |
| | | | | | | Random Forests | 0.874±0.023 | 0.536±0.092 | 0.947±0.009 | 0.594±0.069 | 12334.6 ± 9968.8 |
| | | | | | | CART$_b$ | 0.860±0.041 | 0.422±0.172 | 0.953±0.027 | 0.499±0.150 | 2.7 ± 0.9 |
| | | | | | | CART$_{cv}$ | 0.861±0.037 | 0.482±0.150 | 0.944±0.034 | 0.539±0.116 | 6.8 ± 2.8 |
| | | | | | | SCM$_b$ | **0.891±0.022** | 0.604±0.134 | 0.953±0.021 | 0.650±0.098 | 3.6 ± 0.5 |
| | | | | | | SCM$_{cv}$ | **0.888±0.026** | **0.638±0.098** | 0.940±0.023 | **0.661±0.101** | 4.6 ± 1.3 |
| | ciprofloxacin | 132 | 29 | 103 | 22.5 | L1-logistic | **0.969±0.030** | 0.883±0.150 | **0.994±0.018** | **0.926±0.089** | 381.5 ± 452.3 |
| | | | | | | L2-logistic | 0.808±0.091 | 0.412±0.134 | 0.944±0.048 | 0.519±0.141 | all* |
| | | | | | | Majority | 0.742±0.115 | 0.000±0.000 | **1.000±0.000** | – | – |
| | | | | | | Naive Bayes | 0.708±0.100 | 0.385±0.178 | 0.810±0.092 | – | all |
| | | | | | | Poly-SVM | 0.788±0.125 | 0.287±0.237 | 0.985±0.033 | – | all |
| | | | | | | RBF-SVM | 0.792±0.125 | 0.304±0.246 | 0.985±0.033 | – | all |
| | | | | | | Random Forests | 0.823±0.095 | 0.396±0.166 | 0.978±0.029 | 0.527±0.151 | 1515.5 ± 2132.5 |
| | | | | | | CART$_b$ | **0.965±0.038** | **0.917±0.133** | 0.982±0.029 | **0.934±0.086** | 1.0 ± 0.0 |
| | | | | | | CART$_{cv}$ | **0.962±0.036** | 0.883±0.150 | 0.982±0.029 | 0.914±0.092 | 1.0 ± 0.0 |
| | | | | | | SCM$_b$ | 0.958±0.034 | 0.867±0.145 | 0.982±0.029 | 0.905±0.087 | 1.0 ± 0.0 |
| | | | | | | SCM$_{cv}$ | 0.958±0.034 | 0.867±0.145 | 0.982±0.029 | 0.905±0.087 | 1.0 ± 0.0 |
| | levofloxacin | 491 | 201 | 290 | 43.0 | L1-logistic | **0.937±0.024** | 0.893±0.046 | 0.967±0.033 | 0.921±0.029 | 87.8 ± 9.6 |
| | | | | | | L2-logistic | 0.828±0.043 | 0.789±0.077 | 0.855±0.048 | 0.789±0.060 | all* |
| | | | | | | Majority | 0.588±0.027 | 0.000±0.000 | **1.000±0.000** | – | – |
| | | | | | | Naive Bayes | 0.768±0.051 | 0.666±0.108 | 0.842±0.046 | 0.700±0.078 | all |
| | | | | | | Poly-SVM | 0.773±0.050 | 0.669±0.073 | 0.848±0.058 | 0.708±0.066 | all |
| | | | | | | RBF-SVM | 0.762±0.041 | 0.643±0.103 | 0.846±0.055 | 0.687±0.072 | all |
| | | | | | | Random Forests | 0.874±0.035 | 0.812±0.083 | 0.918±0.040 | 0.840±0.051 | 21600.5 ± 14329.4 |
| | | | | | | CART$_b$ | **0.942±0.028** | 0.926±0.037 | 0.952±0.038 | **0.931±0.031** | 1.1 ± 0.3 |
| | | | | | | CART$_{cv}$ | **0.941±0.021** | **0.963±0.026** | 0.924±0.037 | **0.932±0.020** | 2.5 ± 1.1 |
| | | | | | | SCM$_b$ | **0.939±0.023** | 0.929±0.041 | 0.945±0.034 | **0.927±0.025** | 1.2 ± 0.4 |
| | | | | | | SCM$_{cv}$ | **0.939±0.028** | 0.917±0.048 | 0.954±0.039 | **0.926±0.033** | 1.4 ± 0.5 |
| | meropenem | 380 | 163 | 217 | 39.0 | L1-logistic | **0.720±0.047** | 0.625±0.107 | 0.785±0.043 | 0.646±0.085 | 3827.0 ± 7601.6 |
| | | | | | | L2-logistic | 0.688±0.035 | 0.586±0.079 | 0.761±0.046 | 0.608±0.060 | all* |
| | | | | | | Majority | 0.583±0.035 | 0.000±0.000 | **1.000±0.000** | – | – |
| | | | | | | Naive Bayes | 0.663±0.036 | 0.546±0.057 | 0.746±0.057 | 0.573±0.059 | all |

Continued on next page

**Table S1.** (Continued)

| Species | Antibiotic | Genomes | Resistant | Susceptible | *k*-mers (millions) | Method | Accuracy | Sensitivity | Specificity | F1 score | Complexity |
|---|---|---|---|---|---|---|---|---|---|---|---|
| | | | | | | Poly-SVM | 0.688±0.047 | 0.536±0.105 | 0.798±0.075 | 0.585±0.080 | all |
| | | | | | | RBF-SVM | 0.679±0.038 | 0.535±0.081 | 0.781±0.080 | 0.579±0.058 | all |
| | | | | | | Random Forests | **0.724±0.035** | 0.608±0.059 | 0.805±0.059 | 0.646±0.046 | 8561.0 ± 9899.8 |
| | | | | | | CART$_b$ | **0.724±0.040** | **0.650±0.099** | 0.778±0.069 | **0.659±0.055** | 1.1 ± 0.3 |
| | | | | | | CART$_{cv}$ | 0.711±0.038 | **0.647±0.106** | 0.757±0.072 | 0.647±0.067 | 2.6 ± 3.9 |
| | | | | | | SCM$_b$ | **0.722±0.038** | **0.650±0.099** | 0.776±0.067 | **0.658±0.055** | 1.2 ± 0.4 |
| | | | | | | SCM$_{cv}$ | 0.700±0.038 | 0.619±0.131 | 0.762±0.079 | 0.626±0.073 | 4.1 ± 6.0 |
| *P. difficile* | azithromycin | 461 | 213 | 248 | 19.8 | L1-logistic | 0.947±0.020 | 0.934±0.037 | 0.958±0.026 | 0.941±0.023 | 52144.5 ± 97222.7 |
| | | | | | | L2-logistic | 0.940±0.024 | 0.936±0.034 | 0.944±0.031 | 0.934±0.028 | all* |
| | | | | | | Majority | 0.543±0.027 | 0.000±0.000 | **1.000±0.000** | – | – |
| | | | | | | Naive Bayes | 0.864±0.036 | 0.768±0.051 | 0.946±0.031 | 0.838±0.041 | all |
| | | | | | | Poly-SVM | 0.951±0.023 | 0.943±0.038 | 0.959±0.026 | 0.946±0.028 | all |
| | | | | | | RBF-SVM | 0.947±0.025 | 0.938±0.038 | 0.955±0.031 | 0.941±0.030 | all |
| | | | | | | Random Forests | 0.942±0.016 | 0.929±0.035 | 0.955±0.027 | 0.936±0.020 | 794.1 ± 802.7 |
| | | | | | | CART$_b$ | **0.985±0.009** | **0.981±0.010** | 0.988±0.014 | **0.983±0.011** | 3.0 ± 0.0 |
| | | | | | | CART$_{cv}$ | **0.976±0.017** | 0.965±0.029 | 0.986±0.013 | **0.974±0.019** | 3.9 ± 1.4 |
| | | | | | | SCM$_b$ | **0.978±0.014** | 0.967±0.032 | 0.988±0.014 | **0.976±0.016** | 3.0 ± 0.7 |
| | | | | | | SCM$_{cv}$ | **0.984±0.011** | **0.979±0.013** | 0.988±0.014 | **0.982±0.012** | 3.3 ± 0.5 |
| | ceftriaxone | 212 | 150 | 62 | 11.1 | L1-logistic | 0.902±0.038 | 0.936±0.046 | 0.809±0.161 | **0.934±0.026** | 101937.2 ± 234203.7 |
| | | | | | | L2-logistic | 0.907±0.029 | 0.933±0.041 | 0.844±0.145 | **0.936±0.020** | all* |
| | | | | | | Majority | 0.743±0.055 | **1.000±0.000** | 0.000±0.000 | 0.851±0.036 | – |
| | | | | | | Naive Bayes | 0.824±0.036 | 0.792±0.046 | **0.921±0.062** | 0.869±0.031 | all |
| | | | | | | Poly-SVM | 0.895±0.026 | 0.930±0.045 | 0.798±0.137 | 0.929±0.019 | all |
| | | | | | | RBF-SVM | 0.905±0.034 | 0.942±0.045 | 0.798±0.137 | **0.936±0.024** | all |
| | | | | | | Random Forests | **0.917±0.034** | 0.939±0.042 | 0.858±0.155 | **0.943±0.023** | 436.1 ± 474.3 |
| | | | | | | CART$_b$ | 0.886±0.029 | 0.914±0.057 | 0.822±0.177 | 0.921±0.023 | 1.3 ± 0.5 |
| | | | | | | CART$_{cv}$ | 0.890±0.044 | 0.923±0.060 | 0.810±0.177 | 0.925±0.032 | 2.0 ± 1.2 |
| | | | | | | SCM$_b$ | 0.893±0.036 | 0.929±0.063 | 0.793±0.164 | 0.927±0.028 | 1.2 ± 0.4 |
| | | | | | | SCM$_{cv}$ | 0.890±0.038 | 0.921±0.056 | 0.820±0.175 | 0.925±0.028 | 1.7 ± 0.8 |
| | clarithromycin | 461 | 213 | 248 | 19.8 | L1-logistic | 0.941±0.019 | 0.936±0.044 | 0.946±0.032 | 0.935±0.022 | 153841.5 ± 267155.4 |
| | | | | | | L2-logistic | 0.936±0.018 | 0.924±0.050 | 0.946±0.036 | 0.929±0.020 | all* |
| | | | | | | Majority | 0.543±0.027 | 0.000±0.000 | **1.000±0.000** | – | – |
| | | | | | | Naive Bayes | 0.857±0.028 | 0.748±0.046 | 0.948±0.018 | 0.826±0.033 | all |
| | | | | | | Poly-SVM | 0.947±0.021 | 0.932±0.053 | 0.960±0.028 | 0.941±0.023 | all |
| | | | | | | RBF-SVM | 0.945±0.024 | 0.927±0.057 | 0.960±0.028 | 0.938±0.027 | all |
| | | | | | | Random Forests | 0.937±0.018 | 0.924±0.045 | 0.948±0.026 | 0.930±0.021 | 4726.5 ± 4502.3 |
| | | | | | | CART$_b$ | **0.972±0.022** | 0.970±0.048 | 0.974±0.010 | **0.969±0.025** | 2.9 ± 0.3 |
| | | | | | | CART$_{cv}$ | **0.977±0.008** | **0.981±0.019** | 0.974±0.010 | **0.975±0.009** | 3.0 ± 0.0 |
| | | | | | | SCM$_b$ | **0.972±0.022** | 0.970±0.048 | 0.974±0.010 | **0.969±0.025** | 2.9 ± 0.3 |
| | | | | | | SCM$_{cv}$ | **0.977±0.008** | **0.981±0.019** | 0.974±0.010 | **0.975±0.009** | 3.0 ± 0.0 |
| | clindamycin | 265 | 34 | 231 | 17.8 | L1-logistic | **0.998±0.006** | 0.989±0.035 | **1.000±0.000** | **0.994±0.019** | 1153.1 ± 828.3 |
| | | | | | | L2-logistic | 0.974±0.020 | 0.889±0.107 | 0.986±0.017 | 0.904±0.060 | all* |
| | | | | | | Majority | 0.872±0.057 | 0.000±0.000 | **1.000±0.000** | – | – |
| | | | | | | Naive Bayes | 0.734±0.043 | **1.000±0.000** | 0.695±0.047 | 0.473±0.120 | all |
| | | | | | | Poly-SVM | 0.964±0.017 | 0.877±0.124 | 0.978±0.016 | 0.859±0.041 | all |

Continued on next page

**Table S1.** (Continued)

| Species | Antibiotic | Genomes | Resistant | Susceptible | *k*-mers (millions) | Method | Accuracy | Sensitivity | Specificity | F1 score | Complexity |
|---|---|---|---|---|---|---|---|---|---|---|---|
| | | | | | | RBF-SVM | 0.960 ± 0.023 | 0.889 ± 0.107 | 0.971 ± 0.021 | 0.854 ± 0.043 | all |
| | | | | | | Random Forests | **0.994 ± 0.009** | 0.963 ± 0.059 | **1.000 ± 0.000** | 0.981 ± 0.032 | 60.1 ± 77.0 |
| | | | | | | $CART_b$ | 0.972 ± 0.020 | 0.931 ± 0.112 | 0.981 ± 0.016 | 0.888 ± 0.077 | 1.8 ± 0.4 |
| | | | | | | $CART_{cv}$ | 0.975 ± 0.022 | 0.931 ± 0.112 | 0.985 ± 0.015 | 0.913 ± 0.067 | 1.6 ± 0.5 |
| | | | | | | $SCM_b$ | 0.975 ± 0.022 | 0.967 ± 0.075 | 0.978 ± 0.014 | 0.904 ± 0.083 | 2.0 ± 0.0 |
| | | | | | | $SCM_{cv}$ | 0.975 ± 0.022 | 0.931 ± 0.112 | 0.985 ± 0.015 | 0.913 ± 0.067 | 1.6 ± 0.5 |
| | moxifloxacin | 462 | 188 | 274 | 19.8 | L1-logistic | 0.957 ± 0.027 | 0.921 ± 0.040 | 0.980 ± 0.038 | 0.944 ± 0.033 | 121.8 ± 12.6 |
| | | | | | | L2-logistic | 0.936 ± 0.020 | 0.907 ± 0.042 | 0.955 ± 0.029 | 0.918 ± 0.028 | all* |
| | | | | | | Majority | 0.599 ± 0.029 | 0.000 ± 0.000 | **1.000 ± 0.000** | – | – |
| | | | | | | Naive Bayes | 0.887 ± 0.035 | 0.820 ± 0.065 | 0.931 ± 0.042 | 0.852 ± 0.048 | all |
| | | | | | | Poly-SVM | 0.949 ± 0.014 | 0.904 ± 0.044 | 0.978 ± 0.022 | 0.934 ± 0.020 | all |
| | | | | | | RBF-SVM | 0.951 ± 0.014 | 0.898 ± 0.048 | 0.985 ± 0.018 | 0.935 ± 0.023 | all |
| | | | | | | Random Forests | 0.949 ± 0.015 | 0.904 ± 0.044 | 0.978 ± 0.022 | 0.934 ± 0.021 | 662.2 ± 669.2 |
| | | | | | | $CART_b$ | **0.982 ± 0.009** | **0.959 ± 0.023** | **0.996 ± 0.008** | **0.976 ± 0.012** | 1.0 ± 0.0 |
| | | | | | | $CART_{cv}$ | **0.982 ± 0.009** | **0.959 ± 0.023** | **0.996 ± 0.008** | **0.976 ± 0.012** | 1.1 ± 0.3 |
| | | | | | | $SCM_b$ | **0.982 ± 0.009** | **0.959 ± 0.023** | **0.996 ± 0.008** | **0.976 ± 0.012** | 1.0 ± 0.0 |
| | | | | | | $SCM_{cv}$ | **0.982 ± 0.009** | **0.959 ± 0.023** | **0.996 ± 0.008** | **0.976 ± 0.012** | 1.0 ± 0.0 |
| *S. aureus* | ciprofloxacin | 1229 | 467 | 762 | 12.3 | L1-logistic | **0.983 ± 0.008** | **0.967 ± 0.015** | **0.994 ± 0.005** | **0.978 ± 0.011** | 912.2 ± 1731.0 |
| | | | | | | L2-logistic | **0.975 ± 0.011** | **0.962 ± 0.022** | 0.984 ± 0.011 | 0.969 ± 0.014 | all* |
| | | | | | | Majority | 0.598 ± 0.021 | 0.000 ± 0.000 | **1.000 ± 0.000** | – | – |
| | | | | | | Naive Bayes | 0.892 ± 0.009 | 0.812 ± 0.020 | 0.945 ± 0.013 | 0.858 ± 0.011 | all |
| | | | | | | Poly-SVM | **0.976 ± 0.011** | **0.960 ± 0.019** | 0.986 ± 0.009 | 0.969 ± 0.013 | all |
| | | | | | | RBF-SVM | **0.976 ± 0.010** | **0.960 ± 0.017** | 0.988 ± 0.009 | **0.970 ± 0.012** | all |
| | | | | | | Random Forests | **0.976 ± 0.010** | 0.956 ± 0.023 | 0.989 ± 0.003 | 0.969 ± 0.012 | 16134.3 ± 13601.7 |
| | | | | | | $CART_b$ | **0.983 ± 0.007** | **0.965 ± 0.014** | **0.996 ± 0.004** | **0.979 ± 0.008** | 1.0 ± 0.0 |
| | | | | | | $CART_{cv}$ | **0.983 ± 0.006** | **0.967 ± 0.015** | **0.994 ± 0.006** | **0.978 ± 0.008** | 1.3 ± 0.7 |
| | | | | | | $SCM_b$ | **0.983 ± 0.007** | **0.965 ± 0.014** | **0.996 ± 0.004** | **0.979 ± 0.008** | 1.0 ± 0.0 |
| | | | | | | $SCM_{cv}$ | **0.983 ± 0.006** | **0.965 ± 0.014** | **0.995 ± 0.003** | **0.978 ± 0.008** | 1.2 ± 0.4 |
| | clindamycin | 624 | 350 | 274 | 9.6 | L1-logistic | **0.969 ± 0.013** | 0.978 ± 0.017 | **0.955 ± 0.034** | **0.972 ± 0.012** | 710.4 ± 968.6 |
| | | | | | | L2-logistic | 0.957 ± 0.013 | 0.962 ± 0.029 | **0.949 ± 0.025** | 0.962 ± 0.014 | all* |
| | | | | | | Majority | 0.566 ± 0.045 | **1.000 ± 0.000** | 0.000 ± 0.000 | 0.722 ± 0.039 | – |
| | | | | | | Naive Bayes | 0.866 ± 0.036 | 0.888 ± 0.039 | 0.836 ± 0.052 | 0.882 ± 0.029 | all |
| | | | | | | Poly-SVM | 0.949 ± 0.017 | 0.951 ± 0.038 | 0.944 ± 0.028 | 0.954 ± 0.021 | all |
| | | | | | | RBF-SVM | 0.950 ± 0.011 | 0.954 ± 0.030 | 0.942 ± 0.026 | 0.955 ± 0.015 | all |
| | | | | | | Random Forests | **0.961 ± 0.014** | 0.966 ± 0.026 | **0.953 ± 0.033** | **0.966 ± 0.012** | 3976.7 ± 4930.6 |
| | | | | | | $CART_b$ | **0.961 ± 0.014** | 0.972 ± 0.025 | **0.946 ± 0.033** | **0.965 ± 0.013** | 2.6 ± 1.3 |
| | | | | | | $CART_{cv}$ | 0.958 ± 0.008 | 0.965 ± 0.022 | **0.947 ± 0.030** | **0.963 ± 0.008** | 4.4 ± 2.2 |
| | | | | | | $SCM_b$ | **0.961 ± 0.016** | 0.971 ± 0.020 | **0.947 ± 0.035** | **0.966 ± 0.014** | 2.0 ± 0.0 |
| | | | | | | $SCM_{cv}$ | **0.961 ± 0.016** | 0.971 ± 0.020 | **0.947 ± 0.035** | **0.966 ± 0.014** | 2.2 ± 0.4 |
| | erythromycin | 1305 | 484 | 821 | 12.4 | L1-logistic | **0.976 ± 0.009** | **0.978 ± 0.012** | 0.976 ± 0.016 | **0.970 ± 0.012** | 10563.3 ± 27868.0 |
| | | | | | | L2-logistic | **0.976 ± 0.006** | **0.977 ± 0.008** | 0.976 ± 0.009 | **0.970 ± 0.007** | all* |
| | | | | | | Majority | 0.611 ± 0.019 | 0.000 ± 0.000 | **1.000 ± 0.000** | – | – |
| | | | | | | Naive Bayes | 0.764 ± 0.027 | 0.772 ± 0.060 | 0.759 ± 0.024 | 0.717 ± 0.041 | all |
| | | | | | | Poly-SVM | **0.975 ± 0.010** | **0.979 ± 0.010** | 0.973 ± 0.016 | **0.968 ± 0.013** | all |
| | | | | | | RBF-SVM | **0.973 ± 0.010** | **0.975 ± 0.009** | 0.972 ± 0.018 | **0.966 ± 0.013** | all |

Continued on next page

**Table S1.** (Continued)

| Species | Antibiotic | Genomes | Resistant | Susceptible | $k$-mers (millions) | Method | Accuracy | Sensitivity | Specificity | F1 score | Complexity |
|---|---|---|---|---|---|---|---|---|---|---|---|
| | | | | | | Random Forests | **0.969 ± 0.010** | **0.978 ± 0.007** | 0.963 ± 0.017 | **0.961 ± 0.012** | 6113.3 ± 7868.4 |
| | | | | | | CART$_b$ | **0.976 ± 0.009** | **0.975 ± 0.009** | 0.976 ± 0.016 | **0.969 ± 0.012** | 3.0 ± 0.0 |
| | | | | | | CART$_{cv}$ | **0.974 ± 0.008** | **0.975 ± 0.011** | 0.974 ± 0.016 | **0.967 ± 0.011** | 3.6 ± 1.3 |
| | | | | | | SCM$_b$ | **0.976 ± 0.010** | **0.977 ± 0.008** | 0.976 ± 0.016 | **0.970 ± 0.012** | 3.0 ± 0.0 |
| | | | | | | SCM$_{cv}$ | **0.973 ± 0.012** | **0.975 ± 0.006** | 0.972 ± 0.020 | **0.966 ± 0.015** | 4.6 ± 2.1 |
| | fusidic acid | 986 | 82 | 904 | 11.9 | L1-logistic | **0.984 ± 0.009** | 0.844 ± 0.117 | **0.997 ± 0.003** | 0.896 ± 0.068 | 3120.5 ± 947.0 |
| | | | | | | L2-logistic | 0.969 ± 0.012 | 0.713 ± 0.152 | **0.994 ± 0.005** | 0.793 ± 0.092 | all* |
| | | | | | | Majority | 0.911 ± 0.019 | 0.000 ± 0.000 | **1.000 ± 0.000** | – | – |
| | | | | | | Naive Bayes | 0.675 ± 0.082 | 0.767 ± 0.122 | 0.664 ± 0.092 | 0.301 ± 0.065 | all |
| | | | | | | Poly-SVM | 0.968 ± 0.015 | 0.686 ± 0.153 | **0.995 ± 0.006** | 0.780 ± 0.112 | all |
| | | | | | | RBF-SVM | 0.969 ± 0.015 | 0.712 ± 0.167 | **0.994 ± 0.005** | 0.793 ± 0.108 | all |
| | | | | | | Random Forests | **0.975 ± 0.014** | 0.732 ± 0.145 | **0.999 ± 0.002** | 0.832 ± 0.095 | 4322.9 ± 7941.6 |
| | | | | | | CART$_b$ | **0.976 ± 0.011** | 0.811 ± 0.135 | **0.991 ± 0.005** | 0.843 ± 0.089 | 2.5 ± 0.5 |
| | | | | | | CART$_{cv}$ | **0.984 ± 0.010** | **0.917 ± 0.077** | **0.991 ± 0.005** | **0.907 ± 0.053** | 3.7 ± 0.9 |
| | | | | | | SCM$_b$ | **0.979 ± 0.011** | 0.855 ± 0.114 | **0.991 ± 0.005** | 0.871 ± 0.068 | 2.7 ± 0.5 |
| | | | | | | SCM$_{cv}$ | **0.983 ± 0.010** | **0.917 ± 0.077** | 0.990 ± 0.006 | **0.904 ± 0.054** | 3.2 ± 0.6 |
| | gentamicin | 1306 | 162 | 1144 | 12.4 | L1-logistic | **0.997 ± 0.003** | **0.981 ± 0.018** | **0.999 ± 0.002** | **0.985 ± 0.013** | 136.0 ± 309.3 |
| | | | | | | L2-logistic | **0.993 ± 0.005** | 0.945 ± 0.053 | **0.999 ± 0.003** | 0.966 ± 0.031 | all* |
| | | | | | | Majority | 0.874 ± 0.019 | 0.000 ± 0.000 | **1.000 ± 0.000** | – | – |
| | | | | | | Naive Bayes | 0.949 ± 0.038 | 0.906 ± 0.061 | 0.954 ± 0.042 | 0.826 ± 0.104 | all |
| | | | | | | Poly-SVM | **0.989 ± 0.006** | 0.921 ± 0.056 | **0.998 ± 0.003** | 0.952 ± 0.032 | all |
| | | | | | | RBF-SVM | **0.990 ± 0.006** | 0.921 ± 0.056 | **0.999 ± 0.002** | 0.953 ± 0.033 | all |
| | | | | | | Random Forests | **0.995 ± 0.004** | 0.968 ± 0.043 | **0.999 ± 0.002** | **0.979 ± 0.024** | 432.2 ± 714.2 |
| | | | | | | CART$_b$ | **0.996 ± 0.003** | **0.975 ± 0.019** | **0.999 ± 0.002** | **0.983 ± 0.012** | 1.0 ± 0.0 |
| | | | | | | CART$_{cv}$ | **0.996 ± 0.003** | **0.975 ± 0.019** | **0.999 ± 0.002** | **0.983 ± 0.012** | 1.0 ± 0.0 |
| | | | | | | SCM$_b$ | **0.996 ± 0.003** | **0.975 ± 0.019** | **0.999 ± 0.002** | **0.983 ± 0.012** | 1.0 ± 0.0 |
| | | | | | | SCM$_{cv}$ | **0.994 ± 0.004** | 0.967 ± 0.027 | **0.998 ± 0.002** | **0.977 ± 0.016** | 1.2 ± 0.4 |
| | methicillin | 1593 | 707 | 886 | 13.3 | L1-logistic | **0.988 ± 0.005** | **0.985 ± 0.010** | **0.991 ± 0.007** | **0.987 ± 0.005** | 230.6 ± 212.3 |
| | | | | | | L2-logistic | **0.987 ± 0.003** | **0.984 ± 0.010** | 0.990 ± 0.007 | **0.986 ± 0.003** | all* |
| | | | | | | Majority | 0.544 ± 0.016 | 0.000 ± 0.000 | **1.000 ± 0.000** | – | – |
| | | | | | | Naive Bayes | 0.868 ± 0.019 | 0.875 ± 0.030 | 0.862 ± 0.020 | 0.858 ± 0.019 | all |
| | | | | | | Poly-SVM | **0.987 ± 0.004** | **0.983 ± 0.010** | **0.991 ± 0.007** | **0.986 ± 0.005** | all |
| | | | | | | RBF-SVM | **0.987 ± 0.004** | **0.983 ± 0.010** | 0.990 ± 0.008 | **0.985 ± 0.004** | all |
| | | | | | | Random Forests | **0.987 ± 0.004** | **0.982 ± 0.011** | **0.991 ± 0.007** | **0.986 ± 0.004** | 408.8 ± 570.1 |
| | | | | | | CART$_b$ | **0.987 ± 0.005** | **0.984 ± 0.010** | 0.990 ± 0.007 | **0.986 ± 0.005** | 1.0 ± 0.0 |
| | | | | | | CART$_{cv}$ | **0.987 ± 0.005** | **0.983 ± 0.011** | 0.990 ± 0.007 | **0.985 ± 0.006** | 1.6 ± 1.6 |
| | | | | | | SCM$_b$ | **0.987 ± 0.005** | **0.984 ± 0.010** | 0.990 ± 0.007 | **0.986 ± 0.005** | 1.0 ± 0.0 |
| | | | | | | SCM$_{cv}$ | **0.987 ± 0.005** | **0.983 ± 0.010** | 0.990 ± 0.007 | **0.986 ± 0.005** | 1.9 ± 0.6 |
| | oxacillin | 85 | 39 | 46 | 6.1 | L1-logistic | 0.988 ± 0.025 | 0.980 ± 0.043 | **1.000 ± 0.000** | 0.989 ± 0.023 | 97.6 ± 47.0 |
| | | | | | | L2-logistic | 0.988 ± 0.025 | 0.980 ± 0.043 | **1.000 ± 0.000** | 0.989 ± 0.023 | all* |
| | | | | | | Majority | 0.465 ± 0.131 | 0.100 ± 0.316 | 0.900 ± 0.316 | – | – |
| | | | | | | Naive Bayes | 0.635 ± 0.072 | 0.777 ± 0.136 | 0.500 ± 0.153 | 0.658 ± 0.091 | all |
| | | | | | | Poly-SVM | 0.988 ± 0.025 | 0.980 ± 0.043 | **1.000 ± 0.000** | 0.989 ± 0.023 | all |
| | | | | | | RBF-SVM | **1.000 ± 0.000** | **1.000 ± 0.000** | **1.000 ± 0.000** | **1.000 ± 0.000** | all |
| | | | | | | Random Forests | 0.988 ± 0.025 | 0.980 ± 0.043 | **1.000 ± 0.000** | 0.989 ± 0.023 | 15.9 ± 3.0 |

Continued on next page

**Table S1.** (Continued)

| Species | Antibiotic | Genomes | Resistant | Susceptible | *k*-mers (millions) | Method | Accuracy | Sensitivity | Specificity | F1 score | Complexity |
|---|---|---|---|---|---|---|---|---|---|---|---|
| | | | | | | CART$_b$ | 0.988 ± 0.025 | 0.980 ± 0.043 | **1.000 ± 0.000** | 0.989 ± 0.023 | 1.0 ± 0.0 |
| | | | | | | CART$_{cv}$ | 0.988 ± 0.025 | 0.980 ± 0.043 | **1.000 ± 0.000** | 0.989 ± 0.023 | 1.0 ± 0.0 |
| | | | | | | SCM$_b$ | 0.988 ± 0.025 | 0.980 ± 0.043 | **1.000 ± 0.000** | 0.989 ± 0.023 | 1.0 ± 0.0 |
| | | | | | | SCM$_{cv}$ | 0.988 ± 0.025 | 0.980 ± 0.043 | **1.000 ± 0.000** | 0.989 ± 0.023 | 1.0 ± 0.0 |
| | penicillin | 1042 | 886 | 156 | 12.1 | L1-logistic | **0.974 ± 0.013** | 0.981 ± 0.010 | **0.934 ± 0.051** | **0.985 ± 0.008** | 178881.5 ± 306100.4 |
| | | | | | | L2-logistic | **0.976 ± 0.011** | 0.984 ± 0.009 | **0.931 ± 0.055** | **0.986 ± 0.007** | all* |
| | | | | | | Majority | 0.853 ± 0.022 | **1.000 ± 0.000** | 0.000 ± 0.000 | 0.921 ± 0.013 | – |
| | | | | | | Naive Bayes | 0.518 ± 0.041 | 0.468 ± 0.045 | 0.817 ± 0.059 | 0.622 ± 0.039 | all |
| | | | | | | Poly-SVM | **0.980 ± 0.011** | 0.990 ± 0.007 | 0.923 ± 0.047 | **0.988 ± 0.007** | all |
| | | | | | | RBF-SVM | **0.977 ± 0.013** | 0.988 ± 0.010 | 0.916 ± 0.052 | **0.986 ± 0.008** | all |
| | | | | | | Random Forests | **0.976 ± 0.011** | 0.985 ± 0.007 | **0.927 ± 0.051** | **0.986 ± 0.007** | 4354.3 ± 7268.1 |
| | | | | | | CART$_b$ | **0.973 ± 0.011** | 0.980 ± 0.007 | **0.934 ± 0.051** | **0.984 ± 0.007** | 1.7 ± 0.5 |
| | | | | | | CART$_{cv}$ | **0.971 ± 0.011** | 0.979 ± 0.010 | **0.930 ± 0.047** | **0.983 ± 0.007** | 2.5 ± 0.7 |
| | | | | | | SCM$_b$ | **0.975 ± 0.012** | 0.983 ± 0.007 | **0.927 ± 0.057** | **0.985 ± 0.007** | 1.7 ± 0.5 |
| | | | | | | SCM$_{cv}$ | **0.975 ± 0.012** | 0.985 ± 0.006 | 0.920 ± 0.056 | **0.985 ± 0.007** | 2.5 ± 1.0 |
| | tetracycline | 1232 | 203 | 1029 | 12.3 | L1-logistic | **0.986 ± 0.005** | **0.966 ± 0.029** | **0.991 ± 0.005** | **0.961 ± 0.015** | 78129.4 ± 175973.3 |
| | | | | | | L2-logistic | **0.986 ± 0.006** | **0.957 ± 0.034** | **0.992 ± 0.006** | **0.960 ± 0.017** | all* |
| | | | | | | Majority | 0.820 ± 0.012 | 0.000 ± 0.000 | **1.000 ± 0.000** | – | – |
| | | | | | | Naive Bayes | 0.919 ± 0.012 | 0.774 ± 0.075 | 0.951 ± 0.011 | 0.773 ± 0.044 | all |
| | | | | | | Poly-SVM | **0.982 ± 0.007** | 0.942 ± 0.045 | **0.991 ± 0.005** | 0.949 ± 0.022 | all |
| | | | | | | RBF-SVM | **0.983 ± 0.008** | 0.946 ± 0.044 | **0.991 ± 0.006** | 0.952 ± 0.023 | all |
| | | | | | | Random Forests | **0.987 ± 0.006** | **0.964 ± 0.031** | **0.993 ± 0.007** | **0.965 ± 0.019** | 1572.5 ± 3038.9 |
| | | | | | | CART$_b$ | **0.986 ± 0.005** | **0.966 ± 0.022** | **0.991 ± 0.005** | **0.961 ± 0.015** | 2.0 ± 0.0 |
| | | | | | | CART$_{cv}$ | **0.986 ± 0.005** | **0.966 ± 0.022** | **0.991 ± 0.005** | **0.961 ± 0.015** | 2.0 ± 0.0 |
| | | | | | | SCM$_b$ | **0.986 ± 0.005** | **0.966 ± 0.022** | **0.991 ± 0.005** | **0.961 ± 0.015** | 2.0 ± 0.0 |
| | | | | | | SCM$_{cv}$ | **0.986 ± 0.005** | **0.966 ± 0.022** | **0.991 ± 0.005** | **0.961 ± 0.015** | 2.0 ± 0.0 |
| | trimethoprim/sulfamethoxazole | 320 | 142 | 178 | 6.9 | L1-logistic | 0.947 ± 0.025 | 0.889 ± 0.052 | 0.987 ± 0.018 | 0.931 ± 0.035 | 43517.4 ± 92826.5 |
| | | | | | | L2-logistic | **0.950 ± 0.022** | 0.901 ± 0.049 | 0.985 ± 0.021 | 0.935 ± 0.034 | all* |
| | | | | | | Majority | 0.578 ± 0.054 | 0.000 ± 0.000 | **1.000 ± 0.000** | – | – |
| | | | | | | Naive Bayes | 0.928 ± 0.038 | **0.916 ± 0.038** | 0.936 ± 0.046 | 0.913 ± 0.049 | all |
| | | | | | | Poly-SVM | 0.945 ± 0.025 | 0.889 ± 0.052 | 0.984 ± 0.022 | 0.930 ± 0.035 | all |
| | | | | | | RBF-SVM | 0.941 ± 0.027 | 0.889 ± 0.052 | 0.977 ± 0.029 | 0.923 ± 0.039 | all |
| | | | | | | Random Forests | **0.956 ± 0.019** | 0.901 ± 0.050 | **0.995 ± 0.011** | **0.943 ± 0.026** | 218.3 ± 415.7 |
| | | | | | | CART$_b$ | **0.959 ± 0.020** | 0.901 ± 0.050 | **1.000 ± 0.000** | **0.947 ± 0.027** | 1.0 ± 0.0 |
| | | | | | | CART$_{cv}$ | **0.959 ± 0.020** | 0.901 ± 0.050 | **1.000 ± 0.000** | **0.947 ± 0.027** | 1.0 ± 0.0 |
| | | | | | | SCM$_b$ | **0.959 ± 0.020** | 0.901 ± 0.050 | **1.000 ± 0.000** | **0.947 ± 0.027** | 1.0 ± 0.0 |
| | | | | | | SCM$_{cv}$ | **0.959 ± 0.020** | 0.901 ± 0.050 | **1.000 ± 0.000** | **0.947 ± 0.027** | 1.3 ± 0.9 |
| *S. enterica* | ampicillin | 347 | 279 | 68 | | L1-logistic | 0.875 ± 0.041 | 0.914 ± 0.042 | 0.741 ± 0.172 | 0.920 ± 0.028 | 836.0 ± 972.0 |
| | | | | | | L2-logistic | 0.913 ± 0.026 | 0.940 ± 0.030 | 0.822 ± 0.098 | **0.944 ± 0.018** | all* |
| | | | | | | Majority | 0.791 ± 0.034 | **1.000 ± 0.000** | 0.000 ± 0.000 | 0.883 ± 0.022 | – |
| | | | | | | Naive Bayes | 0.817 ± 0.038 | 0.969 ± 0.017 | 0.246 ± 0.113 | 0.893 ± 0.023 | all |
| | | | | | | Poly-SVM | 0.909 ± 0.033 | 0.971 ± 0.020 | 0.685 ± 0.112 | **0.943 ± 0.021** | all |
| | | | | | | RBF-SVM | 0.907 ± 0.031 | 0.973 ± 0.018 | 0.672 ± 0.111 | **0.943 ± 0.020** | all |
| | | | | | | Random Forests | 0.910 ± 0.037 | 0.967 ± 0.014 | 0.707 ± 0.152 | **0.944 ± 0.023** | 3816.2 ± 4902.6 |

Continued on next page

**Table S1.** (Continued)

| Species | Antibiotic | Genomes | Resistant | Susceptible | $k$-mers (millions) | Method | Accuracy | Sensitivity | Specificity | F1 score | Complexity |
|---|---|---|---|---|---|---|---|---|---|---|---|
| | | | | | | CART$_b$ | 0.894±0.041 | 0.919±0.043 | 0.803±0.165 | 0.932±0.028 | 1.5 ± 0.8 |
| | | | | | | CART$_{cv}$ | **0.925±0.039** | 0.945±0.037 | **0.855±0.087** | **0.951±0.027** | 6.1 ± 3.1 |
| | | | | | | SCM$_b$ | 0.881±0.037 | 0.912±0.037 | 0.769±0.199 | 0.924±0.025 | 1.4 ± 0.5 |
| | | | | | | SCM$_{cv}$ | **0.920±0.040** | 0.950±0.040 | 0.808±0.089 | **0.949±0.027** | 5.5 ± 1.6 |
| | chloramphenicol | | 251 | 96 | | L1-logistic | **0.925±0.039** | 0.953±0.023 | **0.867±0.107** | **0.946±0.030** | 991.2 ± 1463.9 |
| | | | | | | L2-logistic | **0.929±0.033** | 0.959±0.021 | **0.864±0.102** | **0.950±0.026** | all* |
| | | | | | | Majority | 0.709±0.054 | **1.000±0.000** | 0.000±0.000 | 0.828±0.037 | – |
| | | | | | | Naive Bayes | 0.759±0.053 | **0.992±0.011** | 0.198±0.092 | 0.853±0.036 | all |
| | | | | | | Poly-SVM | **0.920±0.030** | 0.970±0.035 | 0.808±0.053 | **0.944±0.023** | all |
| | | | | | | RBF-SVM | **0.928±0.031** | 0.976±0.024 | 0.822±0.084 | **0.949±0.023** | all |
| | | | | | | Random Forests | **0.926±0.029** | 0.984±0.019 | 0.791±0.066 | **0.949±0.022** | 2354.8 ± 2780.1 |
| | | | | | | CART$_b$ | 0.913±0.024 | 0.943±0.035 | 0.848±0.067 | 0.938±0.020 | 1.0 ± 0.0 |
| | | | | | | CART$_{cv}$ | 0.900±0.045 | 0.961±0.032 | 0.761±0.108 | 0.931±0.033 | 3.6 ± 1.3 |
| | | | | | | SCM$_b$ | 0.913±0.024 | 0.943±0.035 | 0.848±0.067 | 0.938±0.020 | 1.0 ± 0.0 |
| | | | | | | SCM$_{cv}$ | 0.907±0.025 | 0.941±0.035 | 0.834±0.089 | 0.934±0.020 | 1.6 ± 1.3 |
| | nalidixic acid | | 35 | 312 | | L1-logistic | **0.978±0.014** | **0.849±0.129** | **0.994±0.008** | **0.876±0.078** | 181.0 ± 42.4 |
| | | | | | | L2-logistic | 0.943±0.029 | 0.622±0.233 | 0.981±0.019 | 0.659±0.175 | all* |
| | | | | | | Majority | 0.906±0.031 | 0.000±0.000 | **1.000±0.000** | – | – |
| | | | | | | Naive Bayes | 0.893±0.034 | 0.049±0.087 | 0.981±0.018 | – | all |
| | | | | | | Poly-SVM | 0.938±0.034 | 0.456±0.269 | **0.991±0.011** | – | all |
| | | | | | | RBF-SVM | 0.942±0.029 | 0.474±0.220 | **0.994±0.008** | 0.592±0.201 | all |
| | | | | | | Random Forests | 0.949±0.024 | 0.589±0.206 | **0.991±0.008** | 0.674±0.136 | 1871.7 ± 3412.6 |
| | | | | | | CART$_b$ | **0.978±0.014** | **0.849±0.129** | **0.994±0.008** | **0.876±0.078** | 1.0 ± 0.0 |
| | | | | | | CART$_{cv}$ | **0.978±0.014** | **0.849±0.129** | **0.994±0.008** | **0.876±0.078** | 1.0 ± 0.0 |
| | | | | | | SCM$_b$ | **0.978±0.014** | **0.849±0.129** | **0.994±0.008** | **0.876±0.078** | 1.0 ± 0.0 |
| | | | | | | SCM$_{cv}$ | **0.978±0.014** | **0.849±0.129** | **0.994±0.008** | **0.876±0.078** | 1.0 ± 0.0 |
| | spectinomycin | 290 | 233 | 57 | 5.6 | L1-logistic | 0.890±0.028 | 0.959±0.028 | **0.629±0.086** | 0.932±0.018 | 4557.6 ± 3948.5 |
| | | | | | | L2-logistic | 0.886±0.023 | 0.959±0.021 | 0.618±0.110 | 0.930±0.015 | all* |
| | | | | | | Majority | 0.791±0.034 | **1.000±0.000** | 0.000±0.000 | 0.883±0.022 | – |
| | | | | | | Naive Bayes | 0.850±0.051 | 0.987±0.018 | 0.341±0.170 | 0.912±0.030 | all |
| | | | | | | Poly-SVM | 0.893±0.028 | 0.983±0.020 | 0.555±0.101 | 0.935±0.018 | all |
| | | | | | | RBF-SVM | 0.888±0.023 | 0.972±0.022 | 0.576±0.073 | 0.932±0.015 | all |
| | | | | | | Random Forests | **0.912±0.033** | **0.993±0.011** | 0.607±0.118 | **0.947±0.021** | 87.8 ± 93.7 |
| | | | | | | CART$_b$ | **0.919±0.026** | **0.996±0.009** | **0.629±0.102** | **0.951±0.016** | 1.0 ± 0.0 |
| | | | | | | CART$_{cv}$ | **0.917±0.023** | **0.994±0.010** | **0.629±0.102** | **0.950±0.014** | 1.4 ± 1.0 |
| | | | | | | SCM$_b$ | **0.919±0.026** | **0.996±0.009** | **0.629±0.102** | **0.951±0.016** | 1.0 ± 0.0 |
| | | | | | | SCM$_{cv}$ | **0.917±0.024** | **0.991±0.011** | **0.638±0.106** | **0.950±0.015** | 1.9 ± 1.6 |
| | streptomycin | 347 | 291 | 56 | 6.9 | L1-logistic | 0.943±0.026 | 0.970±0.029 | 0.826±0.075 | **0.965±0.016** | 90.3 ± 21.2 |
| | | | | | | L2-logistic | 0.929±0.032 | 0.959±0.038 | 0.791±0.062 | 0.956±0.021 | all* |
| | | | | | | Majority | 0.822±0.033 | **1.000±0.000** | 0.000±0.000 | 0.902±0.020 | – |
| | | | | | | Naive Bayes | 0.842±0.032 | 0.975±0.027 | 0.224±0.073 | 0.910±0.020 | all |
| | | | | | | Poly-SVM | 0.925±0.044 | 0.982±0.029 | 0.665±0.156 | 0.955±0.026 | all |
| | | | | | | RBF-SVM | 0.925±0.042 | 0.981±0.030 | 0.668±0.139 | 0.955±0.025 | all |
| | | | | | | Random Forests | 0.938±0.031 | 0.972±0.028 | 0.785±0.135 | 0.962±0.019 | 258.2 ± 370.7 |
| | | | | | | CART$_b$ | 0.943±0.026 | 0.970±0.029 | 0.828±0.069 | **0.965±0.016** | 1.1 ± 0.3 |

Continued on next page

**Table S1.** (Continued)

| Species | Antibiotic | Genomes | Resistant | Susceptible | $k$-mers (millions) | Method | Accuracy | Sensitivity | Specificity | F1 score | Complexity |
|---|---|---|---|---|---|---|---|---|---|---|---|
| | | | | | | CART$_{cv}$ | **0.946±0.030** | 0.975±0.030 | 0.818±0.076 | **0.967±0.019** | 3.3 ± 2.9 |
| | | | | | | SCM$_b$ | **0.946±0.027** | 0.970±0.029 | **0.839±0.067** | **0.967±0.017** | 1.1 ± 0.3 |
| | | | | | | SCM$_{cv}$ | **0.954±0.028** | 0.980±0.030 | **0.831±0.073** | **0.972±0.018** | 1.9 ± 0.6 |
| | sulphonamides | 341 | 306 | 35 | 5.8 | L1-logistic | **0.946±0.017** | 0.987±0.013 | 0.607±0.120 | **0.970±0.010** | 16480.8 ± 24362.6 |
| | | | | | | L2-logistic | **0.951±0.017** | 0.990±0.015 | 0.643±0.136 | **0.973±0.010** | all* |
| | | | | | | Majority | 0.891±0.033 | **1.000±0.000** | 0.000±0.000 | 0.942±0.019 | – |
| | | | | | | Naive Bayes | 0.878±0.033 | 0.964±0.019 | 0.191±0.149 | 0.933±0.019 | all |
| | | | | | | Poly-SVM | 0.943±0.022 | **0.993±0.008** | 0.540±0.138 | **0.968±0.013** | all |
| | | | | | | RBF-SVM | 0.931±0.032 | 0.989±0.013 | 0.480±0.158 | 0.962±0.018 | all |
| | | | | | | Random Forests | **0.954±0.015** | 0.990±0.011 | **0.657±0.165** | **0.975±0.008** | 2614.9 ± 3500.2 |
| | | | | | | CART$_b$ | 0.909±0.021 | 0.973±0.033 | 0.406±0.243 | 0.950±0.011 | 1.6 ± 0.5 |
| | | | | | | CART$_{cv}$ | 0.918±0.026 | 0.981±0.019 | 0.419±0.222 | 0.955±0.014 | 3.0 ± 1.4 |
| | | | | | | SCM$_b$ | 0.913±0.022 | 0.976±0.032 | 0.447±0.263 | 0.952±0.012 | 1.6 ± 0.7 |
| | | | | | | SCM$_{cv}$ | 0.931±0.021 | 0.984±0.016 | 0.492±0.236 | 0.962±0.011 | 2.4 ± 0.7 |
| | tetracycline | 347 | 280 | 67 | 6.9 | L1-logistic | 0.888±0.036 | 0.923±0.042 | 0.769±0.148 | 0.929±0.025 | 1806.7 ± 1360.8 |
| | | | | | | L2-logistic | 0.914±0.032 | 0.949±0.022 | 0.789±0.095 | 0.946±0.021 | all* |
| | | | | | | Majority | 0.793±0.034 | **1.000±0.000** | 0.000±0.000 | 0.884±0.022 | – |
| | | | | | | Naive Bayes | 0.790±0.042 | 0.980±0.026 | 0.064±0.064 | 0.880±0.027 | all |
| | | | | | | Poly-SVM | **0.935±0.016** | 0.971±0.015 | 0.799±0.086 | **0.959±0.010** | all |
| | | | | | | RBF-SVM | **0.933±0.020** | 0.978±0.014 | 0.767±0.068 | **0.959±0.013** | all |
| | | | | | | Random Forests | 0.912±0.041 | 0.958±0.024 | 0.739±0.147 | 0.945±0.025 | 2740.5 ± 2676.5 |
| | | | | | | CART$_b$ | 0.910±0.028 | 0.921±0.019 | **0.877±0.093** | 0.942±0.020 | 2.2 ± 0.6 |
| | | | | | | CART$_{cv}$ | 0.909±0.037 | 0.945±0.032 | 0.778±0.110 | 0.942±0.025 | 6.6 ± 3.2 |
| | | | | | | SCM$_b$ | 0.912±0.036 | 0.923±0.029 | **0.877±0.100** | 0.943±0.025 | 2.0 ± 0.0 |
| | | | | | | SCM$_{cv}$ | 0.906±0.030 | 0.938±0.028 | 0.799±0.143 | 0.940±0.021 | 3.6 ± 1.4 |
| | trimethoprim | 341 | 45 | 296 | 5.8 | L1-logistic | 0.916±0.029 | 0.510±0.232 | 0.969±0.032 | 0.555±0.178 | 109872.9 ± 152324.6 |
| | | | | | | L2-logistic | 0.921±0.027 | **0.544±0.235** | 0.971±0.027 | 0.585±0.159 | all* |
| | | | | | | Majority | 0.887±0.033 | 0.000±0.000 | **1.000±0.000** | – | – |
| | | | | | | Naive Bayes | 0.871±0.040 | 0.220±0.128 | 0.954±0.034 | – | all |
| | | | | | | Poly-SVM | **0.931±0.024** | 0.466±0.155 | 0.990±0.009 | 0.588±0.135 | all |
| | | | | | | RBF-SVM | **0.928±0.026** | 0.458±0.172 | 0.988±0.011 | 0.574±0.145 | all |
| | | | | | | Random Forests | **0.928±0.025** | 0.435±0.189 | 0.990±0.009 | 0.555±0.168 | 1091.7 ± 2035.5 |
| | | | | | | CART$_b$ | **0.937±0.029** | 0.497±0.188 | **0.993±0.011** | **0.626±0.176** | 1.0 ± 0.0 |
| | | | | | | CART$_{cv}$ | **0.929±0.032** | 0.531±0.187 | 0.980±0.025 | **0.617±0.188** | 2.9 ± 2.1 |
| | | | | | | SCM$_b$ | **0.937±0.029** | 0.497±0.188 | **0.993±0.011** | **0.626±0.176** | 1.0 ± 0.0 |
| | | | | | | SCM$_{cv}$ | **0.929±0.032** | 0.531±0.187 | 0.980±0.025 | **0.617±0.188** | 1.7 ± 1.3 |
| *S. haemolyticus* | ciprofloxacin | 120 | 74 | 46 | 5.3 | L1-logistic | **0.925±0.047** | 0.955±0.052 | **0.883±0.102** | **0.938±0.042** | 279.1 ± 616.8 |
| | | | | | | L2-logistic | 0.838±0.057 | 0.894±0.080 | 0.778±0.167 | 0.867±0.060 | all* |
| | | | | | | Majority | 0.629±0.126 | **1.000±0.000** | 0.000±0.000 | 0.765±0.103 | – |
| | | | | | | Naive Bayes | 0.758±0.136 | 0.678±0.216 | **0.892±0.103** | 0.756±0.172 | all |
| | | | | | | Poly-SVM | 0.829±0.077 | 0.856±0.059 | 0.794±0.196 | 0.859±0.067 | all |
| | | | | | | RBF-SVM | 0.846±0.068 | 0.877±0.082 | 0.810±0.117 | 0.871±0.067 | all |
| | | | | | | Random Forests | 0.846±0.040 | 0.903±0.075 | 0.783±0.137 | 0.875±0.042 | 2820.0 ± 3407.8 |
| | | | | | | CART$_b$ | **0.925±0.047** | 0.955±0.052 | **0.883±0.102** | **0.938±0.042** | 1.0 ± 0.0 |
| | | | | | | CART$_{cv}$ | **0.933±0.053** | 0.961±0.054 | **0.892±0.109** | **0.944±0.046** | 1.0 ± 0.0 |

Continued on next page

**Table S1.** (Continued)

| Species | Antibiotic | Genomes | Resistant | Susceptible | $k$-mers (millions) | Method | Accuracy | Sensitivity | Specificity | F1 score | Complexity |
|---|---|---|---|---|---|---|---|---|---|---|---|
| | | | | | | $SCM_b$ | **0.925 ± 0.047** | 0.955 ± 0.052 | **0.883 ± 0.102** | **0.938 ± 0.042** | 1.0 ± 0.0 |
| | | | | | | $SCM_{cv}$ | **0.933 ± 0.053** | 0.961 ± 0.054 | **0.892 ± 0.109** | **0.944 ± 0.046** | 1.0 ± 0.0 |
| | fusidic acid | 114 | 39 | 75 | 5.2 | L1-logistic | **0.832 ± 0.113** | 0.729 ± 0.222 | 0.879 ± 0.107 | **0.749 ± 0.172** | 2732.4 ± 1611.9 |
| | | | | | | L2-logistic | 0.786 ± 0.105 | 0.716 ± 0.181 | 0.825 ± 0.134 | 0.704 ± 0.132 | all* |
| | | | | | | Majority | 0.636 ± 0.091 | 0.000 ± 0.000 | **1.000 ± 0.000** | – | – |
| | | | | | | Naive Bayes | 0.800 ± 0.084 | **0.742 ± 0.117** | 0.821 ± 0.102 | 0.725 ± 0.107 | all |
| | | | | | | Poly-SVM | 0.800 ± 0.123 | 0.702 ± 0.174 | 0.855 ± 0.132 | 0.715 ± 0.155 | all |
| | | | | | | RBF-SVM | 0.809 ± 0.100 | 0.675 ± 0.175 | 0.890 ± 0.084 | 0.714 ± 0.125 | all |
| | | | | | | Random Forests | 0.818 ± 0.132 | 0.723 ± 0.186 | 0.866 ± 0.125 | **0.744 ± 0.182** | 1158.0 ± 2277.1 |
| | | | | | | $CART_b$ | **0.827 ± 0.113** | **0.743 ± 0.241** | 0.872 ± 0.093 | **0.743 ± 0.181** | 1.8 ± 0.4 |
| | | | | | | $CART_{cv}$ | 0.773 ± 0.117 | 0.662 ± 0.243 | 0.820 ± 0.120 | 0.664 ± 0.199 | 2.0 ± 0.7 |
| | | | | | | $SCM_b$ | **0.827 ± 0.109** | 0.700 ± 0.256 | 0.893 ± 0.107 | 0.728 ± 0.182 | 1.7 ± 0.5 |
| | | | | | | $SCM_{cv}$ | 0.782 ± 0.137 | 0.654 ± 0.273 | 0.845 ± 0.096 | 0.664 ± 0.233 | 2.5 ± 1.4 |
| | tetracycline | 100 | 37 | 63 | 5.1 | L1-logistic | 0.780 ± 0.067 | 0.669 ± 0.107 | 0.853 ± 0.082 | 0.698 ± 0.064 | 1550.3 ± 1271.7 |
| | | | | | | L2-logistic | **0.810 ± 0.061** | 0.744 ± 0.102 | 0.856 ± 0.074 | **0.745 ± 0.073** | all* |
| | | | | | | Majority | 0.620 ± 0.082 | 0.000 ± 0.000 | **1.000 ± 0.000** | – | – |
| | | | | | | Naive Bayes | 0.780 ± 0.079 | **0.809 ± 0.155** | 0.769 ± 0.136 | 0.731 ± 0.101 | all |
| | | | | | | Poly-SVM | 0.745 ± 0.093 | 0.574 ± 0.259 | 0.874 ± 0.091 | 0.600 ± 0.187 | all |
| | | | | | | RBF-SVM | 0.750 ± 0.085 | 0.584 ± 0.244 | 0.874 ± 0.091 | 0.613 ± 0.171 | all |
| | | | | | | Random Forests | 0.795 ± 0.055 | 0.623 ± 0.191 | 0.917 ± 0.075 | 0.685 ± 0.108 | 1071.9 ± 1233.9 |
| | | | | | | $CART_b$ | 0.785 ± 0.082 | 0.635 ± 0.163 | 0.892 ± 0.106 | 0.688 ± 0.112 | 1.0 ± 0.0 |
| | | | | | | $CART_{cv}$ | 0.735 ± 0.088 | 0.686 ± 0.174 | 0.772 ± 0.156 | 0.658 ± 0.087 | 2.7 ± 1.9 |
| | | | | | | $SCM_b$ | 0.770 ± 0.116 | 0.603 ± 0.155 | 0.885 ± 0.159 | 0.667 ± 0.139 | 1.0 ± 0.0 |
| | | | | | | $SCM_{cv}$ | 0.730 ± 0.075 | 0.561 ± 0.128 | 0.838 ± 0.075 | 0.606 ± 0.084 | 2.2 ± 0.9 |
| *S. pneumoniae* | cefuroxime | 113 | 68 | 45 | 5.7 | L1-logistic | **0.977 ± 0.039** | 0.983 ± 0.038 | 0.966 ± 0.060 | **0.979 ± 0.038** | 777.9 ± 1049.9 |
| | | | | | | L2-logistic | 0.932 ± 0.069 | 0.934 ± 0.060 | 0.947 ± 0.088 | 0.938 ± 0.072 | all* |
| | | | | | | Majority | 0.618 ± 0.127 | **1.000 ± 0.000** | 0.000 ± 0.000 | 0.757 ± 0.104 | – |
| | | | | | | Naive Bayes | 0.877 ± 0.080 | 0.807 ± 0.164 | 0.978 ± 0.049 | 0.875 ± 0.112 | all |
| | | | | | | Poly-SVM | 0.900 ± 0.082 | 0.894 ± 0.102 | 0.931 ± 0.092 | 0.911 ± 0.079 | all |
| | | | | | | RBF-SVM | 0.891 ± 0.084 | 0.879 ± 0.113 | 0.937 ± 0.087 | 0.901 ± 0.083 | all |
| | | | | | | Random Forests | **0.986 ± 0.031** | 0.983 ± 0.038 | **0.992 ± 0.024** | **0.986 ± 0.035** | 290.2 ± 691.2 |
| | | | | | | $CART_b$ | 0.945 ± 0.052 | 0.976 ± 0.050 | 0.903 ± 0.079 | 0.951 ± 0.050 | 1.0 ± 0.0 |
| | | | | | | $CART_{cv}$ | 0.941 ± 0.043 | 0.969 ± 0.052 | 0.897 ± 0.084 | 0.947 ± 0.046 | 1.0 ± 0.0 |
| | | | | | | $SCM_b$ | 0.945 ± 0.052 | 0.976 ± 0.050 | 0.903 ± 0.079 | 0.951 ± 0.050 | 1.0 ± 0.0 |
| | | | | | | $SCM_{cv}$ | 0.936 ± 0.038 | 0.956 ± 0.051 | 0.911 ± 0.089 | 0.944 ± 0.043 | 1.2 ± 0.4 |
| | chloramphenicol | 409 | 149 | 260 | 6.4 | L1-logistic | 0.948 ± 0.022 | **0.950 ± 0.023** | 0.947 ± 0.036 | 0.927 ± 0.031 | 1391.5 ± 1844.4 |
| | | | | | | L2-logistic | 0.949 ± 0.020 | 0.936 ± 0.029 | 0.957 ± 0.028 | 0.928 ± 0.027 | all* |
| | | | | | | Majority | 0.654 ± 0.036 | 0.000 ± 0.000 | **1.000 ± 0.000** | – | – |
| | | | | | | Naive Bayes | 0.910 ± 0.013 | 0.936 ± 0.034 | 0.896 ± 0.017 | 0.877 ± 0.020 | all |
| | | | | | | Poly-SVM | 0.946 ± 0.021 | 0.929 ± 0.039 | 0.955 ± 0.027 | 0.922 ± 0.030 | all |
| | | | | | | RBF-SVM | 0.944 ± 0.023 | 0.925 ± 0.041 | 0.955 ± 0.031 | 0.920 ± 0.032 | all |
| | | | | | | Random Forests | **0.957 ± 0.019** | **0.947 ± 0.030** | 0.962 ± 0.026 | **0.938 ± 0.027** | 92.2 ± 110.8 |
| | | | | | | $CART_b$ | **0.960 ± 0.018** | **0.951 ± 0.035** | 0.966 ± 0.026 | **0.943 ± 0.026** | 1.0 ± 0.0 |
| | | | | | | $CART_{cv}$ | **0.959 ± 0.018** | **0.947 ± 0.035** | 0.966 ± 0.026 | **0.941 ± 0.026** | 1.0 ± 0.0 |
| | | | | | | $SCM_b$ | **0.960 ± 0.018** | **0.951 ± 0.035** | 0.966 ± 0.026 | **0.943 ± 0.026** | 1.0 ± 0.0 |

Continued on next page

**Table S1.** (Continued)

| Species | Antibiotic | Genomes | Resistant | Susceptible | $k$-mers (millions) | Method | Accuracy | Sensitivity | Specificity | F1 score | Complexity |
|---|---|---|---|---|---|---|---|---|---|---|---|
| | | | | | | SCM$_{cv}$ | **0.959±0.018** | **0.947±0.035** | 0.966±0.026 | **0.941±0.026** | 1.0 ± 0.0 |
| | clindamycin | 145 | 28 | 117 | 6.0 | L1-logistic | **0.986±0.024** | **0.950±0.127** | **0.996±0.013** | 0.959±0.082 | 211.2 ± 208.2 |
| | | | | | | L2-logistic | 0.948±0.034 | 0.833±0.187 | 0.979±0.022 | 0.842±0.112 | all* |
| | | | | | | Majority | 0.810±0.080 | 0.000±0.000 | **1.000±0.000** | – | – |
| | | | | | | Naive Bayes | 0.907±0.046 | 0.809±0.233 | 0.932±0.045 | 0.735±0.192 | all |
| | | | | | | Poly-SVM | 0.938±0.048 | 0.760±0.292 | 0.979±0.022 | – | all |
| | | | | | | RBF-SVM | 0.945±0.037 | 0.780±0.236 | 0.984±0.021 | 0.802±0.190 | all |
| | | | | | | Random Forests | **0.986±0.024** | **0.950±0.127** | **0.996±0.014** | **0.962±0.079** | 225.7 ± 532.5 |
| | | | | | | CART$_{b}$ | **0.990±0.023** | **0.950±0.127** | **1.000±0.000** | **0.970±0.079** | 1.0 ± 0.0 |
| | | | | | | CART$_{cv}$ | **0.990±0.023** | **0.950±0.127** | **1.000±0.000** | **0.970±0.079** | 1.0 ± 0.0 |
| | | | | | | SCM$_{b}$ | **0.990±0.023** | **0.950±0.127** | **1.000±0.000** | **0.970±0.079** | 1.0 ± 0.0 |
| | | | | | | SCM$_{cv}$ | **0.990±0.023** | **0.950±0.127** | **1.000±0.000** | **0.970±0.079** | 1.0 ± 0.0 |
| | erythromycin | 324 | 247 | 77 | 6.3 | L1-logistic | **0.961±0.028** | 0.970±0.023 | 0.932±0.086 | **0.974±0.019** | 4386.0 ± 4378.0 |
| | | | | | | L2-logistic | 0.948±0.029 | 0.966±0.033 | 0.897±0.075 | **0.965±0.020** | all* |
| | | | | | | Majority | 0.742±0.047 | **1.000±0.000** | 0.000±0.000 | 0.851±0.031 | – |
| | | | | | | Naive Bayes | 0.706±0.034 | 0.716±0.034 | 0.686±0.133 | 0.783±0.029 | all |
| | | | | | | Poly-SVM | 0.941±0.030 | 0.964±0.034 | 0.872±0.077 | 0.960±0.022 | all |
| | | | | | | RBF-SVM | 0.941±0.032 | 0.962±0.035 | 0.879±0.069 | 0.960±0.023 | all |
| | | | | | | Random Forests | 0.934±0.040 | 0.976±0.022 | 0.823±0.111 | 0.956±0.028 | 4155.2 ± 5134.4 |
| | | | | | | CART$_{b}$ | **0.952±0.026** | 0.951±0.027 | **0.950±0.078** | **0.966±0.019** | 2.2 ± 0.4 |
| | | | | | | CART$_{cv}$ | 0.948±0.027 | 0.951±0.031 | 0.937±0.080 | 0.964±0.019 | 3.1 ± 1.1 |
| | | | | | | SCM$_{b}$ | **0.952±0.026** | 0.951±0.027 | **0.950±0.078** | **0.966±0.019** | 2.2 ± 0.4 |
| | | | | | | SCM$_{cv}$ | 0.950±0.030 | 0.959±0.024 | 0.920±0.096 | **0.966±0.021** | 2.8 ± 0.8 |
| | meropenem | 114 | 32 | 82 | 5.8 | L1-logistic | **0.864±0.074** | **0.907±0.172** | 0.851±0.122 | **0.783±0.140** | 411.8 ± 470.4 |
| | | | | | | L2-logistic | **0.868±0.079** | 0.812±0.164 | 0.888±0.075 | 0.762±0.201 | all* |
| | | | | | | Majority | 0.705±0.086 | 0.000±0.000 | **1.000±0.000** | – | – |
| | | | | | | Naive Bayes | 0.836±0.089 | 0.818±0.192 | 0.839±0.071 | 0.721±0.214 | all |
| | | | | | | Poly-SVM | 0.818±0.068 | 0.676±0.184 | 0.883±0.077 | 0.666±0.178 | all |
| | | | | | | RBF-SVM | 0.818±0.068 | 0.676±0.184 | 0.883±0.077 | 0.666±0.178 | all |
| | | | | | | Random Forests | 0.850±0.074 | 0.799±0.166 | 0.893±0.100 | 0.742±0.154 | 313.2 ± 209.0 |
| | | | | | | CART$_{b}$ | 0.850±0.077 | **0.914±0.101** | 0.827±0.111 | 0.769±0.150 | 1.0 ± 0.0 |
| | | | | | | CART$_{cv}$ | 0.827±0.093 | 0.733±0.240 | 0.850±0.117 | 0.685±0.220 | 2.4 ± 1.4 |
| | | | | | | SCM$_{b}$ | **0.864±0.091** | 0.876±0.166 | 0.846±0.121 | 0.771±0.205 | 1.0 ± 0.0 |
| | | | | | | SCM$_{cv}$ | 0.850±0.088 | 0.860±0.178 | 0.832±0.108 | 0.750±0.204 | 1.2 ± 0.4 |
| | penicillin | 172 | 113 | 59 | | L1-logistic | **0.994±0.012** | **0.996±0.014** | **0.992±0.024** | **0.995±0.010** | 233.3 ± 219.9 |
| | | | | | | L2-logistic | 0.976±0.033 | 0.988±0.027 | 0.953±0.062 | 0.983±0.024 | all* |
| | | | | | | Majority | 0.694±0.068 | **1.000±0.000** | 0.000±0.000 | 0.818±0.048 | – |
| | | | | | | Naive Bayes | 0.835±0.048 | 0.774±0.066 | 0.970±0.049 | 0.865±0.045 | all |
| | | | | | | Poly-SVM | 0.941±0.046 | 0.964±0.044 | 0.889±0.094 | 0.958±0.031 | all |
| | | | | | | RBF-SVM | 0.950±0.042 | 0.975±0.035 | 0.889±0.094 | 0.964±0.029 | all |
| | | | | | | Random Forests | **0.994±0.012** | **0.996±0.014** | **0.986±0.045** | **0.996±0.009** | 436.0 ± 730.2 |
| | | | | | | CART$_{b}$ | 0.982±0.025 | **0.996±0.014** | 0.949±0.073 | **0.988±0.018** | 1.0 ± 0.0 |
| | | | | | | CART$_{cv}$ | 0.982±0.025 | **0.996±0.014** | 0.949±0.073 | **0.988±0.018** | 1.0 ± 0.0 |
| | | | | | | SCM$_{b}$ | 0.982±0.025 | **0.996±0.014** | 0.949±0.073 | **0.988±0.018** | 1.0 ± 0.0 |
| | | | | | | SCM$_{cv}$ | 0.979±0.020 | 0.983±0.021 | 0.969±0.053 | 0.985±0.014 | 1.0 ± 0.0 |

Continued on next page

**Table S1.** (Continued)

| Species | Antibiotic | Genomes | Resistant | Susceptible | $k$-mers (millions) | Method | Accuracy | Sensitivity | Specificity | F1 score | Complexity |
|---|---|---|---|---|---|---|---|---|---|---|---|
| | tetracycline | 393 | 284 | 109 | 6.2 | L1-logistic | 0.956 ± 0.020 | 0.976 ± 0.015 | 0.909 ± 0.042 | 0.969 ± 0.015 | 9330.3 ± 9714.3 |
| | | | | | | L2-logistic | 0.956 ± 0.022 | 0.978 ± 0.012 | 0.902 ± 0.050 | 0.969 ± 0.016 | all* |
| | | | | | | Majority | 0.714 ± 0.050 | **1.000 ± 0.000** | 0.000 ± 0.000 | 0.832 ± 0.035 | – |
| | | | | | | Naive Bayes | 0.869 ± 0.057 | 0.887 ± 0.067 | 0.822 ± 0.057 | 0.904 ± 0.047 | all |
| | | | | | | Poly-SVM | 0.949 ± 0.018 | 0.980 ± 0.015 | 0.874 ± 0.047 | 0.964 ± 0.013 | all |
| | | | | | | RBF-SVM | 0.942 ± 0.017 | 0.973 ± 0.012 | 0.869 ± 0.050 | 0.960 ± 0.013 | all |
| | | | | | | Random Forests | 0.959 ± 0.025 | 0.985 ± 0.014 | 0.894 ± 0.057 | **0.971 ± 0.017** | 493.3 ± 840.0 |
| | | | | | | CART$_b$ | **0.964 ± 0.019** | 0.985 ± 0.014 | 0.911 ± 0.042 | **0.975 ± 0.013** | 1.0 ± 0.0 |
| | | | | | | CART$_{cv}$ | **0.965 ± 0.019** | 0.977 ± 0.020 | **0.937 ± 0.037** | **0.976 ± 0.014** | 2.6 ± 1.6 |
| | | | | | | SCM$_b$ | **0.964 ± 0.019** | 0.985 ± 0.014 | 0.911 ± 0.042 | **0.975 ± 0.013** | 1.0 ± 0.0 |
| | | | | | | SCM$_{cv}$ | **0.971 ± 0.016** | 0.983 ± 0.016 | **0.937 ± 0.037** | **0.979 ± 0.012** | 2.4 ± 0.8 |
| | trimethoprim/sulfamethoxazole | 2826 | 2187 | 639 | 24.2 | L1-logistic | 0.928 ± 0.011 | 0.942 ± 0.014 | **0.880 ± 0.024** | 0.953 ± 0.008 | 7172.4 ± 6532.7 |
| | | | | | | L2-logistic | 0.926 ± 0.010 | 0.943 ± 0.019 | 0.867 ± 0.030 | 0.952 ± 0.008 | all* |
| | | | | | | Majority | 0.778 ± 0.015 | **1.000 ± 0.000** | 0.000 ± 0.000 | 0.875 ± 0.009 | – |
| | | | | | | Naive Bayes | 0.854 ± 0.019 | 0.852 ± 0.026 | 0.858 ± 0.032 | 0.900 ± 0.015 | all |
| | | | | | | Poly-SVM | **0.935 ± 0.006** | 0.969 ± 0.008 | 0.815 ± 0.041 | **0.958 ± 0.004** | all |
| | | | | | | RBF-SVM | **0.934 ± 0.006** | 0.970 ± 0.009 | 0.811 ± 0.042 | **0.958 ± 0.004** | all |
| | | | | | | Random Forests | **0.943 ± 0.008** | 0.984 ± 0.005 | 0.801 ± 0.025 | **0.964 ± 0.005** | 19693.3 ± 25788.2 |
| | | | | | | CART$_b$ | **0.939 ± 0.010** | 0.973 ± 0.008 | 0.819 ± 0.029 | **0.961 ± 0.007** | 5.4 ± 1.0 |
| | | | | | | CART$_{cv}$ | **0.938 ± 0.010** | 0.971 ± 0.009 | 0.822 ± 0.030 | **0.960 ± 0.007** | 8.6 ± 4.4 |
| | | | | | | SCM$_b$ | **0.938 ± 0.011** | 0.981 ± 0.013 | 0.789 ± 0.017 | **0.961 ± 0.007** | 3.1 ± 0.3 |
| | | | | | | SCM$_{cv}$ | **0.937 ± 0.008** | 0.983 ± 0.006 | 0.778 ± 0.024 | **0.961 ± 0.005** | 4.0 ± 2.8 |

**Table S2.** Extended benchmark. Comparison to state-of-the-art classifiers in terms of accuracy and model complexity. For each dataset the accuracy is shown, along with the number of $k$-mers used by the model (in parentheses). Results are shown for Set Covering Machines (SCM), Classification trees (CART), Random Forests[13] with $\chi^2$ feature selection, Logistic regression with L1 and L2 regularization and $\chi^2$ feature selection (L1-logistic, L2-logistic), Polynomial kernel and RBF kernel Support Vector Machines (Poly-SVM, RBF-SVM), Naive Bayes, and a baseline predictor that predicts the most abundant class in the data (Majority). Accuracies within 1% of the maximum value are shown in bold. Results are averaged over ten repetitions of the experiment.

| Dataset | $SCM_b$ | $CART_b$ | Random forests*, † | L1-logistic*, † | L2-logistic*, † | Poly-SVM† | RBF-SVM† | Naive Bayes | Majority |
|---|---|---|---|---|---|---|---|---|---|
| *A. baumannii* | 0.849 (2.7) | 0.864 (3.4) | **0.892** (6314.6) | 0.880 (3980.5) | **0.885** (1e6) | **0.886** (all) | 0.880 (all) | 0.822 (all) | 0.644 |
| *E. coli* | **0.818** (4.6) | **0.808** (7.0) | **0.812** (39289.6) | 0.792 (3727.2) | 0.789 (1e6) | 0.779 (all) | 0.776 (all) | 0.634 (all) | 0.697 |
| *E. faecium* | **1.000** (1.0) | **1.000** (1.0) | **1.000** (202.6) | **1.000** (142.0) | **1.000** (1e6) | **0.996** (all) | **0.992** (all) | 0.808 (all) | 0.588 |
| *K. pneumoniae* | **0.950** (3.9) | **0.949** (4.3) | **0.956** (42856.8) | **0.952** (7607.4) | **0.948** (1e6) | 0.943 (all) | 0.943 (all) | 0.760 (all) | 0.571 |
| *M. tuberculosis* | **0.963** (4.5) | **0.962** (4.7) | **0.962** (78761.3) | **0.962** (2242.2) | 0.941 (1e6) | 0.934 (all) | 0.930 (all) | 0.789 (all) | 0.658 |
| *N. gonorrhoeae* | **0.935** (3.0) | **0.936** (3.3) | 0.895 (4571.7) | **0.942** (6095.6) | 0.915 (1e6) | 0.906 (all) | 0.905 (all) | 0.736 (all) | 0.529 |
| *P. aeruginosa* | **0.939** (1.2) | **0.942** (1.1) | 0.874 (21600.5) | **0.937** (87.8) | 0.828 (1e6) | 0.773 (all) | 0.762 (all) | 0.768 (all) | 0.588 |
| *P. difficile* | **0.982** (1.0) | **0.982** (1.0) | 0.949 (662.2) | 0.957 (121.8) | 0.936 (1e6) | 0.949 (all) | 0.951 (all) | 0.887 (all) | 0.599 |
| *S. aureus* | **0.987** (1.0) | **0.987** (1.0) | **0.987** (408.8) | **0.988** (230.6) | **0.987** (1e6) | **0.987** (all) | **0.987** (all) | 0.868 (all) | 0.544 |
| *S. enterica* | 0.913 (1.0) | 0.913 (1.0) | **0.926** (2354.8) | **0.925** (991.2) | **0.929** (1e6) | **0.920** (all) | **0.928** (all) | 0.759 (all) | 0.709 |
| *S. haemolyticus* | **0.925** (1.0) | **0.925** (1.0) | 0.846 (2820.0) | **0.925** (279.1) | 0.838 (1e6) | 0.829 (all) | 0.846 (all) | 0.758 (all) | 0.629 |
| *S. pneumoniae* | **0.960** (1.0) | **0.960** (1.0) | **0.957** (92.2) | 0.948 (1391.5) | 0.949 (1e6) | 0.946 (all) | 0.944 (all) | 0.910 (all) | 0.654 |

* For scalability reasons, these algorithms were trained using feature selection to select the one million $k$-mers that were most associated with the phenotypes; all other $k$-mers were discarded (see *Methods*).

† The implementations available in Scikit-Learn 0.18.2 were used. When applicable, the kernel matrices were precomputed using custom code based on the $k$-mer matrices.

**Table S3.** Sample compression bound values for the $SCM_b$ models on each benchmark dataset. We report other quantities that are relevant to the interpretation of the bound, such as the accuracy and complexity (number of rules) of the models and the number of examples in the datasets. We also show the number of $k$-mers in each dataset, since, surprisingly, this value does not take part in the calculation of the bound. The reader is encouraged to observe the expression of the bound (Equation (2)) in parallel to understand how each quantity affects the bound. Note that values that varied over ten repetitions of the experiment are shown as mean $\pm$ standard deviation.

| Dataset | Bound | Accuracy | Complexity | Examples | $k$-mers (millions) |
|---|---|---|---|---|---|
| *A. baumannii* | $0.427 \pm 0.022$ | $0.849 \pm 0.031$ | $2.7 \pm 0.5$ | 499 | 42.4 |
| *E. coli* | $0.473 \pm 0.007$ | $0.818 \pm 0.019$ | $4.6 \pm 1.1$ | 1524 | 48.5 |
| *E. faecium* | $0.236 \pm 0.000$ | $1.000 \pm 0.000$ | $1.0 \pm 0.0$ | 134 | 10.3 |
| *K. pneumoniae* | $0.220 \pm 0.003$ | $0.950 \pm 0.007$ | $3.9 \pm 0.7$ | 2107 | 70.3 |
| *M. tuberculosis* | $0.160 \pm 0.003$ | $0.963 \pm 0.005$ | $4.5 \pm 0.5$ | 5022 | 11.7 |
| *N. gonorrhoeae* | $0.331 \pm 0.018$ | $0.935 \pm 0.030$ | $3.0 \pm 0.0$ | 392 | 4.8 |
| *P. aeruginosa* | $0.284 \pm 0.015$ | $0.939 \pm 0.023$ | $1.2 \pm 0.4$ | 491 | 43.0 |
| *P. difficile* | $0.164 \pm 0.009$ | $0.982 \pm 0.009$ | $1.0 \pm 0.0$ | 462 | 19.8 |
| *S. aureus* | $0.074 \pm 0.005$ | $0.987 \pm 0.005$ | $1.0 \pm 0.0$ | 1593 | 13.3 |
| *S. enterica* | $0.369 \pm 0.012$ | $0.913 \pm 0.024$ | $1.0 \pm 0.0$ | 347 | 6.9 |
| *S. haemolyticus* | $0.431 \pm 0.027$ | $0.925 \pm 0.047$ | $1.0 \pm 0.0$ | 120 | 5.3 |
| *S. pneumoniae* | $0.233 \pm 0.013$ | $0.960 \pm 0.018$ | $1.0 \pm 0.0$ | 409 | 6.4 |

**Table S4.** Sample compression bound values for the $CART_b$ models on each benchmark dataset. We report other quantities that are relevant to the interpretation of the bound, such as the accuracy and complexity (number of rules) of the models and the number of examples in the datasets. We also show the number of $k$-mers in each dataset, since, surprisingly, this value does not take part in the calculation of the bound. The reader is encouraged to observe the expression of the bound (Equation (3)) in parallel to understand how each quantity affects the bound. Note that values that varied over ten repetitions of the experiment are shown as mean ± standard deviation.

| Dataset | Bound | Accuracy | Complexity | Examples | $k$-mers (millions) |
|---|---|---|---|---|---|
| *A. baumannii* | 0.423 ± 0.010 | 0.864 ± 0.042 | 3.4 ± 0.7 | 499 | 42.4 |
| *E. coli* | 0.464 ± 0.008 | 0.808 ± 0.021 | 7.0 ± 0.7 | 1524 | 48.5 |
| *E. faecium* | 0.249 ± 0.000 | 1.000 ± 0.000 | 1.0 ± 0.0 | 134 | 10.3 |
| *K. pneumoniae* | 0.223 ± 0.004 | 0.949 ± 0.007 | 4.3 ± 1.2 | 2107 | 70.3 |
| *M. tuberculosis* | 0.162 ± 0.003 | 0.962 ± 0.004 | 4.7 ± 1.2 | 5022 | 11.7 |
| *N. gonorrhoeae* | 0.341 ± 0.017 | 0.936 ± 0.039 | 3.3 ± 0.5 | 392 | 4.8 |
| *P. aeruginosa* | 0.288 ± 0.015 | 0.942 ± 0.028 | 1.1 ± 0.3 | 491 | 43.0 |
| *P. difficile* | 0.168 ± 0.009 | 0.982 ± 0.009 | 1.0 ± 0.0 | 462 | 19.8 |
| *S. aureus* | 0.076 ± 0.005 | 0.987 ± 0.005 | 1.0 ± 0.0 | 1593 | 13.3 |
| *S. enterica* | 0.373 ± 0.012 | 0.913 ± 0.024 | 1.0 ± 0.0 | 347 | 6.9 |
| *S. haemolyticus* | 0.442 ± 0.027 | 0.925 ± 0.047 | 1.0 ± 0.0 | 120 | 5.3 |
| *S. pneumoniae* | 0.237 ± 0.013 | 0.960 ± 0.018 | 1.0 ± 0.0 | 409 | 6.4 |
